# In vivo single cell transcriptomics reveals *Klebsiella pneumoniae* rewiring of lung macrophages to promote infection

**DOI:** 10.1101/2021.10.18.464784

**Authors:** Amy Dumigan, Oisin Cappa, Brenda Morris, Joana Sá Pessoa, Ricardo Calderon-Gonzalez, Grant Mills, Rebecca Lancaster, David Simpson, Adrien Kissenpfennig, Jose A. Bengoechea

## Abstract

The strategies deployed by antibiotic resistant bacteria to counteract host defences are poorly understood. Here, we elucidate a novel host-pathogen interaction that results in the control of lung macrophage polarisation by the human pathogen *Klebsiella pneumoniae*. We identify interstitial macrophages (IMs) as the main population of lung macrophages associated with *Klebsiella*. Single cell transcriptomics and trajectory analysis of cells uncover that type I IFN and IL10 signalling, and macrophage polarization are characteristic of infected IMs, whereas Toll-like receptor (TLR) and Nod-like receptor signalling are features of infected alveolar macrophages. *Klebsiella*-induced macrophage polarization is a singular M2-type we termed M(Kp). To rewire macrophages towards M(Kp), *K. pneumoniae* hijacks a hitherto unknown TLR-type I IFN-IL10-STAT6 innate axis. Absence of STAT6 limits the intracellular survival of *Klebsiella* whereas the inhibition of STAT6 facilitates the clearance of the pathogen in vivo. Glycolysis characterises M(Kp) metabolism, and inhibition of glycolysis results in clearance of intracellular *Klebsiella*. We demonstrate the capsule polysaccharide is the *Klebsiella* factor governing M(Kp). *Klebsiella* also skews human macrophage polarization towards M(Kp) in a type I IFN-IL10-STAT6-dependent manner. Altogether, our work demonstrates that *Klebsiella* induction of M(Kp) represents a hitherto unknown strategy to overcome host restriction during pneumonia.

## Introduction

Antibiotic resistance is a pandemic claiming more than 750,000 deaths per year. *Klebsiella pneumoniae* exemplifies the threat of this pandemic by the increasing number of strains resistant to fluoroquinolones, third-generation cephalosporins, aminoglycosides, and even carbapenems (1). These infections are associated with high mortality rates and prolonged hospitalization (2). Not surprisingly, the World Health Organization includes *K. pneumoniae* in the “critical” group of pathogens for which new therapeutics are urgently needed.

Less obvious but critical for pathogenesis are *K. pneumoniae* adaptations to the human immune system allowing the pathogen to flourish in human tissues such as the airways. This is an aspect often overlooked because *K. pneumoniae* is not considered a pathogen able to manipulate the host cells because it does not encode type III or IV secretion systems known to deliver effectors into immune cells, or any of the toxins affecting cell biology. Of particular interest is to understand whether *K. pneumoniae* deploys any strategy to manipulate macrophage function. These cells are crucial in host defence against infection by eliminating the invading pathogen via phagocytosis and the subsequent degradation in a phagolysosomal compartment, and by producing cytokines and chemokines following recognition of the pathogen to orchestrate the activation of other immune cells.

Macrophages show a remarkable plasticity allowing them to adapt to different microenvironments. Signals such as tissue damage, presence of a pathogen, and the presence of cytokines and other immune cells dictate different polarization states of macrophages. Depending on the stimuli, macrophages broadly differentiate into type M1, pro-inflammatory showing potent microbicidal activity, M2 with immunomodulatory role to limit tissue damage and to control inflammation, M3 or “switch” state (3), and M4 which is mediated by CXCL4 and observed in differentiated atherosclerotic plaque-associated macrophages (4). Different subsets of M2 macrophages have been identified, from M2a to M2d; all of them have in common the high-level expression of IL10 compared to M1 macrophages (5). Interestingly, increasing the number of M2 macrophages in the lungs as result of alcohol abuse or trauma is associated with increased susceptibility to *K. pneumoniae* infections (6–8). On the contrary, there is an improvement in bacterial clearance when the M2 macrophage population is eliminated (6–8), or after skewing macrophages towards an M1 state (9). These clinical observations suggest a role for macrophage polarization in *K. pneumoniae* infection biology, although this has not been investigated yet. The manipulation of macrophage polarization is emerging as an important virulence strategy of intracellular pathogens. However, and contrary to the conventional wisdom, there is no clear pattern of macrophage polarization induced by intracellular pathogens (10, 11). Furthermore, there are few cases in which this has been studied *in vivo*. Therefore, it remains an open question whether the polarization state induced by a pathogen is a host protective mechanism or represents a virulence strategy.

This work was designed to provide a comprehensive understanding of the *K. pneumoniae*-macrophage interface *in vivo*. We identify interstitial macrophages (IMs) as the main population of lung macrophages associated with *K. pneumoniae*. Single cell transcriptomics uncover the programme induced by *K. pneumoniae* in infected and bystander alveolar macrophages (AMs) and IMs. Pathway analysis reveal a network involved in macrophage polarization, and mechanistic studies demonstrate that *K. pneumoniae* exploits the immune effectors IL10 and type I IFNs to trigger a singular macrophage polarization governed by the signal transducer and activator of transcription (STAT6) following the activation of TLR signalling. Inhibition of this pathway results in clearance of *K. pneumoniae*, illustrating the importance of macrophage polarization in *K. pneumoniae* infection biology. Altogether, our work describes a new polarization state induced by a human pathogen to overcome host restriction during pneumonia.

## Results

### K. pneumoniae *is associated with interstitial macrophages and alveolar macrophages*

In the mouse lungs, and similarly to the human lungs, two main populations of macrophages have been identified: AMs, with an embryonic origin, and IMs, originating mainly from hematopoietic stem cells (12–14). A population of circulating monocytes (MNs) can be also found in the lungs (12–14). To determine to which macrophage subpopulations *K. pneumoniae* associates in the lungs of infected mice, C57BL/6 mice were infected intranasally with the clinical isolate *K. pneumoniae* CIP52.145 (hereafter Kp52145) (15). This strain clusters within the KpI group that includes the strains associated with human infections (16, 17). Moreover, Kp52145 encodes all the loci found in those strains associated with invasive community acquired infections (16, 17). To facilitate the detection of *K. pneumoniae* in vivo, Kp52145 was tagged with mCherry. 24 h and 48 h post-infection, lungs were processed and stained for cytometric analysis of MNs (Ly6C^+^CD11b^+^CD11c^-^), IMs (Ly6C^+^CD11b^+^CD11c^-^SiglecF^-^), and AMs (Ly6C^+^CD11b^-^CD11c^+^Siglec F^+^). The gating strategy is shown in Supplementary data figure 1. The number of MNs, AMs and IMs did not change over time in PBS-treated mice (Fig 1A). The number of MNs was higher in Kp52145-infected mice at 48 h post infection than at 24 h (Fig 1A). The number of AMs was not significantly different between non-infected and infected mice (Fig 1A) which is consistent with previous findings (18). In contrast, the number of IMs was significantly higher in infected mice than in PBS control animals (Fig 1A). However, the number of IMs was not significantly different 48 and 24 h post infection (Fig 1A). Flow cytometric detection of mCherry-tagged Kp52145 showed that at 24 h post infection, 31% of IMs were associated with Kp52145 whereas 1% of MNs and 5% AMs were positive (Fig 1B). At 48 h post infection, MNs remained negative for *Klebsiella* whereas there was a significant increase in the percentage of AMs associated with Kp52145 reaching 15% (Fig 1B). The percentage of IMs associated with Kp52145 at 48 h was not significantly different to that at 24 h post infection (Fig 1B). Collectively, these results demonstrate that in vivo *K. pneumoniae* is associated with two ontogenetically distinct macrophage lineages, namely IMs and AMs.

**Figure 1.**
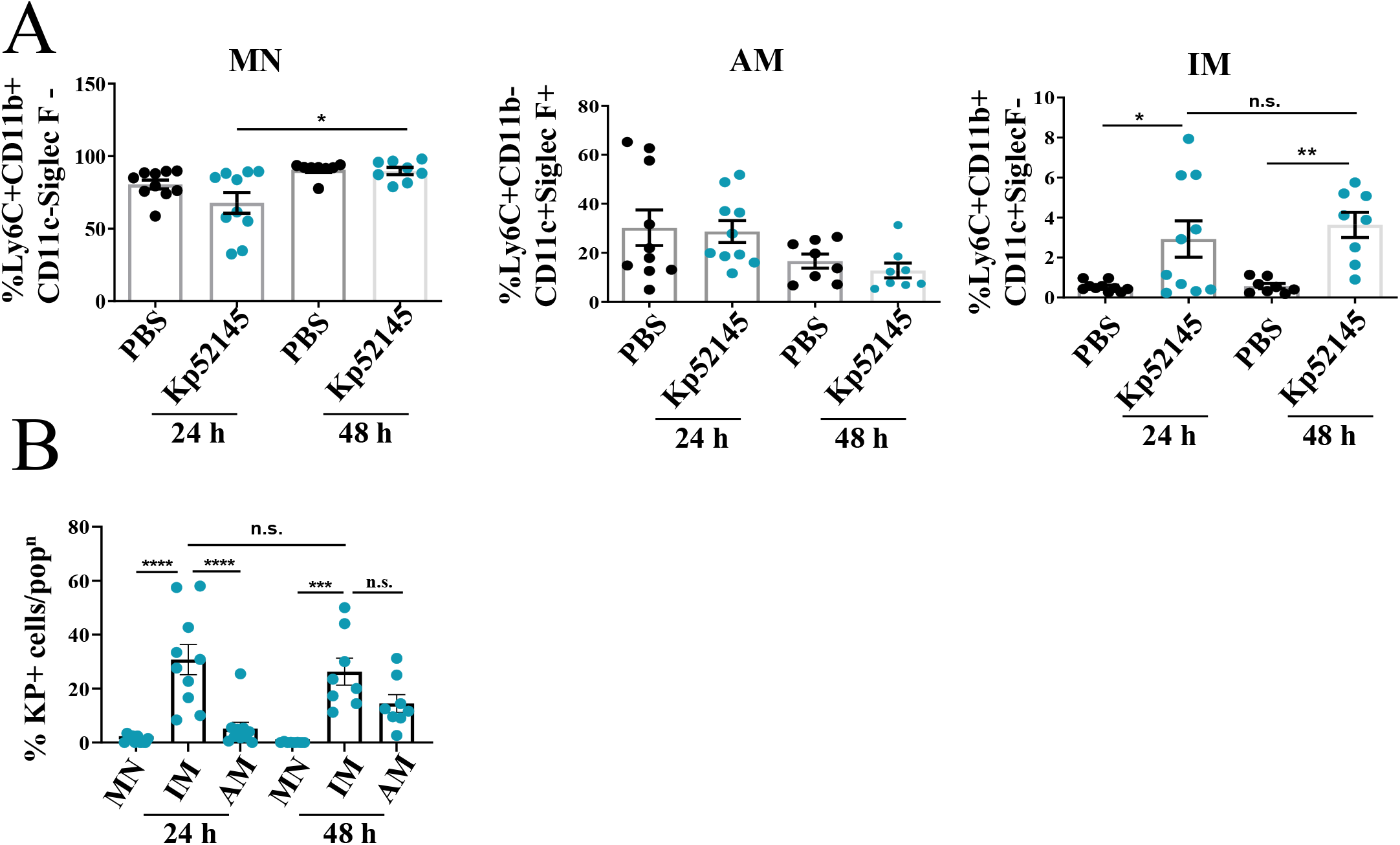
*K. pneumoniae* is associated with interstitial macrophages and alveolar macrophages in vivo. 10-12 week old age and sex matched C57BL/6 mice were infected intranasally with mCherry tagged Kp52145. After 24 h or 48 h post-infection (n=10/condition) lungs were harvested and processed for flow cytometric analysis to assess macrophages populations. A. % Ly6C+CD11b-CD11c-SiglecF-monocyte (MNs), % Ly6C+CD11b-CD11c+SiglecF+ alveolar macrophage (AMs), and % Ly6C+CD11b+CD11c+SiglecF-interstitial macrophage (IMs) in Kp52145–infected animals compared to PBS controls. B. Percentage of MN, AM, and IM associated with Kp52145 from infected individual mice. In all panels, values are presented as the mean ± SEM of two independent experiments. ****P ≤ 0.0001; ***P ≤ 0.001; ** P ≤ 0.01; ns, P > 0.05 for the indicated comparisons determined using One-way ANOVA.

### *Single cell RNA-seq reveals distinct transcriptomes in IM and AM populations during* K. pneumoniae *infection* in vivo

We sought to characterize the transcriptomes of infected IMs and AMs in comparison to those of bystander IMs and AMs from infected mice, and to those of IMs and AMs from PBS-mock infected mice. Mice were infected with mCherry-tagged Kp52145, and the IMs and AMs populations with and without associated bacteria were separated by FACS sorting (Fig 2A). The gating strategy for cell sorting is outlined in Supplementary Figure 1. Single cell mRNA sequencing (scRNA-seq) technology using the 10 x Genomics platform was utilised to determine the transcriptome of the different populations of macrophages. The resulting dataset was curated using the Immgen20 open-source reference database (19) to remove any non-macrophage cells from the data set, resulting in a total of 7,462 macrophages. 512 and 1113 AM and IMs, respectively from PBS-treated mice, 3080 AMs from infected mice, 2281 with associated Kp52145 and 758 bystander AMs, and 2797 IMs from infected mice, 2273 with associated *Klebsiella* and 524 bystander cells. After normalisation of the samples, uniform manifold approximation and projection (UMAP) dimensionality reduction analysis of the combined samples revealed two clusters. One of the clusters comprised the IMs, characterised by the expression of the IM marker *cx3cr1,* whereas the other comprised the AMs, characterised by the expression of the marker *siglecF* (Fig 2B). In the case of the IM cluster, it was possible to distinguish the cluster of cells from non-infected mice from those from infected mice which were further separated between bystander cells and those with associated bacteria (Fig 2C). In contrast, there was no clear separation between the different populations of macrophages in the AM cluster.

**Figure 2.**
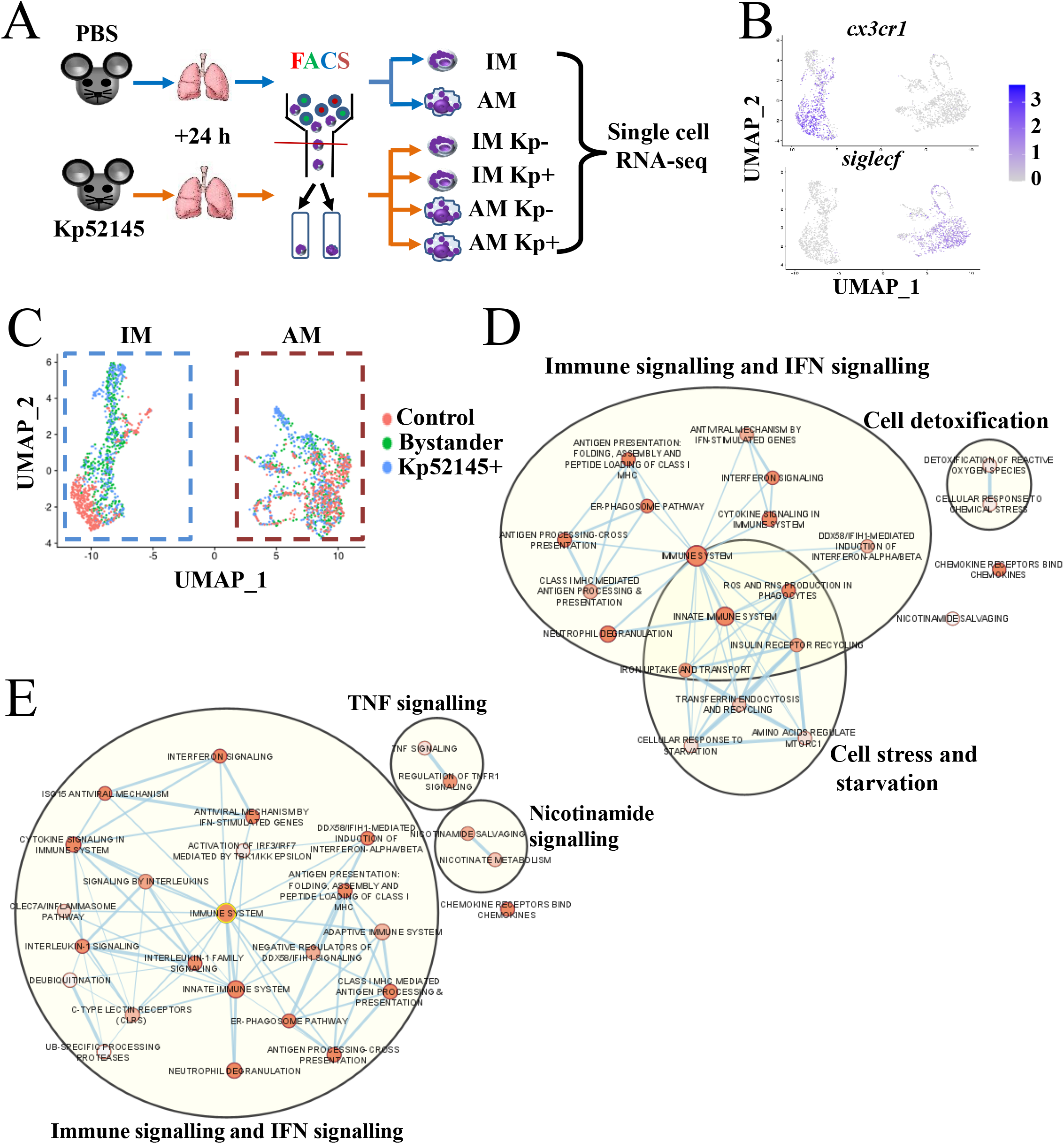
Analysis of *K. pneumoniae*-induced transcriptome in IMs. A. Diagram of the experimental approach to generate the different IMs and AMs samples for single-cell RNA sequencing (scRNAseq). C57BL/6 mice (n=17/group) were infected intranasally with mCherry tagged Kp52145, after 24 h, lungs were excised, and processed for cell sorting. From pooled samples AM and IM populations were sorted from PBS controls and infected mice. In the latter group, cells were sorted to separate bystander cells and cells with associated bacteria. The viability of each of the samples was determined to be higher than 95% before carrying out 10x genomics single cell RNA sequencing. B. Marker gene detection and differential expression testing was performed in Seurat using the MAST package. Higher resolution clustering using uniform manifold approximation and projection (UMAP) dimensionality reduction analysis showing selected genes, *cx3cr1,* IM marker, and *SiglecF,* AM marker. C. UMAP of clustering within cells from PBS mock-infected mice (control), bystander and Kp52145-associated IMs and AM populations. D. Network enrichment mapping generated from significantly upregulated genes of IMs with associated bacteria. Analysis was performed using the g:SCS method for multiple testing correction (gProflier), the Reactome database as a data source, and the default settings for the other parameters in gProflier. Results were exported to Cytoscape and visualized using the AutoAnnotate plug. E. Network enrichment mapping generated from significantly upregulated genes of bystander IMs. Analysis was performed using the g:SCS method for multiple testing correction (gProflier), the Reactome database as a data source, and the default settings for the other parameters in gProflier. Results were exported to Cytoscape and visualized using the AutoAnnoate application

Differential gene expression analysis revealed that 1,083 genes were differentially expressed in IMs from infected mice versus PBS-mock infected mice. Of those, 393 genes were common between bystander and infected IMs, whereas 126 and 171 were only found in bystander cells and infected IMs, respectively. We compared the transcriptome of the different populations of IMs to identify signatures of infected and bystander cells. 890 and 979 genes were differentially expressed in bystander and infected IMs versus PBS-mock infected IMs, respectively (Supplementary Table 1). Of them, 518 and 564 genes were upregulated whereas 372 and 415 were down regulated in bystander and infected IMs versus PBS-mock infected IMs, respectively (Supplementary Table 1). To acquire insights into the biological processes of significance that characterize infected IMs and bystander cells, we performed gene set enrichment analysis (gProfiler) and then constructed network enrichment maps. When considering the downregulated genes, bystander and Kp52145-infected IMs shared an enrichment of networks related to translation (Supplementary Fig 2A and B). Pathways related to TGFβ signalling and Erbb4-Notch signalling were specific of bystander and Kp52145-infected IMs, respectively (Supplementary Fig 2A and B). In the case of the upregulated genes, there was an enrichment of pathways related to immune signalling in IMs from infected mice (Fig 2D and E). It is notable the over representation of gene networks related to interferon signalling in bystander and infected IMs (Fig 2D and 2E). This finding is consistent with an enrichment of motifs for transcriptional factors of the Irf family, and STAT1 in the promoter regions of the upregulated genes of IMs from infected mice as detected interrogating the TRANSFAC database (20). Only in Kp52145-infected IMs, we found gene networks involved in response to oxidative stress, starvation, iron uptake, and macrophage polarization, (*nos2, arg1*, *mrc1/cd206)* (Fig 2D). Interestingly, TRANSFAC-based analysis revealed an enrichment of the motif recognized by the transcriptional factor GKLF/Klf4 only in infected IMs. This transcriptional factor regulates M2 macrophage polarization (21). In contrast, networks found only in bystander IMs included those related to TNF signalling, inflammasome activation and IL1 signalling, and C-type lectin receptors (Fig 2E). Altogether, these results uncover that upregulation of IFN signalling and downregulation of translation are features of IMs following *K. pneumoniae* infection. The signature of infected IMs is the activation of networks connected to cellular stress and macrophage polarization whereas networks connected to antimicrobial defence and sensing of infections are specific of bystander IMs.

Within the AM population, 218 genes were differentially expressed in Kp52145-associated AMs versus PBS-mock infected whereas only 63 genes were differentially expressed in bystander AMs. (Supplementary Table 1). No networks were enriched within the downregulated genes of Kp52145-infected AMs whereas supplementary Figure 3 shows the network enrichment maps corresponding to the upregulated genes. Four clusters were identified, receptor signalling cluster having the most nodes. Gene networks within this cluster are related to Toll-like receptor (TLR) and Nod-like receptor (NLR) signalling. TRANSFAC-based analysis showed that the motif recognized by the NF-κB transcriptional factor is enriched in the promoter region of the genes within this cluster. Connected to this cluster are the clusters of TNF signalling, and antigen presentation. Networks related to immune signalling include cytokines and chemokines, *il1b*, *tnfa, cxclc2, cxcl3*, and calcium-dependent inflammatory proteins, *s100A9*, *s100A8*. No pathways were enriched in bystander AMs. Collectively, these results demonstrate a reduced activation of AMs following infection compared to IMs. The transcriptional pattern of infected AMs is related to TLR and NLR signalling-governed inflammation in an NF-κB-dependent manner.

To establish whether it is possible to construct a trajectory from non-infected to infected cells, we utilised Monocle analysis to determine the temporal pattern of gene expression over pseudotime. Whereas no distinct trajectory was observed in AMs (Supplementary Fig 4), Monocle analysis revealed a clear trajectory in IMs from non-infected cells to bystander cells to Kp52145-infected cells (Fig 3A). Seven modules of genes showing similar pattern of expression were identified (Supplementary Table 2). Heat map analysis revealed that genes included in modules 3, 4 and 6 were upregulated in IMs from infected mice, whereas the expression of genes in modules 3 and 4 were higher in Kp52145-infected IMs than in bystander cells (Fig 3B). Genes in module 6 were upregulated in bystander cells (Fig 3B). In contrast, genes in modules 1 and 7 were upregulated in IMs from non-infected mice (Fig 3B). Enrichment map analysis of modules 1 and 7 using gProfiler revealed one cluster related to regulation of translation (Supplementary Fig 5). Pathway analysis of modules 3 and 4, characteristic of *K. pneumoniae*-infected IMs, revealed an enrichment in pathways related to immune signalling, particularly type I IFN stimulated genes (ISGs) *(isg15, cxc10, ifit1, usp18*, *irgm1, irg1*) and IL10 signalling (*ptgs2, csf1, ptafr*), and IL4 signalling and macrophage polarization (*nos2, arg1, lcn2, socs1, socs3*) (Fig 3C). STRING analysis seeking functional interactions between the 234 genes revealed two clusters (Fig 3D). One cluster includes 39 genes associated with IFN signalling, and the other one encompasses 74 genes related to cytokine signalling and macrophage polarization (Fig 3D). TRANSFAC analysis showed that the regulatory regions of genes within modules 3 and 4 are characterized by motifs for transcriptional factors of the Irf family, and for the p65 subunit of NF-κB. Module 6, characteristic of bystander cells, is enriched with genes related to immune signalling, and IFN signalling (Fig 3E). This module is characterized by the presence of binding motifs for Irf5 and Irf8 in the regulatory regions of the genes. Figure 3F shows the changes over pseudotime of selected top expressed genes within the modules 3, 4 and 6. Data demonstrates the increase in transcription of ISGs IFN (*isg15, cxcl10, irg1, ifi203, mndA, ifi205*), IL10 signalling (*ptgs2, ptafr*), IL4 signalling and macrophage polarization (*nos2, arg1, lcn2, socs1, socs3*) from non-infected to bystander to infected cells. In contrast, there is a decrease in transcription from non-infected cells to bystander to infected cells of genes related to translation (*rnas6, rpsA, rps21*) and some others related to migration of immune cells and activation (*pparg*, *ifngr1, cytip, lsp1*). Altogether, this analysis revealed that IFN and IL10 signalling governed by Irf and NF-κB transcriptional factors is characteristic of *K. pneumoniae* infected IMs whereas Irf-controlled IFN signalling is characteristic of bystander IMs. Furthermore, our data suggested that *K. pneumoniae* infection skews macrophage polarization.

**Figure 3.**
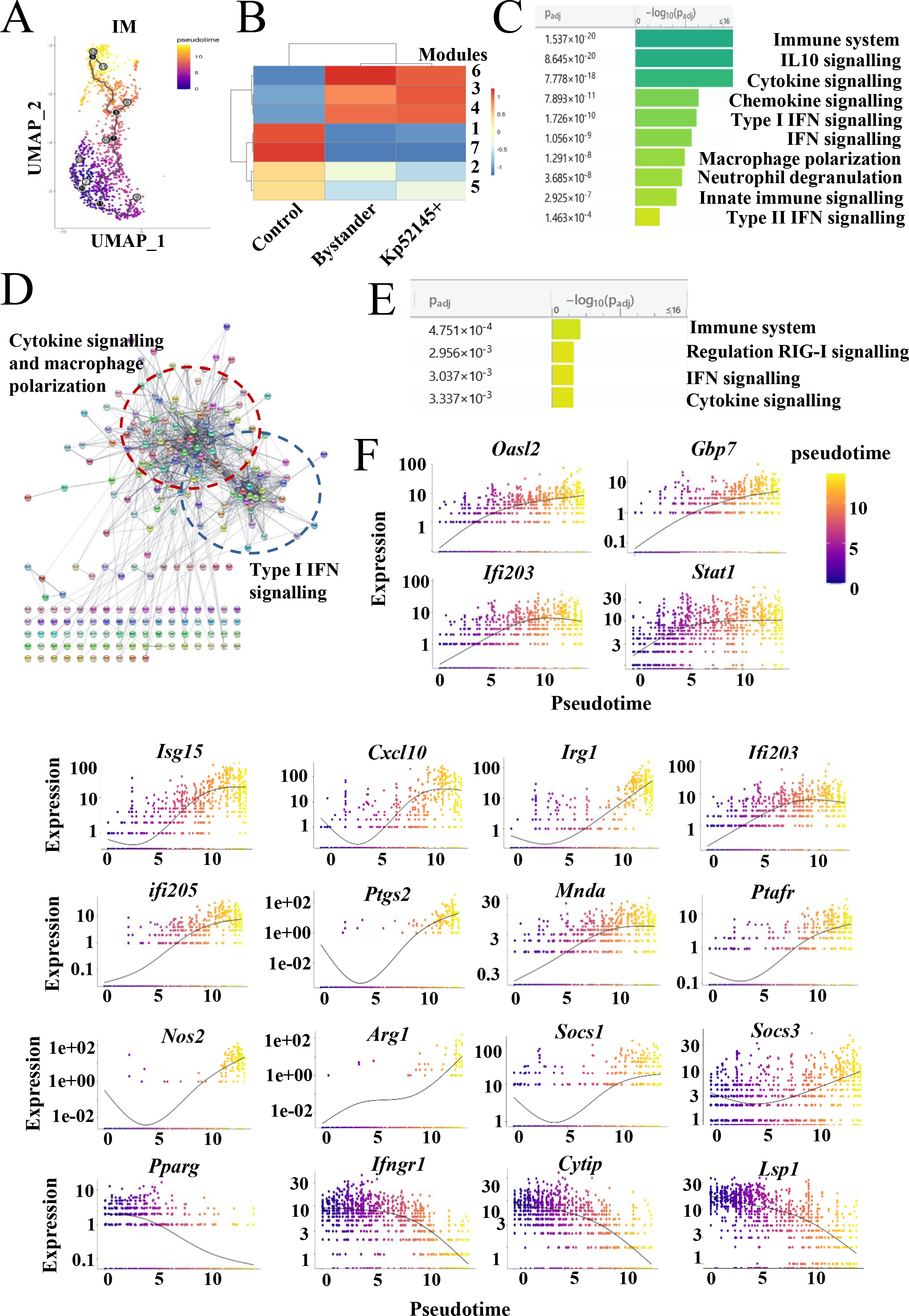
Single-cell trajectory analysis of IMs from non-infected to infected cells. A. Monocle analysis to determine the temporal pattern of gene expression over pseudotime in bystander and Kp52145-associated IMs from infected animals compared to PBS controls. Monocle analysis revealed 7 modules of genes showing similar pattern of expression. B. Heat map showing relative expression of the 7 modules found in IMs. C. Pathway analysis of modules 3 and 4 corresponding to Kp52145-infected IMs. Analysis was performed using the g:SCS method for multiple testing correction, the Reactome database as a data source, and the default settings for the other parameters in G:profiler. D. STRING database was used to predict protein-protein interactions using the clustering algorithm MCL with default parameters using as data source the genes within modules 3 and 4. E. Pathway analysis of module 6 corresponding to bystander IMs. Analysis was performed using the g:SCS method for multiple testing correction, the Reactome database as a data source, and the default settings for the other parameters in G:profiler. F. Changes over pseudotime of selected top expressed genes within the modules 3, 4 and 6.

### K. pneumoniae *induces a singular polarization state in interstitial and alveolar macrophages*

The fact that one of the features of Kp52145-infected macrophages was the upregulation of genes associated with macrophage polarization led us to interrogate further the scRNA-seq data set to assess the expression of genes associated with macrophage polarization. Heat map analysis revealed an upregulation of several M1 related genes in bystander and infected IMs including *il1b, il12b, cd38, cxcl10, cxcl2, and nos2* (Fig 4A). However, the expression of M2 related genes was higher than that of M1 genes (Fig 4A). M2 upregulated genes included *msr1, cxcl16, egr2, arg1, il1rn, mmp14, ccr1, ccr2, clec4b* and *parp14* (Fig 4A). Remarkably, Kp52145-induced polarization has features of several of M2 subsets. *Arg1, il1rn, il10* and *fizz1*are found in M2a cells*, il10, tnfa, il6* and *il12* are typical of M2b cells*, cd163, cd206, il10* and *arg1* are characteristic of M2c macrophages, whereas *il10* and *nos2* are found in M2d cells (5). A similar picture was observed in AMs (Supplementary Figure 6) although the number of upregulated genes was reduced compared to IMs.

**Figure 4.**
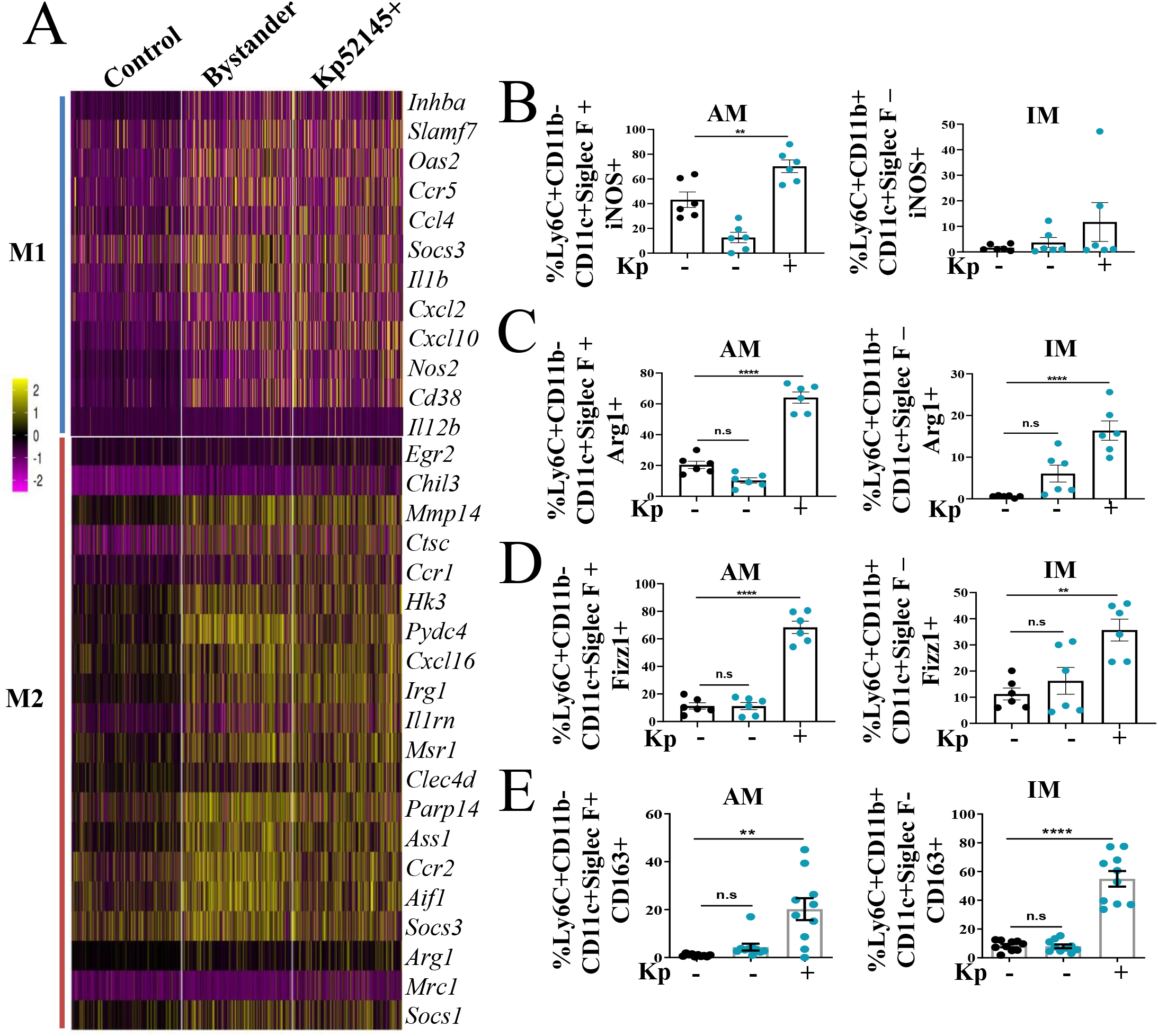
*K. pneumoniae* skews macrophage polarization towards a singular state termed M(Kp). A. Heat map presents relative expression of the indicated genes between IMs from non-infected mice (control), and bystander and Kp52145-associated IMs from infected mice. Selected genes are related to M1 and M2 macrophage polarisation. Analysis by flow cytometry of the levels of M(Kp) markers expressed by cells from PBS mock infected mice (black dots), and by cells from infected mice (blue dots) with and without associated Kp52145. B. Percentage of positive cells for iNOS. C. Percentage of positive cells for Arg1. D. Percentage of positive cells for Fizz1. E. Percentage of positive cells for CD136. Values in panel B-E are presented as the mean ± SEM whereby each dot represents an individual animal. ****P ≤ 0.0001; ***P ≤ 0.001; **P≤ 0.01; *P ≤ 0.05; ns, P > 0.05 for the indicated comparisons using one way-ANOVA with Bonferroni contrast for multiple comparisons test.

Next, we carried out flow cytometry experiments interrogating infected and bystander IMs and AMs to confirm the scRNA-seq observations. iNOS was upregulated only in Kp52145-asscociated AMs but not on the IM population (Fig 4B). The M2 markers Arg1, Fizz1, and CD163 were upregulated in Kp52145-infected IMs and AMs but not in bystander cells (Fig 4C-E), uncovering the importance of macrophage-*Klebsiella* contact for macrophage polarization.

Altogether, these results suggest that *K. pneumoniae* induces a singular polarization in IMs and AMs consistent with M2 polarization state. We term this macrophage polarization as M(Kp) because it cannot be ascribed to any of the known M2 subsets. scRNA-seq data, including pathway analysis and pseudotime data, and the flow cytometry experiments showed that M(Kp) is characterized by the increased expression of Arg1, Fizz1, iNOS, CD163, *cd206*, type I IFN and IL10 signalling-regulated genes, and the decreased expression of *ppar*γ, and inflammatory markers.

### K. pneumoniae-*induced M(Kp) polarization is STAT6 dependent*

We next sought to provide mechanistic insights into the signalling pathway(s) governing M(Kp) polarization. To facilitate this research, we questioned whether *K. pneumoniae* triggers M(Kp) polarization in immortalized bone marrow derived macrophages (iBMDMs). These cells have been widely used to investigate immune signalling and macrophage polarization. Consistent with the in vivo scRNA-seq results, infection of iBMDMs resulted in the upregulation of *arg1* and Arg1 (Supplementary Figure 7A), *fizz1* and Fizz1 ((Supplementary Figure 7B)*, ppar*γ, *nos2, il12*, *il6* and *tnfa* (Supplementary Figure76C-G). *Il10* levels were also upregulated in infected iBMDMs (Supplementary Figure 7H). The increased expression of *il10* was consistent with the increased phosphorylation of the IL10-governed transcriptional factor STAT3 in Kp52145-infected macrophages (Supplementary Figure 7I). We have previously demonstrated the upregulation of type I IFN-dependent genes in *Klebsiella*-infected iBMDMs (18). Altogether, these results demonstrate that infection of iBMDMs recapitulates the in vivo *K. pneumoniae*-induced macrophage polarization.

M2 macrophage polarization involves the activation of STAT6 that controls the transcription of M2-specific genes (22, 23). Therefore, we sought to determine whether STAT6 governs *K. pneumoniae*-induced M(Kp) polarization. We first investigated whether Kp52145 induced the phosphorylation of STAT6 because this is a prerequisite for nuclear localization and DNA binding of STAT6 (24). Immunoblotting experiments confirmed that Kp52145 induced STAT6 phosphorylation (Fig 5A). *K. pneumoniae* strains NJST258-1, NJST258-2 and SHG10 also induced the phosphorylation of STAT6 (Fig 5B), indicating that *K. pneumoniae* activation of STAT6 is not strain dependent. NJST258-1 and NJST258-2 cluster within the epidemic clonal group ST258 producing the *K. pneumoniae* carbapenemase, and SGH10 belongs to the clonal group CG23 causing liver abscesses (25, 26). STAT6 cooperates with KLF4 to regulate M2 macrophage (21). Kp52145 also increased the expression of *klf4* and KLF4 in macrophages (Fig 5C). To connect mechanistically STAT6 activation and *K. pneumoniae*-induced M(Kp) polarization, we infected *stat6^-/-^* macrophages and assessed the expression of M(Kp) markers. Figure 5D shows that *arg1* and Arg1 levels were decreased in infected *stat6^-/-^* macrophages compare to infected wild-type cells. Furthermore, Kp52145 did not upregulate the expression of *il10, klf4, pparg* and *fizz1* in *stat6^-/-^* macrophages (Fig 5 E-H). In contrast, the expressions of *nos2*, *tnfa, il12, il6* were higher in infected *stat6^-/-^* macrophages than in wild-type cells (Fig 5I-K). The levels of *isg15* were not significantly between infected wild-type and *stat6^-/-^* macrophages (Fig 5L). Flow cytometry experiments using mCherry-tagged Kp52145 demonstrated that neither Arg1 nor CD206 were upregulated in infected *stat6^-/-^* macrophages in contrast to wild-type cells with associated bacteria (Fig 5M-N). In contrast, the levels of MHC-II, a well-established M1 marker, were significantly upregulated in *stat6^-/-^* infected macrophages compare to wild-type cells (Fig 5O). Similar results were obtained when *K. pneumoniae*-induced STAT6 activation was supressed using the STAT6 inhibitor AS1517499 (27). When infections were performed in the presence of AS1517499, Kp52145 did not upregulate the expression of *arg1*, *il10,* and *fizz1* (Supplementary Figure 8A-C). In contrast, the expression of *nos2*, and *il12* were upregulated following infection (Supplementary Figure 8D-E). AS1517499 did not affect *Klebsiella*-induced *isg15* (Supplementary Figure 8F) in line with *stat6^-/-^*-infected cells.

**Figure 5.**
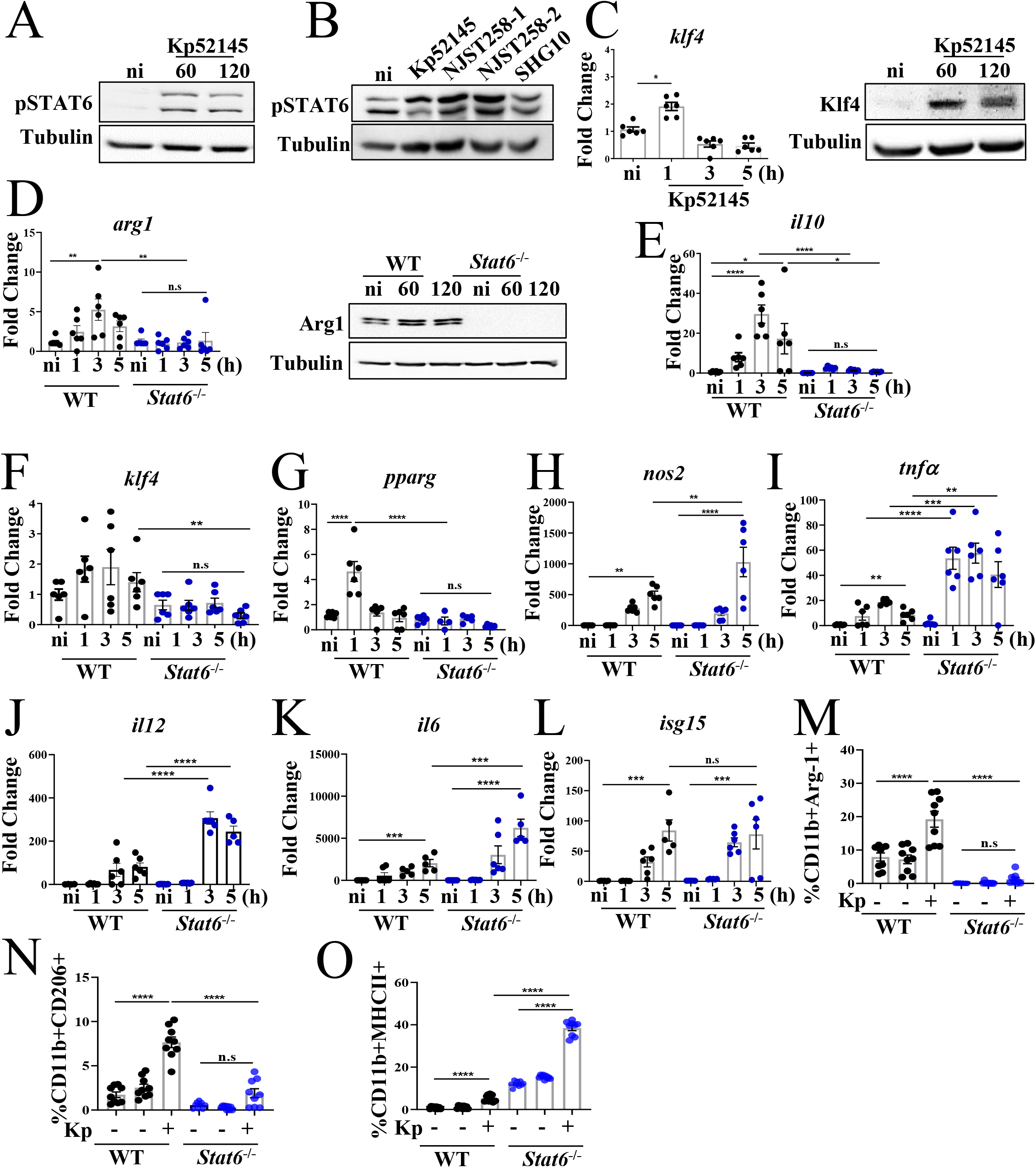
*K. pneumoniae*-induced M(Kp) polarization is STAT6 dependent. A. Immunoblot analysis of phospho-STAT6 (pSTAT6) and tubulin levels in lysates from non-infected (ni) and infected wild-type iBMDMs for 60 or 120 min. After 1 h contact, medium replaced with medium containing gentamycin (100 µg/ml) to kill extracellular bacteria. B. Immunoblot analysis of phospho-STAT6 (pSTAT6) and tubulin levels in lysates from non-infected (ni) and infected with different *K. pneumoniae* strains, Kp52145, NJST258-1, NJST258-2, or SHG10, for 60 min. C. *klf4* mRNA levels were assessed by qPCR in wild-type iBMDMs infected with Kp52145 for 1, 3 or 5 h. Immunoblot analysis of Klf4 and tubulin levels in lysates from non-infected (ni) and infected wild-type iBMDMs for 60 or 120 min. D. *arg1* mRNA levels were assessed by qPCR in wild-type and *stat6^-/-^* iBMDMs non-infected (ni) or infected with Kp52145 for 1, 3 or 5 h. Immunoblot analysis of Arg1 and tubulin levels in lysates from non-infected (ni) and infected wild-type and *stat6^-/-^* iBMDMs for 60 or 120 min. E. *il10* mRNA levels were assessed by qPCR in wild-type (WT) and *stat6^-/-^* iBMDMs non-infected (ni) or infected with Kp52145 for 1, 3 or 5 h. F. *klf4* mRNA levels were assessed by qPCR in wild-type (WT) and *stat6^-/-^* iBMDMs non-infected (ni) or infected with Kp52145 for 1, 3 or 5 h. G. *pparg* mRNA levels were assessed by qPCR in wild-type (WT) and *stat6^-/-^* iBMDMs non-infected (ni) or infected with Kp52145 for 1, 3 or 5 h. H. *nos2* mRNA levels were assessed by qPCR in wild-type (WT) and *stat6^-/-^* iBMDMs non-infected (ni) or infected with Kp52145 for 1, 3 or 5 h. I. *tnfa* mRNA levels were assessed by qPCR in wild-type (WT) and *stat6^-/-^* iBMDMs non-infected (ni) or infected with Kp52145 for 1, 3 or 5 h. J. *il12* mRNA levels were assessed by qPCR in wild-type (WT) and *stat6^-/-^* iBMDMs non-infected (ni) or infected with Kp52145 for 1, 3 or 5 h. K. *il6* mRNA levels were assessed by qPCR in wild-type (WT) and *stat6^-/-^* iBMDMs non-infected (ni) or infected with Kp52145 for 1, 3 or 5 h. L. *isg15* mRNA levels were assessed by qPCR in wild-type (WT) and *stat6^-/-^* iBMDMs non-infected (ni) or infected with Kp52145 for 1, 3 or 5 h. M. Percentage of wild-type (WT) and *stat6^-/-^* iBMDMs with and without associated Kp52145 positive for Arg1 1, 3 or 5 h post infection. Kp52145 was tagged with mCherry. N. Percentage of wild-type (WT) and *stat6^-/-^* iBMDMs with and without associated Kp52145 positive for CD206 1, 3 or 5 h post infection. Kp52145 was tagged with mCherry. O. Percentage of wild-type (WT) and *stat6^-/-^* iBMDMs with and without associated Kp52145 positive for MHC-II 1, 3 or 5 h post infection. Kp52145 was tagged with mCherry. For all infections, after 1 h contact, medium replaced with medium containing gentamycin (100 µg/ml) to kill extracellular bacteria. qPCR and flow cytometry values are presented as the mean ± SEM of three independent experiments measured in duplicate. Images are representative of three independent experiments. ****P ≤ 0.0001; ***P ≤ 0.001; **P 0.01; *P ≤ 0.05; ns, P > 0.05 for the indicated comparisons using one way-ANOVA with Bonferroni contrast for multiple comparisons test.

Collectively, these results demonstrate that STAT6 acts as a key regulator of *K. pneumoniae*-induced M(Kp) polarization. Furthermore, in the absence of STAT6 *K. pneumoniae* induces a macrophage phenotype consistent with M1 polarization.

### K. pneumoniae *activation of STAT6 promotes infection*

To establish the importance of *K. pneumoniae*-induced activation of STAT6 on *K. pneumoniae* infection biology, we first investigated whether STAT6 contributes to *K. pneumoniae* subversion of cell-intrinsic immunity. We asked whether absence of STAT6 impairs *K. pneumoniae* intracellular survival. While no differences were observed in the adhesion of Kp52145 to *stat6^-/-^* and wild-type macrophages (Fig 6A), the phagocytosis of Kp52145 was reduced in *stat6^-/-^* macrophages compared to wild-type cells (Fig 6B). Time course experiments showed that the intracellular survival of Kp52145 was diminished in *stat6^-/-^* macrophages (Fig 6C). Previously, we have demonstrated that *K. pneumoniae* manipulates the traffic of the phagosome following phagocytosis to create a vacuole that does not fuse with lysosomes, the KCV, allowing the intracellular survival of *Klebsiella* (28). We then sought to determine whether the reduced intracellular survival observed in *stat6^-/-^* cells was due to an increase colocalization of lysosomes with the KCV. Lysosomes were labelled with the membrane-permeant fluorophore cresyl violet (29), and cells were infected with GFP-labelled Kp52145 to assess the KCV at the single cell level by immunofluorescence. Confocal microscopy experiments revealed that the majority of the KCVs from wild-type macrophages did not colocalize with cresyl violet (Fig 6D and Fig 6E), corroborating our previous work (28). In contrast, there was an increase of colocalization of the KCV from *stat6^-/-^* macrophages with the marker cresyl violet (Fig 6D and Fig 6E), demonstrating that the absence of STAT6 results in the fusion of the KCV with lysosomes with a concomitant reduction in the numbers of intracellular bacteria.

**Figure 6.**
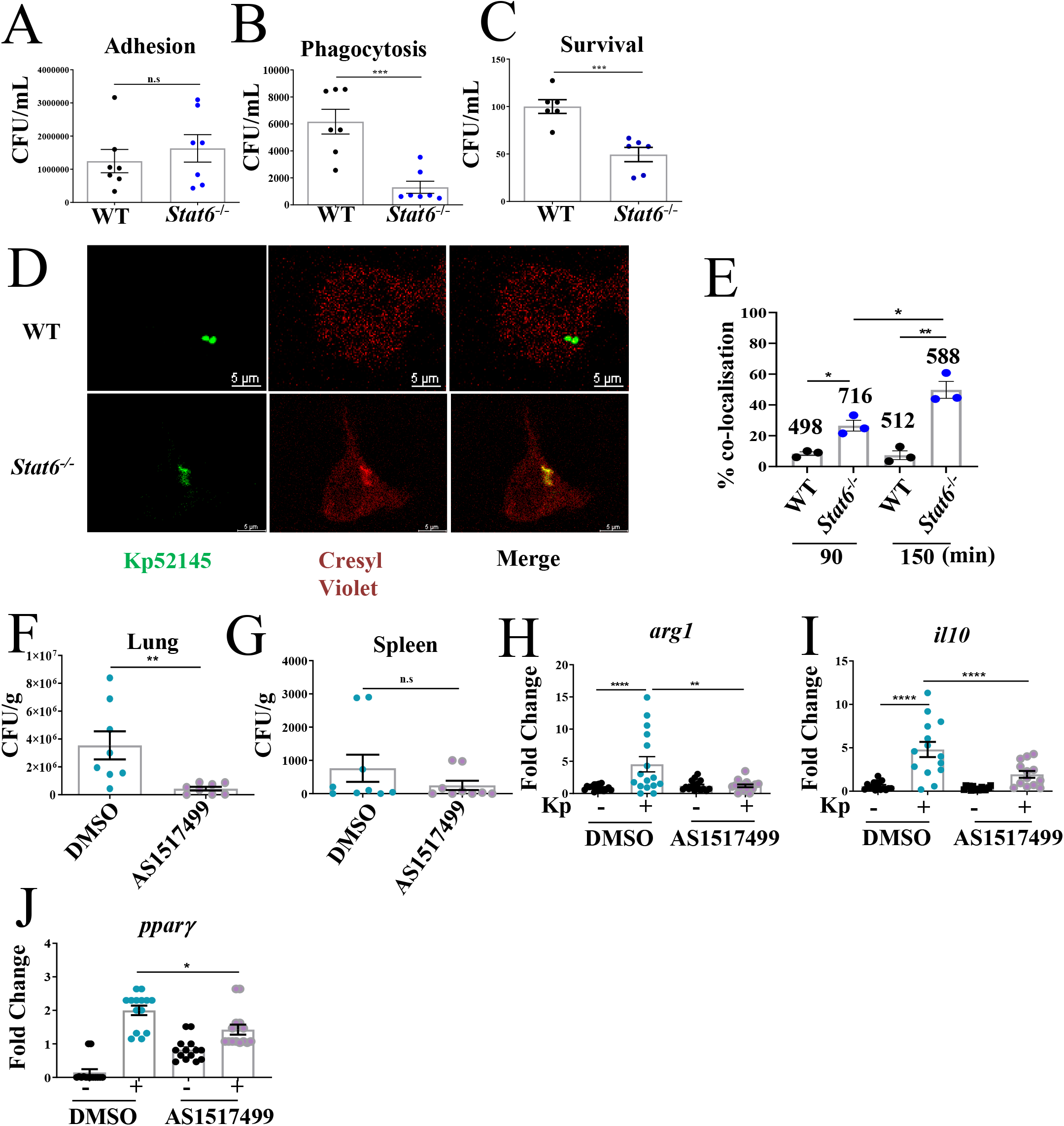
STAT6 promotes *K. pneumoniae* infection. A. Kp52145 adhesion to wild-type (WT) and *stat6^-/-^* iBMDMs. Cells were infected with Kp52145 for 30 min, washed, cell lysed with saponin, and bacteria quantified after serial dilution followed by plating on LB agar plates. B. Phagocytosis of Kp52145 by wild-type (WT) and *stat6^-/-^* iBMDMs. Cells were infected for 30 min, wells were washed, and it was added medium containing gentamicin (100 µg/ml) to kill extracellular bacteria. After 30 min, cells were washed, cell lysed with saponin, and bacteria quantified after serial dilution followed by plating on LB agar plates. C. Kp52145 intracellular survival in wild-type (W)T and *stat6^-/-^* 5 h after addition of gentamycin (30 min of contact). Results are expressed as % of survival (CFUs at 5 h versus 30 min in *stat6^-/-^* cells normalized to the results obtained in wild-type macrophages set to 100%). D. Immunofluorescence confocal microscopy of the colocalization of Kp52145 harbouring pFPV25.1Cm, and cresyl violet dye in wild-type (WT) and *stat6^-/-^* macrophages. The images were taken 90 min post infection. Images are representative of duplicate coverslips of three independent experiments. E. Percentage of Kp52145 harbouring pFPV25.1Cm co-localization with cresyl violet over a time course. Wild-type (WT) and *stat6^-/-^* macrophages were infected; coverslips were fixed and stained at the indicated times. Values are given as mean percentage of Kp52145 co-localizing with the marker☐±☐SEM. The number of infected cells counted per time in three independent experiments are indicated in the figure. F. C56BL/6 mice were treated 24 h prior to infection with the STAT6 inhibitor AS1517499 (10mg/kg in 200 μL volume by i.p) and 6 h post infection (5 mg/kg in 30 μl volume intranasally) or vehicle control DMSO. Bacterial burden was established by serial dilutions of lung homogenates on *Salmonella-Shigella* agar. Each dot represents one animal. G. Bacterial dissemination assessed by quantifying CFUs in the spleens from infected mice treated with AS1517499 or vehicle control DMSO. Each dot represents one animal. H. *arg1* mRNA levels were assessed by qPCR in the lungs of non-infected or infected wild-type mice for 24 h treated with the STAT6 inhibitor AS1517499 or DMSO vehicle control. Each dot represents different mice. I. *il10* mRNA levels were assessed by qPCR in the lungs of non-infected or infected wild-type mice for 24 h treated with the STAT6 inhibitor AS1517499 or DMSO vehicle control. Each dot represents different mice. J. *pparg* mRNA levels were assessed by qPCR in the lungs of non-infected or infected wild-type mice for 24 h treated with the STAT6 inhibitor AS1517499 or DMSO vehicle control. Each dot represents different mice. Values are presented as the mean ± SEM of three independent experiments measured in duplicate. In panels A, B, C, E, F and G, unpaired t test was used to determine statistical significance. In panels H, I and J, statistical analysis were carried out using one-way ANOVA with Bonferroni contrast for multiple comparisons test. ****P ≤ 0.0001; ***P ≤ 0.001; ** P ≤ 0.01; ns, P > 0.05 for the indicated comparisons determined using unpaired t test. ****P ≤ 0.0001; ***P ≤ 0.001; ** P ≤ 0.01; *P ≤ 0.05; ns, P > 0.05 for the indicated comparisons.

To obtain a global view of the role of STAT6 in *K. pneumoniae* infection biology, we examined the effect of the STAT6 inhibitor AS1517499 on the ability of wild-type mice to control *K. pneumoniae* infection. At 24 h post infection, the bacterial loads in the lungs of mice pre-treated with AS1517499 were significantly lower than those of mice pre-treated with the vehicle solution (Fig 6F). Bacterial loads in the spleens were not significantly different between the two groups of mice (Fig 6G). The expression of M2 polarisation markers *arg1, il10* and *pparg* were significantly reduced in whole lungs from infected animals treated with AS1517499 compared to DMSO controls (Fig 6H and Fig 6J). These results establish the importance of STAT6 activation for *K. pneumoniae* survival *in vivo*.

### K. pneumoniae-*induced M(Kp) polarization is governed by TLR2 and TLR4 signalling*

We next sought to identify the signalling pathway(s) utilized by *K. pneumoniae* to activate STAT6 to induce M(Kp) polarization. Previous work of our laboratory demonstrates that *K. pneumoniae* manipulates pattern recognition receptors (PRRs) as a virulence strategy to control inflammation (30). We then asked whether *K. pneumoniae* may exploit TLR signalling to activate STAT6 to induce M(Kp). Kp52145 did not induced the phosphorylation of STAT6 in *tlr2^-/-^*, *tlr4^-/-^*, and *tlr2/tlr4^-/-^* macrophages (Fig 7A). Consistent with the lack of activation of STAT6 in macrophages lacking TLR2 and TLR4, Kp52145 did not increase the levels of Arg1 (Fig 7B), *arg1* (Fig 7C), and *fizz1* (Fig 7D) in *tlr2^-/-^*, *tlr4^-/-^*, and *tlr2/4^-/-^* macrophages. The expressions of *il10*, *nos2* and the ISGs *isg15*, and *mx1* were only upregulated in *tlr2^-/-^* macrophages following infection (Fig 7E-H), indicating that TLR4 controls the levels of these M(Kp) markers. In contrast, TLR2 controls the expression of *ppar*γ because Kp52145 did not upregulate *ppar*γ in *tlr2* and *tlr2/tlr4* macrophages (Fig 7I). Altogether, these results demonstrate that *K. pneumoniae*-induced M(Kp) polarization is TLR2 and TLR4 dependent.

**Figure 7.**
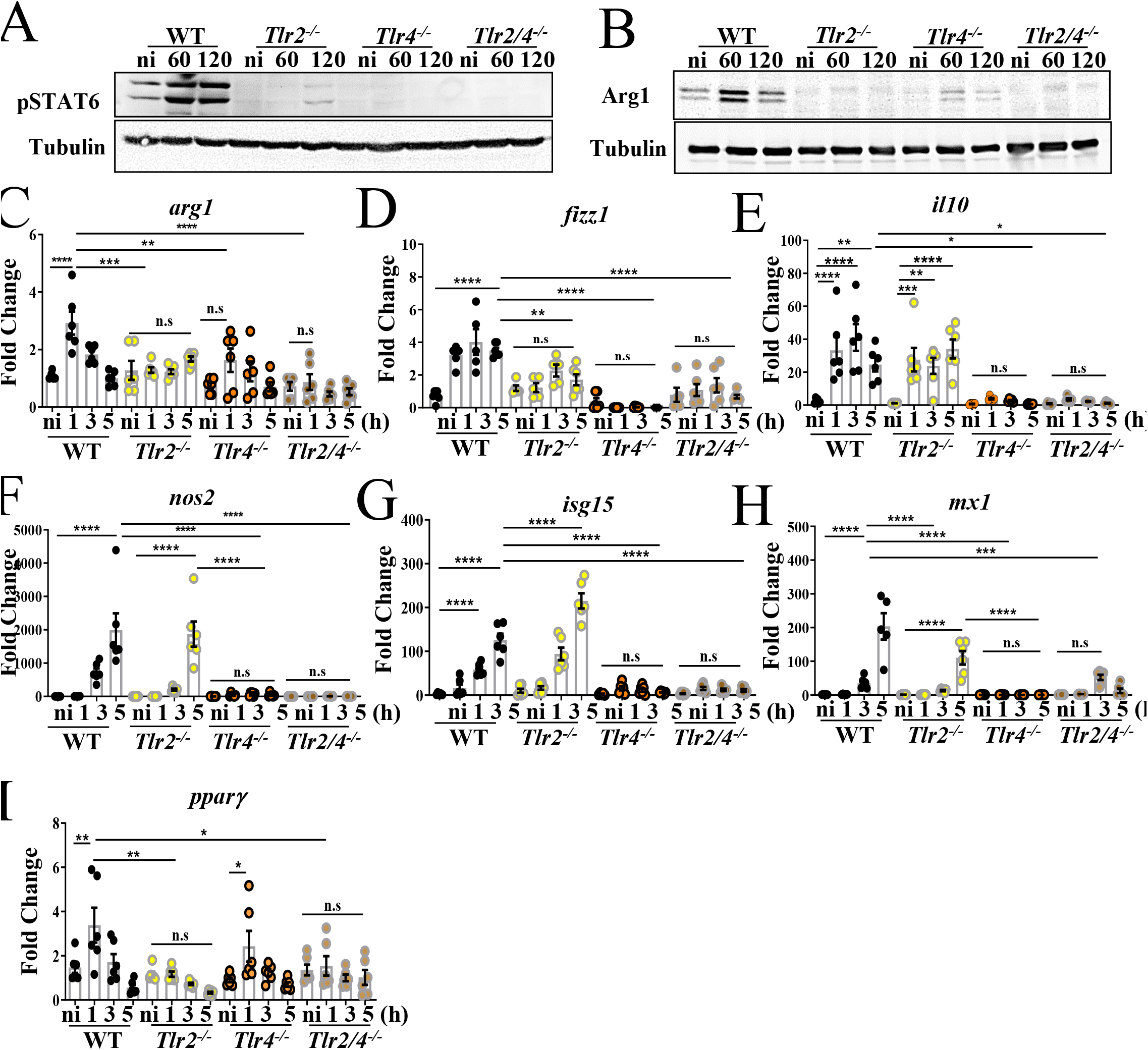
*K. pneumoniae*-induced M (Kp) polarisation is dependent on TLR signalling. A. Immunoblot analysis of phospho-STAT6 (pSTAT6) and tubulin levels in lysates from non-infected (ni) and infected wild-type (WT), *tlr2^-/-^*, *tlr4^-/-^* and *tlr2/4^-/-^* iBMDMs for 60 or 120 min. B. Immunoblot analysis of Arg1and tubulin levels in lysates from non-infected (ni) and infected wild-type (WT), *tlr2^-/-^*, *tlr4^-/-^* and *tlr2/4^-/-^* iBMDMs for 60 or 120 min. C. *arg1* mRNA levels were assessed by qPCR in wild-type (WT) and *tlr2^-/-^*, *tlr4^-/-^* and *tlr2/4^-/-^* iBMDMs non-infected (ni) or infected with Kp52145 for 1, 3 or 5 h. D. *fizz1* mRNA levels were assessed by qPCR in wild-type (WT) and *tlr2^-/-^*, *tlr4^-/-^* and *tlr2/4^-/-^* iBMDMs non-infected (ni) or infected with Kp52145 for 1, 3 or 5 h. E. *il10* mRNA levels were assessed by qPCR in wild-type (WT) and *tlr2^-/-^*, *tlr4^-/-^* and *tlr2/4^-/-^* iBMDMs non-infected (ni) or infected with Kp52145 for 1, 3 or 5 h. F. *nos2* mRNA levels were assessed by qPCR in wild-type (WT) and *tlr2^-/-^*, *tlr4^-/-^* and *tlr2/4^-/-^* iBMDMs non-infected (ni) or infected with Kp52145 for 1, 3 or 5 h. G. *isg15* mRNA levels were assessed by qPCR in wild-type (WT) and *tlr2^-/-^*, *tlr4^-/-^* and *tlr2/4^/-^* iBMDMs non-infected (ni) or infected with Kp52145 for 1, 3 or 5 h. H. *mx1* mRNA levels were assessed by qPCR in wild-type (WT) and *tlr2^-/-^*, *tlr4^-/-^* and *tlr2/4^-/-^* iBMDMs non-infected (ni) or infected with Kp52145 for 1, 3 or 5 h. I. *pparg* mRNA levels were assessed by qPCR in wild-type (WT) and *tlr2^-/-^*, *tlr4^-/-^* and *tlr2/4^-/-^*iBMDMs non-infected (ni) or infected with Kp52145 for 1, 3 or 5 h. For all infections, after 1 h contact, medium replaced with medium containing gentamycin (100 µg/ml) to kill extracellular bacteria. Error bars are presented as the mean ± SEM of three independent experiments in duplicate. Images are representative of three independent experiments. ****P ≤ 0.0001; ***P ≤ 0.001; **P 0.01; *P ≤ 0.05; ns, P > 0.05 for the indicated comparisons using one way-ANOVA with Bonferroni contrast for multiple comparisons test.

### *The TLR adaptors MyD88, TRAM, and TRIF govern* K. *pneumoniae-induced M(Kp) polarization*

TLR signalling involves a series of different adaptors. Myeloid-Differentiation factor-88 (MyD88) is a universal adaptor used by all TLRs except TLR3, Toll/IL-1R domain-containing adaptor-inducing IFN-β (TRIF) is used by TLR3 and TLR4, whereas TRIF-related adaptor molecule (TRAM) is recruited by endosomal located TLR and TLR4 (31). Therefore, we asked whether MyD88, TRAM, and TRIF are required to induce the M(Kp) polarization. Phosphorylation of STAT6 was not detected in infected *myd88^-/-^*, and *tram/trif^-/-^* macrophages (Fig 8A). As anticipated Arg1 (Fig 8B) and *arg1* (Fig 8C) were not upregulated in infected *myd88^-/-^*, and *tram/trif^-/-^* macrophages. Kp52145 induction of *il10* was MyD88-dependent because Kp52145 upregulated *il10* only in *tram/trif^-/-^* macrophages (Fig 8D). The expressions of *pparg* and *fizz1* were abrogated in the absence of Myd88 and TRAM/TRIF (Fig 8E-F). In addition, *isg15* and *mx1*were not upregulated in infected *tram/trif^-/-^* macrophages (Fig 8G-H), which is consistent with our recent evidence demonstrating that *K. pneumoniae* induces type I IFNs and ISGs in a TRAM-TRIF-dependent manner (18). Kp52145 did not upregulate *nos2* in *tram/trif^-/-^* macrophages (Fig 8I) which is in agreement with *nos2* being an ISG (32). In contrast, *nos2* levels were upregulated in infected *myd88^-/-^* macrophages (Fig 8I). This result is consistent with the facts that MyD88 signalling is needed for *K. pneumoniae* induction of *il10* (Fig 8D) and IL10 reduces the levels of *nos2* in *K. pneumoniae*-infected macrophages (Fig 10H). Altogether, these findings establish that *K. pneumoniae*-induced M(Kp) is MyD88, TRAM and TRIF dependent.

**Figure 8.**
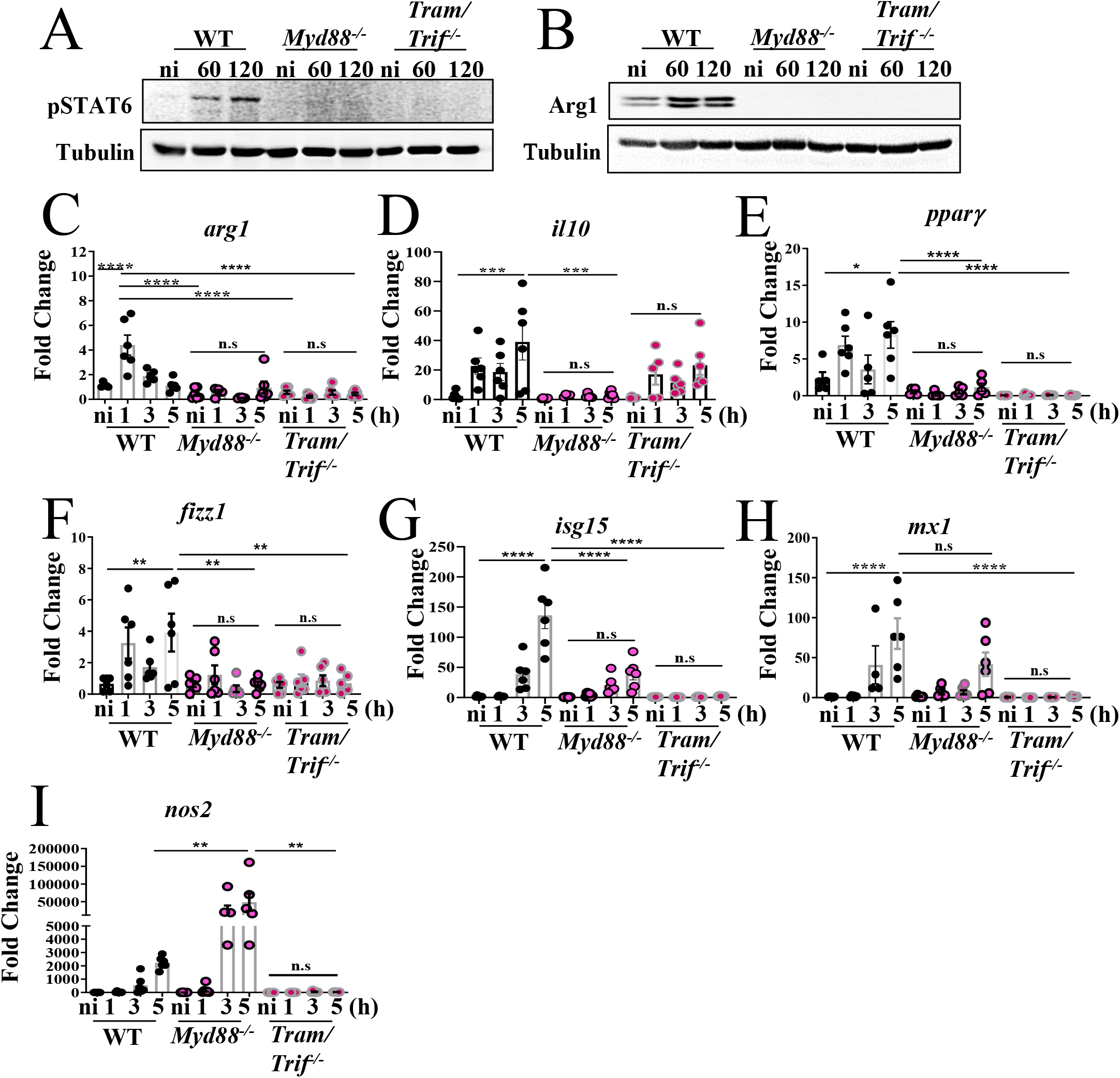
*K. pneumoniae*-induced M (Kp) polarisation is dependent on the TLR adaptors MyD88, TRAM and TRIF. A. Immunoblot analysis of phospho-STAT6 (pSTAT6) and tubulin levels in lysates from non-infected (ni) and infected wild-type (WT), *myd88^-/-^*, *tram/trif^-/-^* for 60 or 120 min. B. Immunoblot analysis of Arg1 and tubulin levels in lysates from non-infected (ni) and infected wild-type (WT), *myd88^-/-^*, *tram/trif^-/-^* for 60 or 120 min. C. *arg1* mRNA levels were assessed by qPCR in wild-type (WT), *myd88^-/-^*, *tram/trif^-/-^* non-infected (ni) or infected with Kp52145 for 1, 3 or 5 h. D. *il10* mRNA levels were assessed by qPCR in wild-type (WT), *myd88^-/-^*, *tram/trif^-/-^* non-infected (ni) or infected with Kp52145 for 1, 3 or 5 h. E. *pparg* mRNA levels were assessed by qPCR in wild-type (WT), *myd88^-/-^*, *tram/trif^-/-^* non-infected (ni) or infected with Kp52145 for 1, 3 or 5 h. F. *fizz1* mRNA levels were assessed by qPCR in wild-type (WT), *myd88^-/-^*, *tram/trif^-/-^* non-infected (ni) or infected with Kp52145 for 1, 3 or 5 h. G. *isg15* mRNA levels were assessed by qPCR in wild-type (WT), *myd88^-/-^*, *tram/trif^-/-^* non-infected (ni) or infected with Kp52145 for 1, 3 or 5 h. H. *mx1* mRNA levels were assessed by qPCR in wild-type (WT), *myd88^-/-^*, *tram/trif^-/-^* non-infected (ni) or infected with Kp52145 for 1, 3 or 5 h. I. *nos2* mRNA levels were assessed by qPCR in wild-type (WT), *myd88^-/-^*, *tram/trif^-/-^* non-infected (ni) or infected with Kp52145 for 1, 3 or 5 h. For all infections, after 1 h contact, medium replaced with medium containing gentamycin (100 µg/ml) to kill extracellular bacteria. Error bars are presented as the mean ± SEM of three independent experiments in duplicate. Images are representative of three independent experiments. ****P ≤ 0.0001; ***P ≤ 0.001; **P 0.01; *P ≤ 0.05; ns, P > 0.05 for the indicated comparisons using one way-ANOVA with Bonferroni contrast for multiple comparisons test.

### K. pneumoniae *exploits type I IFN signalling to induce M(Kp) polarization*

Given that TLR4-TRAM-TRIF pathway is essential for M(Kp) activation, that this pathway controls type I IFN signalling in *K. pneumoniae* infections (18), and that type I IFN signalling is a feature of IMs following infection, we decided to elucidate whether *K. pneumoniae* exploits type I IFN to induce M(Kp) polarization. To address this question, we infected type I IFN receptor-deficient (*ifnar1^-/-^*) macrophages and assessed M(Kp) markers. Kp52145 did not induce the phosphorylation of STAT6 in *ifnar1^-/-^* macrophages (Fig 9A). As expected, Arg1 and *arg1* were not upregulated in Kp52145-infected *ifnar1^-/-^* cells (Fig 9B). Similar result was observed for *nos2*, *ppar*γ and *fizz1* (Fig 9 C-E). In contrast, Kp52145 still upregulated *il10* in *ifnar1^-/-^* macrophages (Fig 9F). Flow cytometry experiments showed that Kp52145 did not upregulate the expression of Arg1 (Fig 9G), and CD206 in *ifnar1^-/-^* cells (Fig 9H) whereas the expression of MHC-II was higher in *ifnar1^-/-^* macrophages than in wild-type cells following infection (Fig 9I). Type I IFN stimulation alone did not induce STAT6 phosphorylation nor upregulate Arg1 in wild-type cells (Supplementary Figure 9A-B).

**Figure 9.**
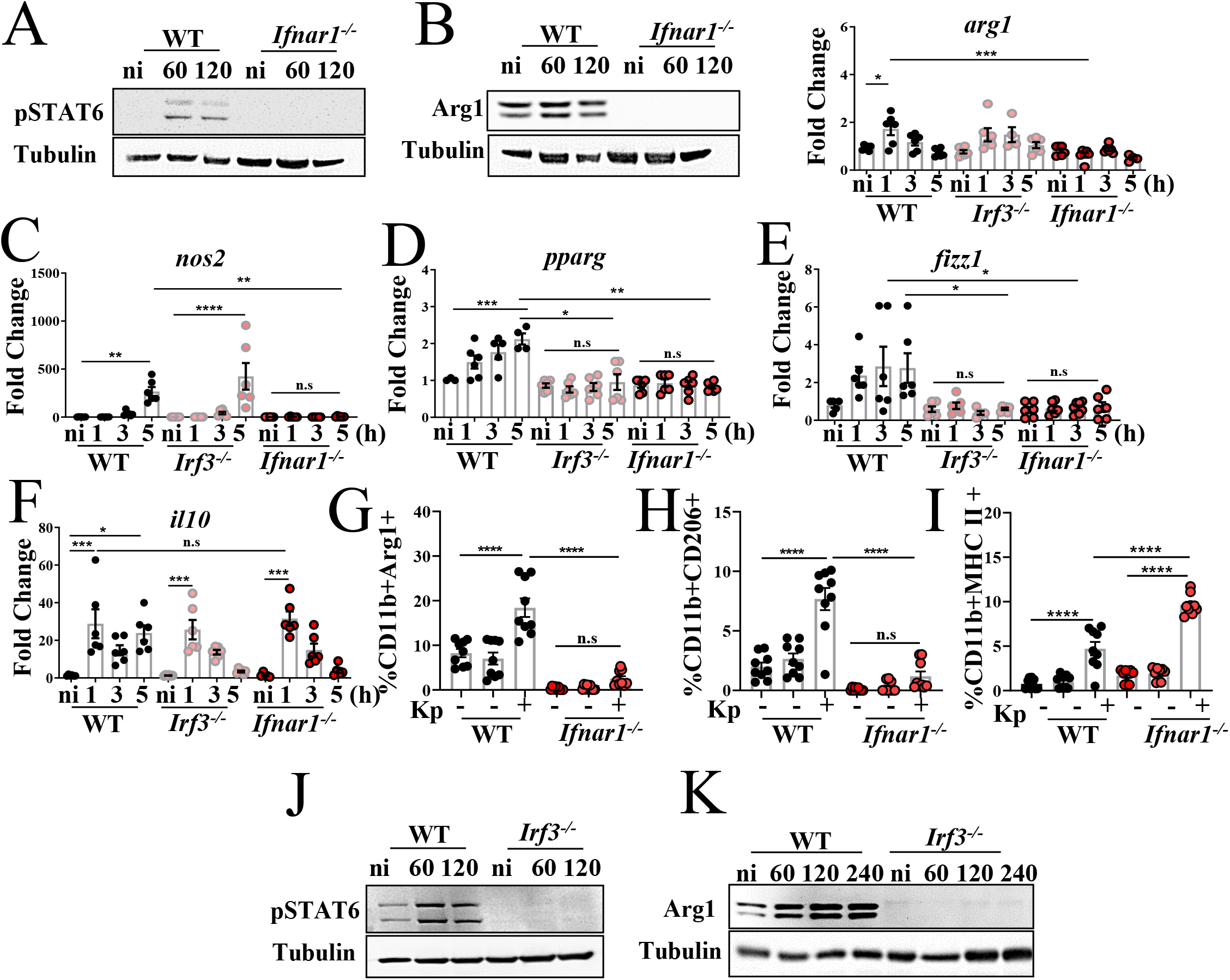
*K. pneumoniae* exploits type I IFN signalling to induce M(Kp) polarisation. A. Immunoblot analysis of phospho-STAT6 (pSTAT6) and tubulin levels in lysates from non-infected (ni) and infected wild-type (WT), or *ifnar1^-/-^* for 60 or 120 min. B. Immunoblot analysis of Arg1 and tubulin levels in lysates from non-infected (ni) and infected wild-type (WT), or *ifnar1^-/-^* for 60 or 120 min. *arg1* mRNA levels were assessed by qPCR in wild-type (WT), *irf3 ^-/-^*, *ifnar1^-/-^* non-infected (ni) or infected with Kp52145 for 1, 3 or 5 h. C. *nos2* mRNA levels were assessed by qPCR in wild-type (WT), *irf3 ^-/-^*, *ifnar1^-/-^* non-infected (ni) or infected with Kp52145 for 1, 3 or 5 h. D. *pparg* mRNA levels were assessed by qPCR in wild-type (WT), *irf3 ^-/-^*, *ifnar1^-/-^* non-infected (ni) or infected with Kp52145 for 1, 3 or 5 h. E. *fizz1* mRNA levels were assessed by qPCR in wild-type (WT), *irf3 ^-/-^*, *ifnar1^-/-^* non-infected (ni) or infected with Kp52145 for 1, 3 or 5 h. F. *il10* mRNA levels were assessed by qPCR in wild-type (WT), *irf3 ^-/-^*, *ifnar1^-/-^* non-infected (ni) or infected with Kp52145 for 1, 3 or 5 h. G. Percentage of wild-type (WT) and *ifnar1^-/-^* iBMDMs with and without associated Kp52145 positive for Arg1 1, 3 or 5 h post infection. Kp52145 was tagged with mCherry. H. Percentage of wild-type (WT) and *ifnar1^-/-^* iBMDMs with and without associated Kp52145 positive for CD206 1, 3 or 5 h post infection. Kp52145 was tagged with mCherry. I. Percentage of wild-type (WT) and *ifnar1^-/-^* iBMDMs with and without associated Kp52145 positive for MHCII 1, 3 or 5 h post infection. Kp52145 was tagged with mCherry. J. Immunoblot analysis of phospho-STAT6 (pSTAT6) and tubulin levels in lysates from non-infected (ni) and infected wild-type (WT), or *irf3^-/-^* for 60 or 120 min. K. Immunoblot analysis of Arg1 and tubulin levels in lysates from non-infected (ni) and infected wild-type (WT), or *irf31^-/-^* for 60 or 120 min. For all infections, after 1 h contact, medium replaced with medium containing gentamycin (100 µg/ml) to kill extracellular bacteria. Error bars are presented as the mean ± SEM of three independent experiments in duplicate. Images are representative of three independent experiments. ****P ≤ 0.0001; ***P ≤ 0.001; **P≤ 0.01; *P ≤ 0.05; ns, P > 0.05 for the indicated comparisons using one way-ANOVA with Bonferroni contrast for multiple comparisons test.

**Figure 10.**
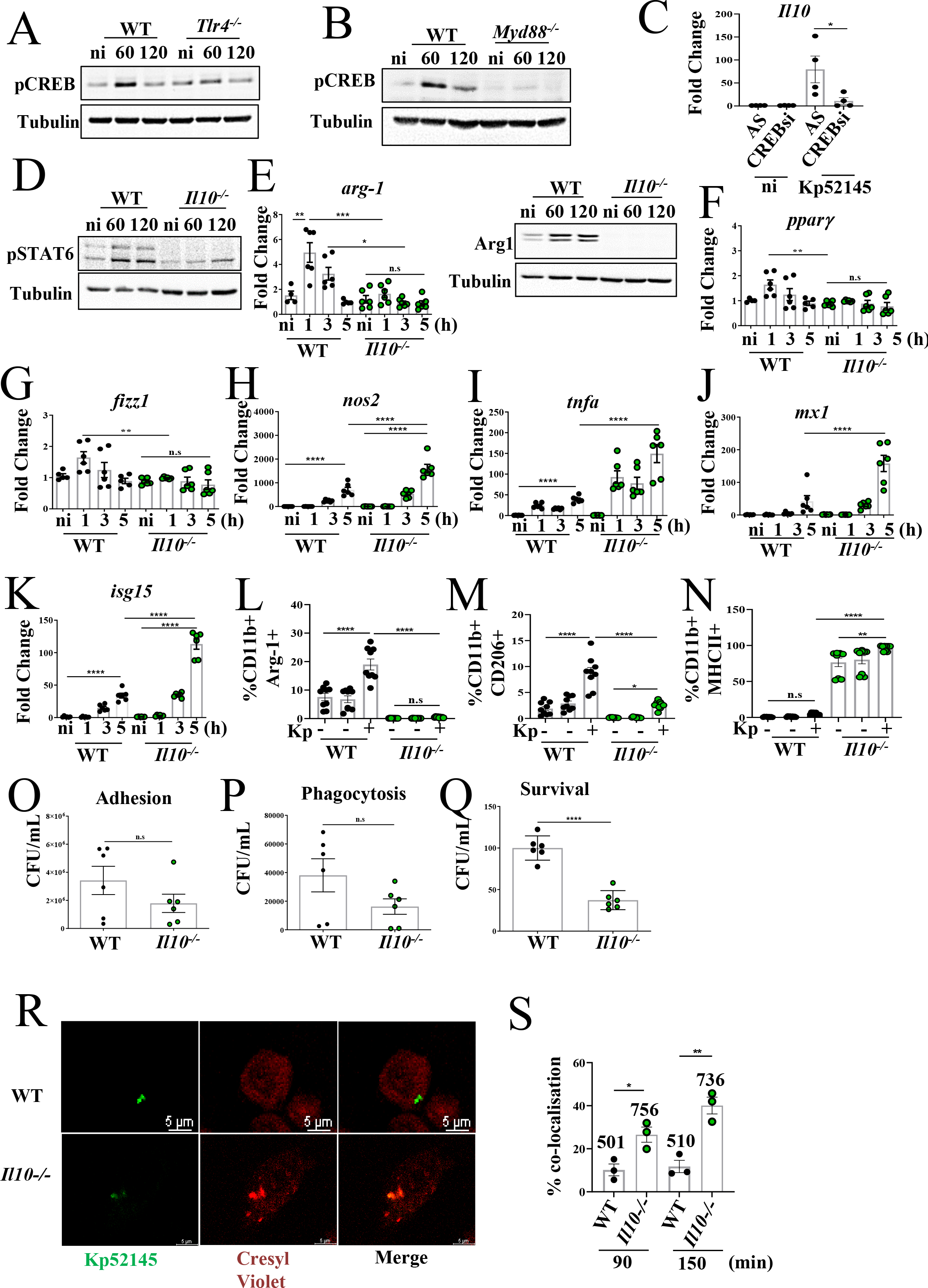
*K. pneumoniae*-induced M(Kp) polarisation is dependent on IL10. A. Immunoblot analysis of phospho-CREB (pCREB) and tubulin levels in lysates from non-infected (ni) and infected wild-type (WT), or *tlr4^-/-^* for 60 or 120 min. B. Immunoblot analysis of phospho-CREB (pCREB) and tubulin levels in lysates from non-infected (ni) and infected wild-type (WT), or *myd88^-/-^* for 60 or 120 min. C. *il0* mRNA levels were assessed by qPCR in iBMDMs transfected with All Stars siRNA control (AS), or CREB siRNA (CREBsi) non-infected (ni) or infected with Kp52145 for 3 h. After 1 h contact, the medium was replaced with medium containing gentamicin (100 µg/ml) to kill extracellular bacteria. D. Immunoblot analysis of phospho-STAT6 (pSTAT6) and tubulin levels in lysates from non-infected (ni) and infected wild-type (WT), or *il10^-/-^* for 60 or 120 min. E. *arg1* mRNA levels were assessed by qPCR in wild-type (WT), *il10^-/-^* non-infected (ni) or infected with Kp52145 for 1, 3 or 5 h. Immunoblot analysis of Arg1 and tubulin levels in lysates from non-infected (ni) and infected wild-type (WT), or *il10^-/-^* for 60 or 120 min. F. *pparg* mRNA levels were assessed by qPCR in wild-type (WT), *il10^-/-^* non-infected (ni) orinfected with Kp52145 for 1, 3 or 5 h. G. *fizz1* mRNA levels were assessed by qPCR in wild-type (WT), *il10^-/-^* non-infected (ni) or infected with Kp52145 for 1, 3 or 5 h. H. *nos2* mRNA levels were assessed by qPCR in wild-type (WT), *il10^-/-^* non-infected (ni) or infected with Kp52145 for 1, 3 or 5 h. I. *tnfa* mRNA levels were assessed by qPCR in wild-type (WT), *il10^-/-^* non-infected (ni) or infected with Kp52145 for 1, 3 or 5 h. J. *mx1* mRNA levels were assessed by qPCR in wild-type (WT), *il10^-/-^* non-infected (ni) or infected with Kp52145 for 1, 3 or 5 h. K. *isg15* mRNA levels were assessed by qPCR in wild-type (WT), *il10^-/-^* non-infected (ni) or infected with Kp52145 for 1, 3 or 5 h. L. Percentage of wild-type (WT) and *il10^-/-^* iBMDMs with and without associated Kp52145 positive for Arg1 1, 3 or 5 h post infection. Kp52145 was tagged with mCherry. M. Percentage of wild-type (WT) and *il10^-/-^* iBMDMs with and without associated Kp52145 positive for CD206 1, 3 or 5 h post infection. Kp52145 was tagged with mCherry. N. Percentage of wild-type (WT) and *il10^-/-^* iBMDMs with and without associated Kp52145 positive for MHCII 1, 3 or 5 h post infection. Kp52145 was tagged with mCherry. O. Kp52145 adhesion to wild-type (WT) and *il10^-/-^* iBMDMs. Cells were infected with Kp52145 for 30 min, washed, cell lysed with saponin, and bacteria quantified after serial dilution followed by plating on LB agar plates. P. Phagocytosis of Kp52145 by wild-type (WT) and *il10^-/-^* iBMDMs. Cells were infected for 30 min, wells were washed, and it was added medium containing gentamicin (100 µg/ml) to kill extracellular bacteria. After 30 min, cells were washed, cell lysed with saponin, and bacteria quantified after serial dilution followed by plating on LB agar plates. Q. Kp52145 intracellular survival in wild-type (W)T and *il10^-/-^* 5 h after addition of gentamycin (30 min of contact). Results are expressed as % of survival (CFUs at 5 h versus 30 min in *stat6^-/-^* cells normalized to the results obtained in wild-type macrophages set to 100%). R. Immunofluorescence confocal microscopy of the colocalization of Kp52145 harbouring pFPV25.1Cm, and cresyl violet dye in wild-type (WT) and *il10^-/-^* macrophages. The images were taken 90 min post infection. Images are representative of duplicate coverslips of three independent experiments. S. Percentage of Kp52145 harbouring pFPV25.1Cm co-localization with cresyl violet over a time course. Wild-type (WT) and *il10^-/-^* macrophages were infected; coverslips were fixed and stained at the indicated times. Values are given as mean percentage of Kp52145 co-localizing with the marker☐±☐SEM. The number of infected cells counted per time in three independent experiments are indicated in the figure. For all infections, after 1 h contact, medium replaced with medium containing gentamycin (100 µg/ml) to kill extracellular bacteria. Error bars are presented as the mean ± SEM of three independent experiments in duplicate. Images are representative of three independent experiments. In panels O, P, and Q unpaired t test was used to determine statistical significance. In all the other panels, statistical analysis were carried out using one-way ANOVA with Bonferroni contrast for multiple comparisons test. ****P ≤ 0.0001; ***P ≤ 0.001; **P≤ 0.01; *P ≤ 0.05; ns, P > 0.05 for the indicated comparisons.

Because Irf3 controls type I IFN production in *K. pneumoniae* in vitro and in vivo (18), we postulated that Irf3 is required for *K. pneumoniae* induction of M(Kp). Indeed, Kp52145 did not phosphorylate STAT6 or induced Arg1 in *irf3^-/-^* macrophages (Fig 9J-K). As anticipated, *arg1* (Fig 9B)*, ppar*γ (Fig 9D), and *fizz1* (Fig 9E) levels were not increase in *irf3^-/-^* cells following infection whereas the levels of *il10* were similar that those found in infected wild-type cells (Fig 9F). The fact that *nos2* levels were upregulated in infected *irf3^-/-^* macrophages (Fig 9C) indicates that Irf3 does not control the transcription of this gene.

Collectively, these experiments demonstrate that *K. pneumoniae* leverages the immunomodulatory properties of type I IFN to induce M(Kp) polarization.

IL10 is required for K. pneumoniae-governed M(Kp).

Our findings indicate that IL10 is one of the signatures of M(Kp). Therefore, we sought to identify the signalling pathways governing *K. pneumoniae*-induction of IL10. Our previous results revealed that TLR4-MyD88 signalling controls IL10 production following Kp52145 infection (Fig 7E and Fig 8D). The fact that the transcriptional factor CREB controls IL10 production in macrophages following TLR stimulation (33) led us to ascertain whether CREB governs *K. pneumonia-*induced IL10 production. The phosphorylation of CREB is a key event regulating its transcriptional activity (34). Immunoblotting experiments confirmed that Kp52145 triggered the phosphorylation of CREB in wild-type macrophages (Fig 10A and Fig 10B). However, CREB phosphorylation was reduced in infected *tlr4^-/-^* (Fig 10A) and *myd88^-/-^* (Fig 10B) macrophages. To connect CREB activation and IL10 production in *K. pneumoniae* infected cells, we used a siRNA-based approach to knockdown CREB in macrophages. The efficiency of CREB knockdown in wild-type macrophages is shown in (Supplementary Figure 10). Kp52145 induction of *il10* was abrogated in CREB knockdown macrophages (Fig 10C), demonstrating the role of CREB activation in *K. pneumoniae* induction of IL10. Collectively, these experiments uncover that a TLR4-MyD88-CREB signalling pathway mediates *K. pneumoniae* induction of IL10.

To determine whether *K. pneumoniae* exploits the immunomodulatory properties of IL10 to skew macrophage polarization, we infected *il10^-/-^* macrophages and assessed different M(Kp) markers. We did not detect the phosphorylation of STAT6 in infected *il10^-/-^* macrophages (Fig 10D). Consistent with the lack of activation of STAT6, the levels of *arg1*, Arg1 were not upregulated in infected *il10^-/-^* macrophages (Fig 10E). The expressions of *pparγ* (Fig 10F) and *fizz1* (Fig 10G) were not upregulated in infected *il10^-/-^* macrophages. In contrast, the levels of *nos2*, *tnf*α*, mx1* and *isg15* were significantly increased in infected *il10^-/-^* macrophages compared to wild-type controls (Fig 10H and Fig 10K). Flow cytometric analysis showed that Kp52145 did not increase Arg1 (Fig 10L) and CD206 (Fig 10M) in *il10^-/-^* macrophages whereas the levels of MHC-II were higher in *il10^-/-^* macrophages,with and without bacteria, than in the wild-type ones (Fig 10N). Recombinant IL10 neither alone nor in combination with type I IFN induced the phosphorylation of STAT6 or the upregulation of Arg1 in wild-type macrophages (Supplementary Figure 9A-B). Altogether, these data indicate that IL10 is necessary for *K. pneumoniae*-induction of M(Kp).

Given the importance of IL10 in *K. pneumoniae*-macrophage interplay, we asked whether IL10 is required for *K. pneumoniae* intracellular survival. No differences were observed in the adhesion (Fig 10O) and phagocytosis (Fig 10P) of Kp52145 between wild-type and *il10^-/-^* macrophages. Assessment of the numbers of intracellular bacteria over time showed a 50% decrease of Kp52145 survival in *il10^-/-^* macrophages (Fig 10Q). Confocal microscopy experiments showed an increase in the colocalization of the KCV from *il10^-/-^* macrophages with cresyl violet (Fig 10R and Fig 10S), demonstrating that the lack of IL10 results in the fusion of the KCV with lysosomes with a concomitant reduction in the numbers of intracellular bacteria.

In summary, our data demonstrates that *K. pneumoniae* exploits IL10 following the activation of a TLR4-MyD88-CREB pathway to induce M(Kp) polarization. IL10 is crucial for *K. pneumoniae*-governed control of the phagosome maturation to survive inside macrophages.

### Glycolysis characterizes the M(Kp) metabolism

Metabolic reprogramming is a key aspect in the regulation of macrophage polarisation and function (35, 36). Therefore, we sought to characterize the metabolism associated with *K. pneumoniae*-induced M(Kp) polarization. Depending on the stimuli received, macrophages can switch between an aerobic metabolism, based on oxidative phosphorylation, to an anaerobic one, based on glycolysis. M1 macrophages display enhanced glycolysis whereas M2 macrophages utilize fatty acid oxidation (FAO) and mitochondrial oxidative phosphorylation (OXPHOS) to obtain energy (36). We interrogated the scRNA-seq data to reveal whether *K. pneumonaie* infection is associated with changes in the expression of metabolic genes. It has been established that metabolic changes in macrophages are associated with changes in the transcription of genes governing the different metabolic pathways (35). Dot plot analysis showed an upregulation of genes related to glycolysis in Kp52145-associated and bystander IMs including *Pfkp, Hif1a, HK3, gapdh* and *Ldha* (Fig 11A). In contrast, the expression of genes related to FAO and OXPHOS was downregulated (Fig 11A). In AMs, the glycolysis related genes *pgamI*, *HK3*, and *hif1a* were also upregulated in bystander and infected cells (Supplementary Figure 11). Taken together, these data suggest that glycolysis may characterize *K. pneumoniae*-triggered M(Kp) polarization.

**Figure 11.**
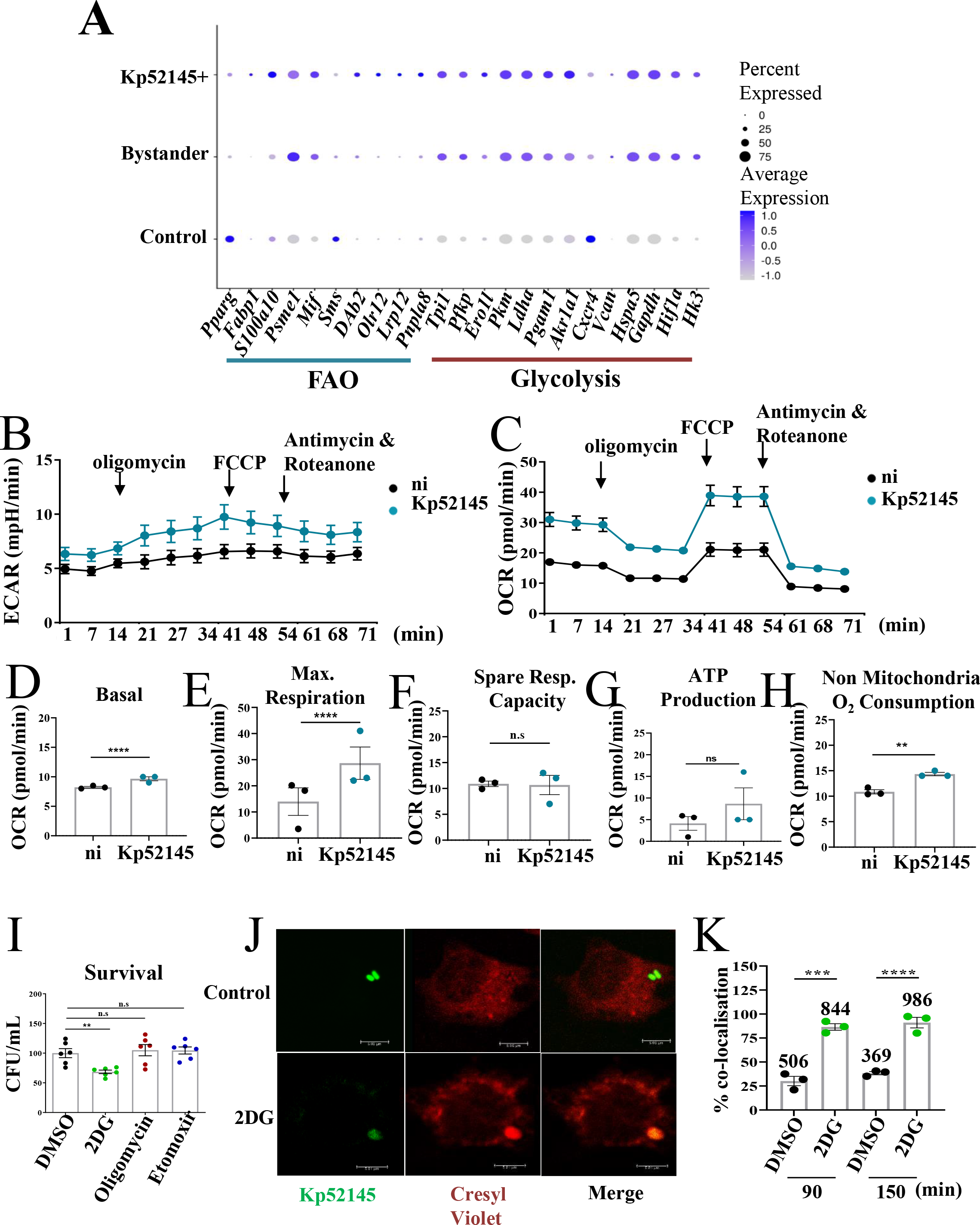
Glycolysis characterizes *K. pneumoniae*-induced M(Kp) polarisation. A. Dot Plot analysis of the expression levels of genes related to fatty acid oxidation (FAO) and glycolysis from the scRNAseq data set of PBS-infected IMs (control), and bystander and Kp52145-associated IMs. Dot size reflects percentage of cells in a cluster expressing each gene; dot colour intensity reflects expression level as indicated on legend. B. Extracellular acidification rate (ECAR, in mpH/min) of non-infected (ni) and Kp52145-infected iBMDMS (Kp52145) measured using Mito-stress test kit and the Seahorse XF analyser. When indicated oligomycin (2.5 μM), FCCP (2 μM), antimycin and roteanone (0.5 μM) were added to the cells. C. Oxygen consumption rates (OCR, in pMoles/min) of non-infected (ni) and Kp52145-infected iBMDMS (Kp52145) measured using Mito-stress test kit and the Seahorse XF analyser. When indicated oligomycin (2.5 μM), FCCP (2 μM), antimycin and roteanone (0.5 μM) were added to the cells. D. Basal respiration of non-infected (ni) and Kp52145-infected iBMDMs. E. Maximal respiration of non-infected (ni) and Kp52145-infected iBMDMs. F. Spare respiratory capacity of non-infected (ni) and Kp52145-infected iBMDMs. G. ATP production by non-infected (ni) and Kp52145-infected iBMDMs. H. Non mitochondrial O_2_ consumption by non-infected (ni) and Kp52145-infected iBMDMs. I. Kp52145 intracellular survival in wild-type iBMDMs 5 h after addition of gentamycin (30 min of contact). Results are expressed as % of survival (CFUs at 5 h versus 30 min in *stat6^-/-^* cells normalized to the results obtained in wild-type macrophages set to 100%). Cells were treated with DMSO vehicle, or 2-deoxyglucose (2DG, 3 μM), oligomycin (3 μM), etomoxir (50 μM) 2 h before infection and maintained thought. J. Immunofluorescence confocal microscopy of the colocalization of Kp52145 harbouring pFPV25.1Cm, and cresyl violet dye in wild-type macrophages treated with DMSO vehicle solution (control) or 2DG. The images were taken 90 min post infection. Images are representative of duplicate coverslips of three independent experiments. K. Percentage of Kp52145 harbouring pFPV25.1Cm co-localization with cresyl violet over a time course. Wild-type iBMDMs treated with DMSO vehicle solution (control) or 2DG. were infected; coverslips were fixed and stained at the indicated times. Values are given as mean percentage of Kp52145 co-localizing with the marker☐±☐SEM. The number of infected cells counted per time in three independent experiments are indicated in the figure. Error bars are presented as the mean ± SEM of three independent experiments in duplicate. Images are representative of three independent experiments. In panels D, E, F, G, H, and K unpaired t test was used to determine statistical significance. In all the other panels, statistical analysis were carried out using one-way ANOVA with Bonferroni contrast for multiple comparisons test. ****P ≤ 0.0001; ***P ≤ 0.001; **P 0.01; *P ≤ 0.05; ns, P > 0.05 for the indicated comparisons.

To provide experimental evidence on the metabolism linked to M(Kp), we monitored glycolysis parameters (glycolysis and glycolytic reserve) and mitochondrial function characteristics (basal mitochondrial respiration, ATP production, maximal respiration, and spare respiratory capacity) in infected macrophages by measuring the extracellular acidification rate (ECAR) and the oxygen consumption rate (OCR) using a Seahorse XFe96 analyser. Infection of wild-type macrophages with Kp52145 resulted in an increase in ECAR (Fig 11B), representing the glycolysis rate. Inhibition of the mitochondrial F_1_F_O_-ATPase with oligomycin resulted in a modest increase in ECAR (Fig 11B). The lack of increase in ECAR following the addition of the ionophore FCCP, that uncouples mitochondrial respiration by increasing H+ transport across the inner mitochondrial membrane, and rotenone and antimycin A, inhibitors of mitochondrial complex I and III, respectively, indicates that the maximal glycolytic capacity was already reached (Fig 11B). This result suggests that ATP production is mostly dependent on glycolysis in Kp52145-infected macrophages because carbon flux is coupled to glycolytic ATP production. The increase of OCR following Kp52145 infection reflected an increase in mitochondrial basal respiration (Fig 11C and Fig 11D). Addition of oligomycin triggered a decrease of cellular OCR, however the OCR was still significantly higher in Kp52145 infected cells compared to non-infected ones indicating that not all oxygen consumption is used for ATP production in *K. pneumoniae*-infected cells (Fig 11C). Subsequent addition of FCCP, which stimulates respiration, showed that the maximal respiration capacity was higher in Kp52145-infected macrophages than in non-infected ones (Fig 11C and Fig 11E). However, the spare respiratory capacity was not significantly different between Kp52145-infected macrophages and non-infected ones (Fig 11F), indicating that *K. pneumoniae* infection does not deplete the cellular energy via an increased OXPHOS. No differences were found in ATP production between non-infected and Kp52145-infected cells (Fig 11G). The fact that OCR was higher in Kp52145-infected cells as compared to non-infected ones after the addition of rotenone and antimycin A indicates that the non-mitochondrial respiration is increased in *K. pneumoniae*-infected macrophages (Fig 11H). When ECAR and OCR analysis following infection were done in the presence of the STAT6 inhibitor AS1517499, we observed a reduction in both measurements (Supplementary Figure 12A-B), connecting *Klebsiella*-induced metabolism with STAT6 activation. Altogether, these results are consistent with the model in which glycolysis is characteristic of *K. pneumoniae*-induced M(Kp) without impairment of mitochondrial bioenergetics.

To determine the effect of *K. pneumoniae*-induced metabolism on *Klebsiella*-macrophage interface, we used 2-doxyglucose (2DG) to inhibit glycolysis, and oligomycin and etomoxir to inhibit OXPHOS. Control experiments showed that the drugs did not affect the growth of Kp52145 (Supplementary Figure 13A). The drugs did not affect the adhesion of Kp52145 to macrophages (Supplementary Figure 13B). Oligomycin and etomoxir pre-treatments had no effect on the phagocytosis of Kp52145 whereas inhibition of glycolysis using 2DG resulted in an increase of phagocytosis (Supplementary Figure 13C). Time course experiments revealed that oligomycin and etomoxir pre-treatments did not impair Kp52145 intracellular survival (Fig 11I). In stark contrast, 2DG pre-treatment significantly reduced the survival of Kp52145 (Fig 11I). Moreover, 2DG pre-treatment increased the colocalization of the KCV with cresyl violet, indicating that inhibition of glycolysis results in an increased fusion of the KCV with lysosomes (Fig 11J and Fig 10K). Interestingly, oligomycin and etomoxir pre-treatments had no effect on the activation of NF-κB and Irf3 following infection whereas 2DG pre-treatment resulted in a significant decrease in NF-κB and Irf3 activation after Kp52145 infection (Supplementary Figure 13D-E). The latter result is consistent with the importance of glycolysis to mount inflammatory responses following infection (37). Altogether, these experiments demonstrate that glycolysis is associated with *K. pneumoniae*-induced M(Kp) polarization. Moreover, our results highlight the importance of host cell glycolysis for *K. pneumonaie* survival while OXPHOS seems dispensable.

### K. pneumoniae-*governed M(Kp) is dependent on the capsule polysaccharide*

We next sought to identify the *K. pneumoniae* factor(s) governing M(Kp) polarization. Given that TLR4 governs M(Kp) and that the *K. pneumoniae* capsule polysaccharide (CPS) and the LPS O-polysaccharide are recognized by TLR4 (38, 39), we asked whether these polysaccharides mediate *K. pneumoniae* induction of the M(Kp) polarization. The LPS-O-polysaccharide mutant, strain 52145-Δ*glf* (30), induced the phosphorylation of STAT6 (Fig 2A). The mutant also upregulated the levels of the M(Kp) marker CD206 to the same levels as Kp52145-infected macrophages (Fig 12B). In contrast, the *cps* mutant, strain 52145-Δ*wca_K2_* (40), did not trigger the phosphorylation of STAT6 (Fig 12C), and did not induce *arg1* (Fig 12D). Single cell experiments using flow cytometry demonstrated that the levels of Arg1, and CD206 were significantly lower in macrophages infected with the *cps* mutant than in those infected with the wild-type strain (Fig 12E and Fig12F). The opposite was found for the M1 marker MHC-II (Fig 12G). Taken together, this evidence supports the notion that *K. pneumoniae* CPS controls M(Kp) polarization.

**Figure 12.**
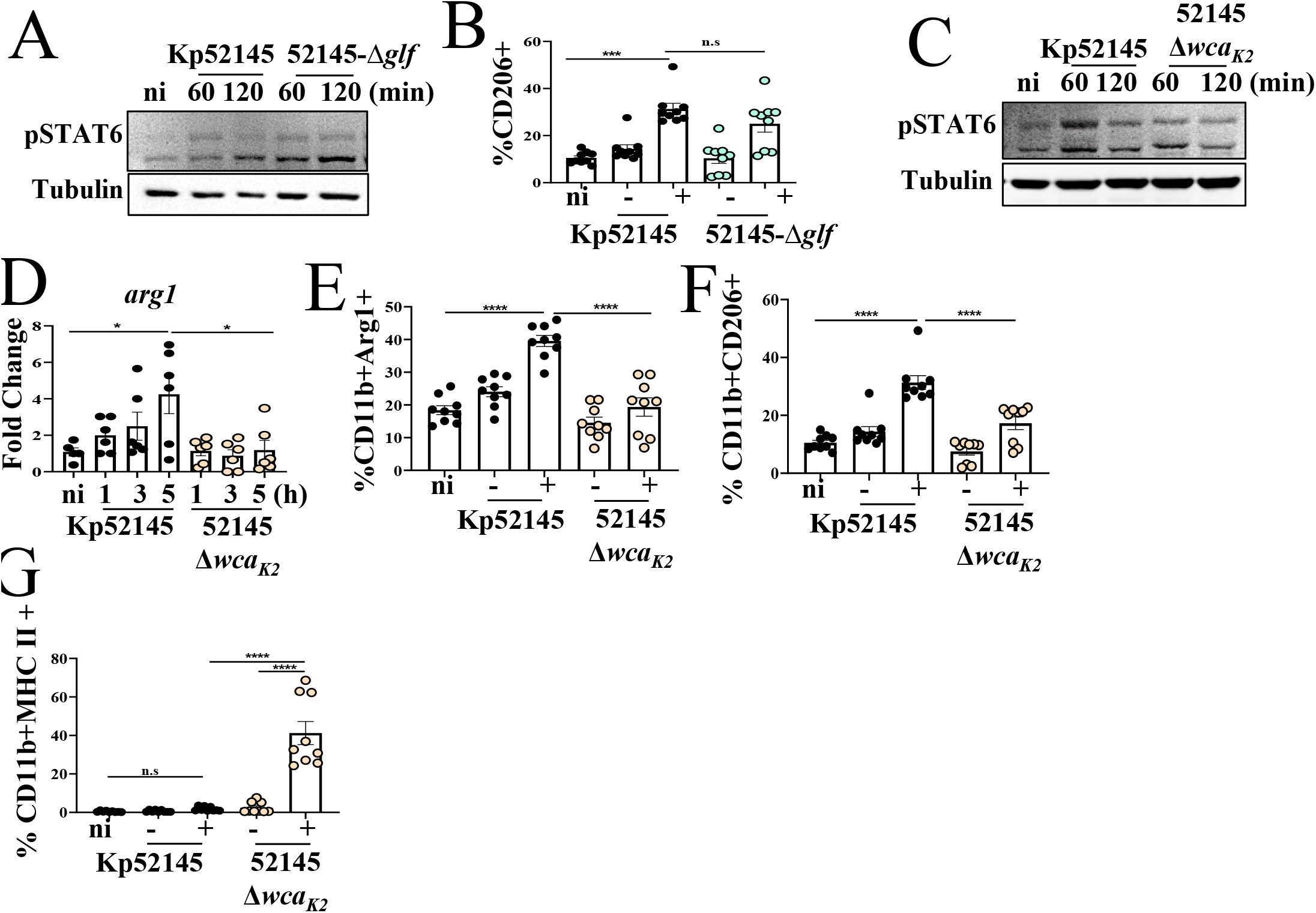
*K. pneumoniae*-governed M(Kp) is dependent on the capsule polysaccharide. A. Immunoblot analysis of phospho-STAT6 (pSTAT6) and tubulin levels in lysates from wild-type macrophages non-infected (ni), or infected with Kp52145 or the LPS O-polysaccharide mutant, strain 52145-Δ*glf*, for 60 or 120 min. B. Percentage of wild-type macrophages with and without associated Kp52145 or 52145-Δ*glf* positive for CD206 5 h post infection. Bacteria were tagged with mCherry. C. Immunoblot analysis of phospho-STAT6 (pSTAT6) and tubulin levels in lysates from wild-type macrophages non-infected (ni), or infected with Kp52145 or the CPS mutant, strain 52145-Δ*wca_K2_*, for 60 or 120 min. D. *arg1* mRNA levels were assessed by qPCR in wild-type macrophages non-infected (ni) or infected Kp52145 or the CPS mutant, strain 52145-Δ*wca_K2_*,for 1, 3 or 5 h. E. Percentage of wild-type macrophages with and without associated Kp52145 or 52145-*wca_K2_* positive for Arg1 5 h post infection. Bacteria were tagged with mCherry. F. Percentage of wild-type macrophages with and without associated Kp52145 or 52145-*wca_K2_* positive for CD206 5 h post infection. Bacteria were tagged with mCherry. G. Percentage of wild-type macrophages with and without associated Kp52145 or 52145-*wca_K2_*positive for MHCII 5 h post infection. Bacteria were tagged with mCherry. For all infections, after 1 h contact, medium replaced with medium containing gentamycin (100 µg/ml) to kill extracellular bacteria. Error bars are presented as the mean ± SEM of three independent experiments in duplicate or triplicate. Images are representative of three independent experiments. Statistical analysis were carried out using one-way ANOVA with Bonferroni contrast for multiple comparisons test. ****P ≤ 0.0001; ***P ≤ 0.001; *P ≤ 0.05; ns, P > 0.05 for the indicated comparisons.

### K. pneumoniae *induces M(Kp) polarization in human macrophages*

We next focused to extend the work performed in mice to humans to determine whether *K. pneumoniae* also triggers M(Kp) polarization in human macrophages. To address this question, we infected macrophages generated from human PBMCs from six different healthy donors. Violin plots show that Kp52145 infection increased the levels of the M(Kp) markers *arg1*, *il10*, *chi3l1, ppar*γ*, mrc1, nos2* and *isg56* (Fig 13A-G). Kp52145 also upregulated the levels of *il1rn* and *ido*, indoleamine 2,3-dyoxygenase (Fig 13H-I). *il1rn* and *ido* are two of the markers often associated with M2 polarization in human macrophages (41). These results suggest that *K. pneumonaie* also skews macrophage polarization in human macrophages towards M(Kp).

**Figure 13.**
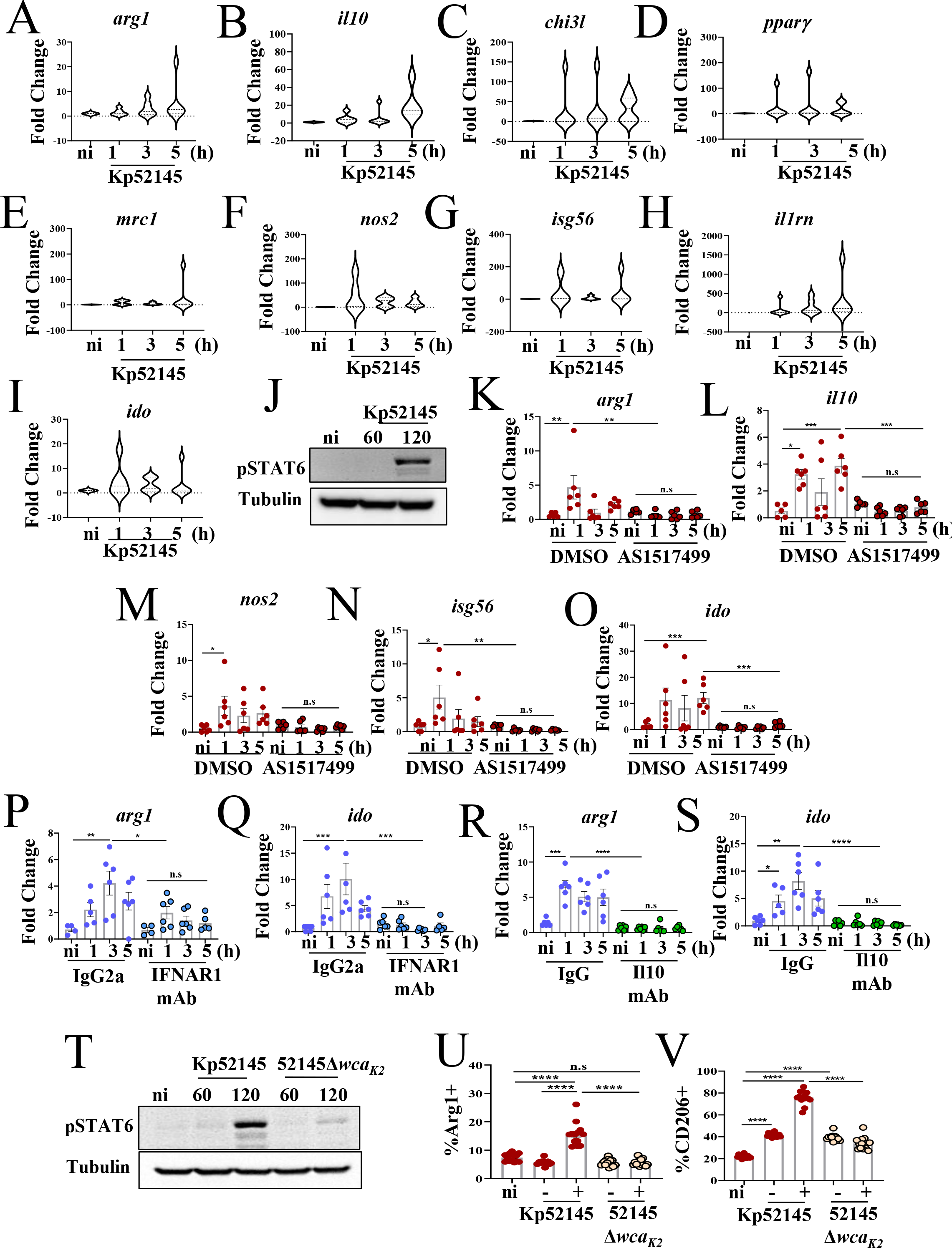
*K. pneumoniae* induces M(Kp) polarization in human macrophages. A. *arg1* mRNA levels were assessed by qPCR in hM-CSF-treated PBMCs from 6 donors non-infected (ni) or infected Kp52145 for 1, 3 or 5 h. B. *il10* mRNA levels were assessed by qPCR in M-CSF-treated PBMCs from 6 donors non-infected (ni) or infected Kp52145 for 1, 3 or 5 h. C. *chi3l* mRNA levels were assessed by qPCR in hM-CSF-treated PBMCs from 6 donors non-infected (ni) or infected Kp52145 for 1, 3 or 5 h. D. *pparg* mRNA levels were assessed by qPCR in hM-CSF-treated PBMCs from 6 donors non-infected (ni) or infected Kp52145 for 1, 3 or 5 h. E. *mrc1* mRNA levels were assessed by qPCR in hM-CSF-treated PBMCs from 6 donors non-infected (ni) or infected Kp52145 for 1, 3 or 5 h. F. *nos2* mRNA levels were assessed by qPCR in hM-CSF-treated PBMCs from 6 donors non-infected (ni) or infected Kp52145 for 1, 3 or 5 h. G. *isg56* mRNA levels were assessed by qPCR in hM-CSF-treated PBMCs from 6 donors non-infected (ni) or infected Kp52145 for 1, 3 or 5 h. H. *il1rn* mRNA levels were assessed by qPCR in hM-CSF-treated PBMCs from 6 donors non-infected (ni) or infected Kp52145 for 1, 3 or 5 h. I. *ido* mRNA levels were assessed by qPCR in hM-CSF-treated PBMCs from 6 donors non-infected (ni) or infected Kp52145 for 1, 3 or 5 h. J. Immunoblot analysis of phospho-STAT6 (pSTAT6) and tubulin levels in lysates from PMA-treated THP-1 macrophages non-infected (ni), or infected with Kp52145 for 60 or 120 min K. *arg1* mRNA levels were assessed by qPCR in PMA-treated THP-1 macrophages non-infected (ni) or infected Kp52145 for 1, 3 or 5 h and treated with the STAT6 inhibitor AS1517499 or DMSO vehicle control. L. *il10* mRNA levels were assessed by qPCR in PMA-treated THP-1 macrophages non-infected (ni) or infected Kp52145 for 1, 3 or 5 h and treated with the STAT6 inhibitor AS1517499 or DMSO vehicle control. M. *nos2* mRNA levels were assessed by qPCR in PMA-treated THP-1 macrophages non-infected (ni) or infected Kp52145 for 1, 3 or 5 h and treated with the STAT6 inhibitor AS1517499 or DMSO vehicle control. N. *isg56* mRNA levels were assessed by qPCR in PMA-treated THP-1 macrophages non-infected (ni) or infected Kp52145 for 1, 3 or 5 h and treated with the STAT6 inhibitor AS1517499 or DMSO vehicle control. O. *ido* mRNA levels were assessed by qPCR in PMA-treated THP-1 macrophages non-infected (ni) or infected Kp52145 for 1, 3 or 5 h and treated with the STAT6 inhibitor AS1517499 or DMSO vehicle control. P. *arg1* mRNA levels were assessed by qPCR in PMA-treated THP-1 macrophages non-infected (ni) or infected Kp52145 for 1, 3 or 5 h and treated with IFNAR1 blocking antibody or isotype control. Q. *ido* mRNA levels were assessed by qPCR in PMA-treated THP-1 macrophages non-infected (ni) or infected Kp52145 for 1, 3 or 5 h and treated with IFNAR1 blocking antibody or isotype control. R. *arg1* mRNA levels were assessed by qPCR in PMA-treated THP-1 macrophages non-infected (ni) or infected Kp52145 for 1, 3 or 5 h and treated with IL10 blocking antibody or isotype control. S. *ido* mRNA levels were assessed by qPCR in PMA-treated THP-1 macrophages non-infected (ni) or infected Kp52145 for 1, 3 or 5 h and treated with IL10 blocking antibody or isotype control. T. Immunoblot analysis of phospho-STAT6 (pSTAT6) and tubulin levels in lysates from PMA-treated THP-1 macrophages non-infected (ni), or infected with Kp52145 or the CPS mutant, strain 52145-*wca_K2_*, for 60 or 120 min. U. Percentage of PMA-treated THP-1 macrophages with and without associated Kp52145 or 52145-Δ*wca_K2_* positive for Arg1 5 h post infection. Bacteria were tagged with mCherry. V. Percentage of PMA-treated THP-1 macrophages with and without associated Kp52145 or 52145-Δ*wca_K2_* positive for CD206 5 h post infection. Bacteria were tagged with mCherry. For all infections, after 1 h contact, medium replaced with medium containing gentamycin (100 µg/ml) to kill extracellular bacteria. Error bars are presented as the mean ± SEM of three independent experiments in duplicate. Images are representative of three independent experiments. Statistical analysis were carried out using one-way ANOVA with Bonferroni contrast for multiple comparisons test. ****P ≤ 0.0001; ***P ≤ 0.001; **P≤ 0.01; *P ≤ 0.05; ns, P > 0.05 for the indicated comparisons.

To ascertain whether *K. pneumoniae* exploits the same molecular mechanisms in human macrophages than in mouse macrophages to induce M(Kp) polarization, we switched to PMA-differentiated THP-1 human macrophages. This cell line derived from a patient with acute monocytic leukaemia and it is used commonly to model the activation of human macrophages. Immunoblotting experiments confirmed that Kp52145 induced the phosphorylation of STAT6 (Fig 13J). RT-qPCR experiments showed that Kp52145 infection increased the levels of the M(Kp) markers *arg1*, *il10*, *nos2*, the ISG *isg56,* and *ido* in THP-1 cells in a STAT6-dependent manner because the STAT6 inhibitor AS1517499 abrogated *Klebsiella*-mediated upregulation of these M(Kp) markers (Fig 13K-O). Collectively, these results demonstrate that *K. pneumoniae* triggers M(Kp) polarization in THP-1 cells in a STAT6-dependent manner.

To determine whether *K. pneumoniae*-induced type I IFN and IL10 would also govern the induction of M(Kp) polarization in human macrophages, cells were infected in the presence of blocking antibodies against human IFNAR1 receptor, and IL10. Figure 13P-Q shows that Kp52145 did not upregulate the expression *of arg1* and *ido* when infections were done in the presence of IFNAR1 blocking antibody (Fig 13P and Fig 13Q). Similar results were obtained following IL10 suppression (Fig 13R and Fig 13S). Together, these results demonstrate that type I IFN and IL10 signalling are crucial for *K. pneumoniae* induction of M(Kp) polarization in human macrophages.

To establish whether *K. pneumoniae* CPS also controls M(Kp) polarization in human macrophages, PMA-differentiated THP-1 cells were infected with the *cps* mutant, strain *wca_K2_*. Immunoblotting experiments showed that the *cps* mutant did not induce the phosphorylation of STAT6 in THP-1 cells (Fig 13T). As anticipated, single cell analysis by flow cytometry revealed that infection with the *cps* mutant did not increase the levels of Arg1 (Fig 13U) and CD206 (Fig 13V). Taken together, these results demonstrate that *K. pneumoniae* CPS also governs M(Kp) polarization in human macrophages.

## Discussion

*K. pneumoniae* is one of the pathogens sweeping the World in the antibiotic resistant pandemic. Although, there is a wealth of knowledge on how *K. pneumoniae* develops resistance to different antibiotics, we still lack a complete understanding of what makes *K. pneumoniae* a successful pathogen. Of particular interest is to uncover whether *K. pneumoniae* has evolved strategies to overcome macrophages. These cells are an integral component of the tissue immune surveillance, response to infection and the resolution of inflammation. Therefore, pathogens such as *Klebsiella* cannot avoid innate immune responses if they are not able to overcome macrophages. Macrophages are a diverse population of immune cells of different ontogenic origins that undergo differentiation according to molecular cues provided by the microenvironment. In this work, we demonstrate that *K. pneumoniae* induces a singular polarization state, termed M(Kp), in ontogenically distinct populations of macrophages, AMs and IMs. Our findings demonstrate the central role of STAT6 in *K. pneumoniae*-governed M(Kp). Mechanistic studies revealed that *K. pneumoniae* hijacks the immune effectors type I IFN and IL10 following the activation of TLR-controlled pathways to activate STAT6-controlled M(Kp). These results illustrate how the co-evolution of *K. pneumoniae* with the immune system has resulted in the pathogen exploiting immune effectors and receptors sensing infections to thwart innate immune defences. We establish that STAT6 is necessary for *K. pneumoniae* intracellular survival whereas inhibition of STAT6 in vivo facilitates the clearance of the pathogen, revealing that STAT6-governed macrophage polarization plays an integral role in *K. pneumoniae* infection biology.

Despite four decades of extensive investigation, our knowledge of the interface between macrophages and bacterial pathogens still relies on interrogating cellular models in vitro. Therefore, we have a poor understanding of the *in vivo* molecular dynamics of the interface between pathogens and the populations of tissue resident macrophages. Our results establish that lung IMs are the main target of *K. pneumoniae*. This population of macrophages is derived from postnatal blood monocytes (42, 43), and they exhibit a distinct gene expression pattern than that of AMs in the murine lung at steady state (42, 43). IMs show marked upregulation of gene sets related to reactive oxygen species (ROS) biosynthesis, high levels of nitric oxide, iron sequestration, and inflammatory responses (42, 43). Therefore, IMs are considered to show increased microbicidal activity. Consistent with this role, infection triggers the recruitment of IMs (this work and (44, 45)) and, for example, they control *Mycobacterium tuberculosis* in vivo (46). This is in agreement with our results showing an enrichment of pathways related to sensing infections and antimicrobial defence in bystander IMs from *K. pneumoniae*-infected mice. It was not surprising that *K. pneumoniae*-infected IMs also showed some of these features including the enrichment of pathways related to cellular stress. *K. pneumoniae* also targeted AMs, although compared to IMs the number of genes differentially expressed was lower. Pathway analysis revealed an enrichment in gene networks related to TLR and NLR signalling, and inflammation in infected AMs. The fact that similar networks were found in AMs infected with *M. tuberculosis* (47) suggests that this transcription programme is part of a common AM response to bacterial pathogens. However, a distinct feature of the interaction between *K. pneumoniae* and IMs and AMs was the upregulation of networks related to an M2-like anti-inflammatory polarization state. The markers characteristic of *K. pneumoniae*-controlled macrophage polarization were Arg1, Fizz1, iNOS, CD163, *cd206*, type I IFN and IL10 signalling-regulated genes, and the decreased expression of *ppar*γ, and inflammatory markers. The fact that this macrophage polarization resembled an M2-like state but cannot be ascribed to any of the M2 subtypes (5) led us to name it as M(Kp) following international guidelines on macrophage polarization (5). To the best of our knowledge, *K. pneumoniae* is the first pathogen skewing the polarization of lung macrophages. Our results are consistent with a scenario in which rewiring macrophage polarization is a *K. pneumoniae* virulence strategy to survive in the lungs.

Interestingly, *K. pneumoniae*-induced macrophage polarization is species independent because similar findings were obtained in mouse, human and porcine macrophages (this work and (48)). The fact that there are significant differences between macrophages from different species (49) suggests that *K. pneumoniae* cellular targets should be conserved throughout evolution. Indeed, our results demonstrate that *K. pneumoniae* targets STAT6 across species to induce M(Kp) polarization (this work and (48)). This transcriptional factor arose early during the evolution of vertebrates (50) and belongs to the STAT family which is conserved through evolution from the first single cell organisms (50). Remarkably, and despite the role of STAT6 governing macrophage polarization, this transcriptional factor is not a common target of pathogens to control macrophage biology, being *Klebsiella* the first bacterial pathogen doing so.

It was intriguing to determine how *K. pneumoniae* manipulates the polarization of macrophages because it does not encode any type III or IV secretion systems known to deliver effectors into immune cells, or any of the toxins affecting cell biology. Our work demonstrates that *K. pneumoniae*-controlled STAT6 activation was dependent on type I IFN and IL10 following the activation of TLR4-governed signalling pathway. Type I IFNs, IL10, and TLR4 also appeared early during the evolution of vertebrates (51, 52). Therefore, *K. pneumoniae* has evolved to manipulate an innate axis conserved during evolution formed by TLR4-type I IFN-IL10-STAT6 to rewire macrophages. This is a previously unknown axis exploited by a human pathogen to manipulate immunity. This underappreciated anti-immune strategy is radically different from that employed by other pathogens, such as *Listeria, Salmonella, Shigella*, or *Mycobacterium,* who deliver bacterial proteins into host cells to manipulate the host. However, we believe that indeed it is an emerging theme in the infection biology of bacterial pathogens. Providing further support to this notion, TLR-controlled signalling is exploited by *S. Typhimurium* to survive in macrophages (53), and *L. monocytogenes* leverages type I IFN for intracellular survival (54, 55).

Our work sheds new light into the role of IL10 on *K. pneumoniae* infection biology. The importance of IL10 *in vivo* is marked by the fact that neutralization of the cytokine enhances the clearance of the *K. pneumoniae* (56). However, it remained an open question the exact role of IL10 in *K. pneumoniae*-host interaction beyond the well-known role of IL10 to downregulate inflammation. In this work, we establish that IL10 is essential to skew macrophage polarization and, in fact, IL10 is one of the signatures of M(Kp). In addition, our results demonstrate that IL10 is essential for the intracellular survival of *Klebsiella*. How *K. pneumoniae* induces IL10 was unknown. Our results demonstrate that *K. pneumoniae*-induced IL10 is controlled by TLR4-MyD88-CREB signalling pathway. This is a well-established pathway governing the expression of IL10 (33). The facts that CREB affects the activation of host defence responses independently of IL10, and that CREB regulates T cells (57) suggest that *K. pneumoniae* may leverage the immunomodulatory roles of CREB beyond IL10 production. Current efforts of the laboratory are devoted to investigate the role of CREB during *K. pneumoniae* infection.

Another novel finding of our work is the importance of glycolysis in *Klebsiella*-macrophage interface. Although reports indicate that OXPHOS is the metabolic signature of M2 macrophages (36), our results demonstrate that glycolysis characterises M(Kp). This finding is not totally unprecedented and, for example, the M2 tumour-associated macrophages are metabolically distinct from conventional M2 polarized subset in prioritizing usage of glycolysis as a key metabolic pathway (58). Notably, glycolysis is required for optimal survival of *K. pneumoniae* in macrophages. At present we can only speculate why glycolysis benefits *K. pneumoniae* survival. It is possible that glycolysis yields metabolites that *Klebsiella* needs as nutrients when residing in the KCV. These metabolites may also result in the regulation of the virulence factors governing the intracellular lifestyle of *Klebsiella*. Supporting this possibility, inhibition of glycolysis resulted in an increased colocalization of the KCV with lysosomes. Future studies are warranted to ascertain the effect of glycolysis and glycolysis-derived metabolites on *K. pneumonaie* virulence. Intriguingly, recent evidence suggest that this could be a general phenotype as Rosenberg and colleagues have shown that the glycolysis metabolite succinate activates *Salmonella* virulence during intracellular infection (59).

We were keen to identify the *K. pneumoniae* factor(s) governing the STAT6-mediated M(Kp) polarization. We first focused on the CPS and the LPS-O-polysaccharide because both polysaccharides are sensed by TLR4 that governs *Klebsiella*-induced activation of STAT6. Our results establish that only the CPS induced the activation of STAT6-controlled M(Kp). Importantly, the CPS is essential for *K. pneumoniae* survival in mice (pneumonia model) (60), underlining the importance of M(Kp) induction as a *K. pneumoniae* virulence trait since this process is abrogated in this mutant strain. Previous studies from the laboratory and others demonstrate the role of the CPS limiting the engulfment by phagocytes (61–64), illustrating the contribution of the CPS to *K. pneumoniae* stealth behaviour (65). However, the results of this work and our study demonstrating that the CPS activates an EGF receptor pathway to blunt inflammatory responses in epithelial cells (66) support that *K. pneumoniae* CPS is also one of the virulence factors of *K. pneumoniae* devoted to manipulate cell signalling.

*K. pneumoniae* exemplifies the global threat posed by antibiotic resistant bacteria. *K. pneumoniae*-triggered pulmonary infection has a high mortality rate reaching 50% even with antimicrobial therapy and may approach 100% for patients with alcoholism and diabetes. Not surprisingly, the World Health Organization includes *K. pneumoniae* among those pathogens for which new therapeutics are urgently needed. Absence of STAT6 resulted in macrophages consistent with M1 polarization with increased ability to clear intracellular *Klebsiella*. In vivo experiments probing a pre-clinical pneumonia mouse model demonstrated increased clearance of *K. pneumoniae* following the inhibition of STAT6. Altogether, these results strongly support that STAT6 is a target to boost human defence mechanisms against *K. pneumoniae*. Host-directed therapeutics aiming to interfere with host factors required by pathogens to counter the immune system are emerging as untapped opportunities that are urgently needed in the face of the global pandemic of antibiotic resistant infections. There is research to develop STAT6 inhibitors due to the implication of STAT6 signalling in colorectal cancer, melanoma, and allergic lung diseases. Based on our novel results, we propose that these drugs shall show a beneficial effect to treat *K. pneumoniae* infections alone or as a synergistic add-on to antibiotic treatment. Future studies shall confirm whether this is the case.

## Materials and methods

### Study approval

All animal procedures were performed in compliance with the UK Home Office and approved by the Queen’s University Belfast Animal Welfare and Ethical Review Body (AWERB). The work described in this work was carried out under project licences PPL2778 and PPL2910.

Ethical approval for the use of blood from healthy volunteers was approved by the Research Ethics Committee of the Faculty of Medicine, Health and Life Sciences of Queen’s University Belfast (approval reference MHLS 20_136). Whole blood was obtained from the Northern Ireland Blood Transfusion Service.

### Animals and infection model

C57BL/6 mice were purchased from Charles River Laboratories. *Ifnar1^-/-^* and *stat6^-/-^* animals were used to generate iBMDM cell lines in this study*. Ifnar1^-/-^* mice are maintained in Queen’s University Belfast animal facility whereas s*tat6^-/-^* animals were purchased from The Jackson Laboratory (Stock reference 005977). Mice were housed under standard laboratory conditions (12/12 h light/dark cycle with a room temperature of 21 °C and water and food available *ad libitum*). Mice used for experiments were aged between 8-12 weeks old and sex matched. For in vivo infections, bacteria in the stationary phase, were sub-cultured and grown at 37°C with agitation to reach mid log phase. Subsequently, bacteria were harvested by centrifugation (20 min, 2500 *x g*, 24°C), resuspended in PBS and adjusted to 5 x 10^4^–1 x 10^5^ colony forming units (CFU) per 30 μl as determined by plating serial dilutions on LB plates. Mice were anesthetized with isoflurane, and infected intranasally with *K. pneumoniae* in 30 μl PBS or mock-infected with PBS. In the experiments probing the STAT6 inhibitor AS1517499, mice were treated 24 h prior to infection intraperitoneally with 10mg/kg AS1517499 (AXON) or DMSO vehicle control in 200 μl volume. 6 h post-infection, mice received intranasally an additional dose of AS1517499 or DMSO (5 mg/kg in 30 μl volume). After 24 h post infection, mice were euthanized and lungs, and spleen isolated for assessment of bacterial load (CFU), or lungs processed for flow cytometry or RNA. CFUs were determined by homogenising organs in 1 mL sterile PBS and plating serial dilutions on *Salmonella-Shigella* agar plates (SIGMA). Plates were incubated overnight at 37°C before counting of colonies.

### Generation of stat6^-/-^ and ifnar1^-/-^ iBMDMs

Tibias and femurs were obtained from *ifnar1^-/-^* and *stat6^-/-^* mice (C57BL/6 background) and the bone marrow extracted under sterile conditions flushing bones with complete medium (DMEM, GlutaMAX, supplemented with 10% heat-inactivated foetal calf serum (FCS) and 1% penicillin-streptomycin). Red blood cells were lysed via incubation with ammonium-chloride-potassium (ACK) lysis buffer (A1049201; Gibco) for 5 min at room temperature. Cells were then washed in 10 ml of complete medium and passed through a 70-mm cell strainer (2236348; Fisherbrand) prior to centrifugation. Cell pellet was dislodged before plating on 20-cm petri dishes (Sarstedt) in complete medium supplemented with 20% syringe-filtered L929 supernatant, as source of macrophage colony-stimulating factor, and maintained at 37°C in a humidified atmosphere of 5% CO2. Medium was replaced with fresh medium supplemented with L929 after 1 day of culture. After 5 days, BMDMs were immortalized by exposing them to the J2 CRE virus (carrying v-myc and v-Raf/v-Mil oncogenes, kindly donated by Avinash R. Shenoy, Imperial College London) for 24 h. This step was repeated 2 days later (day 7), followed by continuous culture in DMEM supplemented with 20% (vol/vol) filtered L929 cell supernatant for 4 to 6 weeks. The presence of a homogeneous population of macrophages was assessed by flow cytometry using antibodies for CD11b (clone M1/70; catalogue number 17-0112-82; eBioscience) and CD11c (clone N418; catalogue number 48-0114-82; eBioscience).

### Culture of iBMDMs

*Wild-type, Tlr2^-/-^. Tlr4^-/-^, Tlr2/4^-/-^, Myd88^-/-^, Tram/Trif^-/-^* immortalised bone marrow-derived macrophages (iBMDMs) were obtained from BEI Resources (NIAID, NIH) (repository numbers NR-9456, NR-9457. NR-9458, NR-19975, NR-15633 and NR-9568, respectively). *Il10^-/-^* iBMDMs were described previously (67). iBMDMs were maintained in DMEN (Gibco 41965) supplemented with 10% heat-inactivated FCS, 100 U/mL penicillin, and 0.1 mg/mL streptomycin (Sigma-Aldrich) at 37 °C in a humidified 5% CO_2_ incubator. Cells were routinely tested for *Mycoplasma* contamination.

### Human PBMCs

PBMCs were isolated using density gradient media Ficoll-Paque PLUS (Cytiva) after centrifugation at 790 x g for 30 minutes. Resulting buffy coats were extracted and treated with ACK Lysis buffer (Gibco) to remove red blood cells, prior to freezing cells at a density of 1 x 10^6^ per cryovial and stored at -80°C. Cells were broken out via rapid thawing in 37°C water bath before removal of DMSO via suspension in complete medium and centrifugation. Cells were then plated at a density of 3x10^5^/well in 12-well tissue culture plates (Sarstedt) in DMEM (Gibco 41965) supplemented with 10% FCS, 100 U/ml penicillin, and 0.1 mg/mL streptomycin (Sigma−Aldrich ) supplemented with 10 ng/ml human M-CSF (Cat: 75057, Stemcell) at 37 °C in a humidified 5% CO_2_ incubator for 5 days to allow differentiation of macrophages.

### Culture and differentiation of THP-1 cells

Human THP-1 monocytes (ATCC TIB-202) were maintained in complete medium (RPMI, supplemented with 10% FCS and 5% penicillin/streptomycin) before seeding for differentiation in the presence of PMA (5 ng/mL) for 48 h prior to infection at a density of 3 x 10^5^ cells/well in 24-well plates (Sarstedt).

*Bacterial strains and culture conditions*.

Kp52145 is a clinical isolate (serotype O1:K2) previously described (15, 17). 52145-*wca_K2_* is an isogenic mutant of Kp52145 lacking the capsule polysaccharide which has been previously described (40). 52145-Δ*glf* lacks the LPS O-polysaccharide and expresses similar levels of CPS than Kp52145 (30). mCherry expressing strains were generated by electroporation of pUC18T-mini-Tn7T-Apr-mCherry plasmid (68) to Kp52145, 52145-Δ*wca_K2_* and 52145-Δ*glf*. mCherry expression was confirmed by confocal microscopy and flow cytometry. pFPV25.1Cm plasmid (61) was conjugated to Kp52145 to generate bacteria expressing GFP constitutively. Strains were grown in LB medium at 37°C on an orbital shaker (180 rpm). When appropriate, the following antibiotics were added to the growth medium at the indicated concentrations: carbenicillin, 50 μg/ml; chloramphenicol, 25 μg/ml.

### Growth curve analysis

For growth kinetics analysis, 5 μl of overnight cultures were diluted in 250 μl of LB containing DMSO vehicle control or metabolic inhibitors, and incubated at 37°C with continuous, normal shaking in a Bioscreen Automated Microbial Growth Analyzer (MTX Lab Systems, Vienna, VA, USA). Optical density (OD; 600 nm) was measured and recorded every 20☐min.

### Macrophage infections

iBMDMs were seeded into 24-well plates (1.6 x 10^5^ cells/well) for microscopy, 12-well dishes (5 x 10^5^ cells/well) for immunoblotting, and for assessing intracellular survival, and 6-wells (1 x 10^6^ cells/well) for RNA and flow cytometry in complete media and allowed to adhere overnight. THP-1 cells were differentiated with PMA in 12-well dishes (3 x 10^5^ cells/well). On the day of infection, cells were washed with 1ml PBS, and 1 ml of antibiotic free media was added to the wells. To prepare the inoculum for infections, bacteria were grown until mid-exponential phase in 5 ml LB medium, supplemented with the appropriate antibiotics when required, at 37°C on an orbital shaker (180 rpm). Bacteria were recovered by centrifugation (3,220 × *g*, 20 min, 22°C), washed once with PBS, and diluted in PBS to an OD_600_ of 1.0, which corresponds to approximately 5 × 10^8^ CFU/ml. A multiplicity of infection of 70 bacteria per cell was used in 1 ml of appropriate medium without antibiotics. To synchronise infection, plates were centrifuged at 200 × g for 5 min. After 1 h post infection, media was removed, cells washed with 1ml 1 PBS, and 1 ml of antibiotic free media supplemented with 100 µg/mL gentamicin (Sigma-Aldrich) were added to the wells. At the indicated time points, supernatants were removed, and cells processed for immunoblotting, RNA extraction, or flow cytometry.

### Blocking antibodies, cytokines stimulations, and treatment with inhibitors

For cytokine stimulation experiments, cells were treated with recombinant mouse IFNβ (1000 U/ml, Stratech) or IL-10 (250 ng/ml, catalogue 210-10 Peprotech) 3 h prior to collection for immunoblotting.

To inhibit STAT6, cells incubated with the chemical STAT6 inhibitor AS 1517499 (50 μg/ml, 919486-40-1, AXON) or DMSO as vehicle control for 2 h prior to infection and maintained throughout.

To inhibit cellular metabolism, macrophages were treated with DMSO vehicle control, or Cells were treated with DMSO vehicle, or 2-deoxyglucose (2DG, 3 μM), oligomycin (3 μ etomoxir (50 μM) 2 h before infection and maintained thought. All of them were purchased from Sigma-Aldrich.

For IL-10 neutralisation experiments, PMA differentiated THP-1 cells were incubated 2 h prior to infection and maintained throughout with either,1 μg/mL mouse monoclonal anti-human IL-10 antibody (R&D Systems. Ref: AF-217-NA), or equivalent concentrations of human IgG (Invitrogen) as control. To target IFNAR1/2, PMA differentiated THP-1 cells were treated with 10 μg/mL mouse monoclonal anti-human IFNAR2 (ThermoFisher Scientific, MMHAR-2), or human IgG (Invitrogen) as control. After 1 h of contact with bacteria, cells were washed once with PBS and complete medium without antibiotics supplemented with 100 µg/ml gentamicin (Sigma−Aldrich) and with the relevant antibodies. *RNA extraction, and RT-qPCR*

Lung tissue was homogenised using a VDI 12 tissue homogeniser (VWR) in 1 ml of TRizol reagent (Ambion) and incubated at room temperature for 5 min before storage at -80°C. Total RNA was extracted according to manufacturer’s instructions. 5 μg of total RNA were treated with recombinant DNase I (Roche Diagnostics Ltd) at 37°C for 30 min and then purified using a standard phenol–chloroform method. The RNA was precipitated with 20 μl 3 M sodium acetate (pH 5.2) and 600 μl 98% (v/v) ethanol at -20°C, washed twice in 75% (v/v) ethanol, dried and then resuspended in RNase-free H_2_O.

To purify RNA from cells, they were washed once with 1 ml PBS before lysis in TRIzol reagent (Ambion) according to the manufacturer’s instructions.

Duplicate cDNA preparations from each sample were generated from 1 μg of RNA using Moloney murine leukaemia virus (M-MLV) reverse transcriptase (Sigma−Aldrich) according to the manufacturer’s instructions. qPCR analysis of gene expression was undertaken using the KAPA SYBR FAST qPCR kit and the Rotor-Gene Q5Plex qPCR system (Qiagen). Thermal cycling conditions were as follows: 95°C for 3 min for enzyme activation, 40 cycles of denaturation at 95°C for 10 s and annealing at 60°C for 20 s. Primers used in qPCRs are listed in Supplementary Table 3. cDNA samples were tested in duplicates, and relative mRNA quantity was determined by the comparative threshold cycle (ΔΔCT) method, using hypoxanthine phosphoribosyl transferase 1 gene normalization for mouse and human samples.

### Flow cytometry

Lung tissue was homogenised using a VDI 12 tissue homogeniser (VWR) in 1 m sterile PBS and filtered through a 70 μm cell strainer (2236348 Fisherbrand) to generate single cell suspension. Suspensions were centrifuged and red cells lysed using ACK lysis buffer (A1049201, Gibco), and washed once with 1 ml PBS. Cell lines were washed once with 1 ml ice cold PBS 5 h post infection and dislodged by scraping in a further 1 m PBS. Murine samples were treated with Fc block (clone 93, Ref: 101302, BioLegend) at 4°C for 15 min at 4°C. ∼5 x 10^5^ cells per tube were washed prior to incubation with combinations of the following rat anti-mouse antibodies: against cell surface markers Ly6C APC/Cy7 (clone HK1.4, Ref: 128026), CD11b APC (Ref: 101212), CD11c Pacific Blue (Ref: 117322), SiglecF FITC (Ref: 155504, BioLegend) or SiglecF APCCy7 (Ref: 565527, Thermofisher), CD163 purified antibody (Ref: 155302) conjugated with FITC conjugation kit (Ref: ab102884, Abcam), CD206 FITC (Ref: 141704). Cells were incubated with antibodies for 15 min at 4°C prior to wash with 1 ml per tube FACS buffer (PBS with 2% FCS) prior to analysis by flow cytometry.

For experiments including the intracellular targets this protocol was extended to include 10 min incubation at room temperature in 100 μl of fixative (eBiosciences FOXp3/Transcription factor staining buffer set, ref: 00-5523-00) followed by wash in 1 ml permeabilization buffer (eBiosciences FOXp3/Transcription factor staining buffer set, ref: 00-5523-00) before incubation overnight with Arg1 FITC (Ref: 53-3697-82, Thermofisher), and iNOS PE (Ref: 12-3920-82, Thermofisher) antibodies in 100 μl permeabilization buffer at 4°C. Cells were then washed once more in 1ml permeabilization buffer before flow cytometric analysis using the Canto II (BD).

THP-1 human macrophages were treated with mouse anti-human Fc block (Ref: 422302, BioLegend), for 15 min at 4°C prior to wash in 1 ml PBS and centrifugation at 1600 rpm for 5 min and surface molecule staining with CD206 APCCy7 (BioLegend), and intracellular staining of anti-human Arginase1 FITC (Ref: 53-3697-82, Thermofisher). Cell surface markers were stained directly while intracellular staining required additional processing using a cell fixation and permeabilization kit as described above (eBiosciences FOXp3/Transcription factor staining buffer set, ref: 00-5523-00). Cell suspensions were analysed using CANTO-II analyser (BD). FlowJo V10 (Tree Star) software was used for data analysis and graphical representation.

### Cell Sorting and single-cell RNA - seq

C57BL/6 age and sex matched animals (15 per group) were infected intranasally under isoflurane anaesthesia with mCherry expressing Kp52145 or PBS. After 24 h, lungs were homogenized, pooled and red cells lysed using ACK buffer (Gibco) for 5 min at room temperature. Cells were then washed in 10 ml PBS before centrifugation at 1600 x rpm for 5 min. Supernatants were aspirated and cells were then treated with Fc block (clone 93, Ref: 101302, BioLegend, 1:1000) at 4°C for 15 min. Cells were washed again prior to incubation with combinations of the following rat anti-mouse antibodies: against cell surface markers Ly6C APC/Cy7 (clone HK1.4, Ref: 128026), CD11b APC (Ref: 101212), CD11c Pacific Blue (Ref: 117322), SiglecF FITC (Ref: 155504, BioLegend). Using the FACS Aria II (BD Biosciences) Ly6C+CD11b+CD11c+SiglecF-IMs and Ly6C+CD11b-CD11c+Siglec F+ AMs were sorted from PBS control mice. From infected animals, four separate populations were retrieved namely IMs infected with Kp52145 (Ly6C+CD11b+CD11c+SiglecF-mCherry+), bystander IMs (Ly6C+CD11b+CD11c+SiglecF-mCherry-), Kp52145-infected AMs (Ly6C+CD11b-CD11c+Siglec F+mCherry+) and bystander AMs (Ly6C+CD11b-CD11c+Siglec F+-mCherry-) populations. Cells were collected into sterile 1 x PBS and viability of all six populations was confirmed by trypan blue staining (Sigma-Aldrich) and found to be at or above 95% viable. Cells were sequenced using 10x Genomics by the Genomics Core Technology Unit, Queen’s University Belfast.

### ScRNA-seq analysis

The data has been deposited to the NCBI Gene Expression Omnibus repository with the accession number GSE184290.

Cell Ranger (version 3.0.2) was used to process raw sequencing data. Using Mkfastq, Bcl files were converted to fastq format and demultiplexed into 6 libraries corresponding to the individual samples. Reads were quantitatively aligned to *Mus musculs* reference transcriptome (mm10) with Count. Cell Ranger was used to distinguish between data from viable cells and background signal, providing filtered gene-cell count matrices for downstream analysis.

#### QC and Clustering

filtered count matrices were analysed in R (3.6.2) using Seurat 3.1.3 (69). For each library, a further QC step was performed to remove genes expressed in less than 3 cells, and cells with fewer than 200 genes or with >25% counts mapping to mitochondrial genes. The libraries were then merged resulting in a dataset of 7,462 cells with 15,547 genes. Strong overlap of replicates within the alveolar and interstitial samples indicated the absence of batch effect, therefore library integration with batch correction was not required.

After log normalisation, SingleR 1.4.0 (70) was used to predict cell type in comparison to Immunological Genome Project (71) reference data, based on gene expression correlation. Predicted cell type was used to achieve in silico purification, keeping only those cells identified as macrophages/monocytes. After this step, the combined dataset contained 5,677 cells, with the libraries ranging between 422 and 1927 cells. All other libraries were randomly down sampled to the lowest total (repeated x3 for robustness), resulting in a final dataset of six libraries at 422 cells each, totalling 2,532 cells.

The resulting data was scaled to regress out cell variation attributed to mitochondrial and ribosomal gene expression and total counts per cell. The top 2,000 variable genes were identified and used as input for principal component analysis; 50 principal components (PCs) were tested in Seurat’s JackStraw function, from which the first 38 PCs were identified as explaining more variability than expected by chance. These 38 PCs were used as input to SNN clustering (repeated at resolutions between 0.3 and 1.2) and UMAP generation.

#### Differential Expression Analysis

Marker gene detection and differential expression testing was performed in Seurat using the MAST package (version 1.12.0) (72). Genes expressed in at least 10% of cells in either group being tested, with log fold-change >0.25, and with adjusted P-value <0.05 were considered significantly differentially expressed. Differentially expressed genes were displayed as volcano plots using EnhancedVolcano 1.4.0.

#### Pseudotime Analysis

Purified data was exported to Monocle 3.0 (73) for pseudotime analysis to model expression changes in pseudotime between control and infected cells. Inferred cell trajectories were calculated on UMAP embeddings as generated in Seurat, resulting in several distinct trajectory branches. Where possible, nodes relating to control cells were considered the ‘start’ of each trajectory, with each terminal node in the Bystander and Kp52145+ groups considered distinct endpoints. Each distinct trajectory was tested separately, with correlation between gene expression and pseudotime trajectory calculated using the Moran’s I test in Monocle, filtered for significance on P- and Q-value <0.05, and Moran’s I test statistic >0.2.

#### Pathway analysis and gene networks

Pathway enrichment analysis was performed using gProfiler. For each comparison, we created a list of genes as query, selecting only those genes where adjusted P value <0.05. The analysis was performed using the g:SCS method for multiple testing correction, the Reactome database as a data source, and the default settings for the other parameters in g:Profiler. Results were exported to Cytoscape (version 3.8.2) and visualized using the AutoAnnotate application. The enrichment of transcriptional factors, their genomic binding sites and DNA binding profiles were analysed using TRANSFAC (20) included within g:Profiler. STRING database was used to predict protein-protein interactions using the clustering algorithm MCL with default parameters.

### Seahorse analysis

iBMDMs were seeded in Seahorse XF Cell Culture Microplates at a density of 2 x 10^4^ per well and allowed to adhere overnight in complete media at 37 °C in a humidified 5% CO_2_ incubator overnight. Seahorse cartridge was hydrated overnight in Seahorse XF Calibrant solution (Ref: 100840-000, Agilent) at 37°C overnight in a non-CO_2_ incubator. iBMDMs were washed once with warmed PBS and maintained in 200 μl of complete media without antibtiotics. When indicated, cells were treated with DMSO or STAT6 inhibitor AS1715499 (50 nM/mL, 919486-40-1; AXON Medchem) 2 h prior to infection and maintained throughout. iBMDMs were infected at 70:1 multiplicity of infection. The infection was synchronised by centrifugation at 200 x g for 5 min. After 1 h contact, cells were washed with PBS, and replenished with 180 μl XF base medium (Ref 103334-100, Agilent) pH 7.4 supplemented with 1 mM pyruvate, 2 mM glutamine, 10 mM glucose and 30 μg/mL gentamicin to eliminate extracellular bacteria. Metabolic activity of cells was assessed using the Seahorse Mito Stress Test Kit (Ref: 103015-100, Agilent) according to manufacturer’s instructions, and results were analysed with the Seahorse XFe96 analyser. Eight wells were used per condition in three independent experiments.

### Immunoblotting

Macrophages were infected with *K. pneumoniae* strains for time points indicated in the figure legends. Cells were then washed in 1 ml of ice-cold PBS and lysed in 90 μl Laemmli buffer (4% SDS, 10% 2-mercaptoehtanol, 20% glycerol, 0.004% bromophenol blue, 0.125 M Tris-HCl pH 6.8). The cell lysates were sonicated for 10 s at 10% amplitude (Branson Sonifier), boiled at 95°C for 5 min, and centrifuged at 12,000 x g for 1 min. 10 μl of the cell lysates were resolved by standard 10% SDS-PAGE gel and electroblotted onto 0.2 mm nitrocellulose membrane (Biotrace, VWR) using a semi-dry transfer unit (Bio-Rad). Membranes were blocked with 3% bovine serum albumin in Tris-buffered saline with Tween 20 (TBST).

Primary antibodies used were: p-Stat6 (Tyr641) (D8S9Y) (1:2000, Ref. 56554), p-STAT1 (T701) (58D6) (1:2000, Ref: 9167), p-STAT3 (Y705) (1:2000, Ref. 9145), Total STAT3 (1:2000, Ref. 12640), Arginase-1 (D4E3M) XP^®^ (1:2000, Ref: 93668), KLF4 (1:2000, Ref: 4038) all from Cell Signalling Technologies. FIZZ-1/RELM (1:1000, Ref. AF1523, R&D Systems), pCREB (1:1000, Ref: sc-7978-R, Santa Cruz BT), Tubulin (1:4000, Ref, T6074. Sigma-Aldrich). Immunoreactive bands were visualized by incubation with horseradish peroxidase-conjugated goat anti-rabbit immunoglobulins (1:5,000, Bio-Rad 170-6515) or goat anti-mouse immunoglobulins (1:5,000, Bio-Rad 170-6516). Protein bands were visualised using chemiluminescence reagents and a G:BOX Chemi XRQ chemiluminescence imager (Syngene).

To assess loading, membranes were stripped of previously used antibodies using a pH 2.2 glycine-HCl-SDS buffer and reprobed for tubulin (1:5,000, Sigma−Aldrich 6074).

### Knock-down of CREB using siRNA

Transfection of siRNAs was performed at the time of cell seeding in a 96-well plate format (2 × 10^4^ cells/well). Lipofectamine 2000 transfection reagent (Invitrogen) was used following the manufacturer’s instructions. Transfection experiments were carried out in Opti-MEM reduced serum medium (Invitrogen). siRNAs were used at a concentration of 20 nM, and experiments were carried out 48 h after transfection. The knockdown efficiency of the siRNA targeting murine CREB1 (Dharmacon, Ref: SO-2681745G) was quantified using RT-qPCR using conditions described above.

### Adhesion, phagocytosis, and intracellular survival assay

Intracellular survival experiments were carried out as previously described with minor modifications (28). Briefly, macrophages were seeded in 12-well tissue culture and allowed to adhere overnight at 37 °C in a humidified 5% CO_2_ incubator. Cells were infected at an MOI of 70:1 in a final volume of 500 μl antibiotic free complete medium. To synchronize infection, plates were centrifuged at 200 × *g* for 5 minutes. Plates were incubated at 37 °C in a humidified 5% CO_2_ atmosphere. After 1 hour of contact, cells were washed twice with PBS and incubated for additional 30 minutes with 500 μl complete medium containing gentamicin (100 μg /ml) to eliminate extracellular bacteria. For time course experiments, after 90 min, cells were washed with PBS and incubated with 500 μL complete medium containing gentamicin (5μg/ml). To determine intracellular bacterial load, cells were washed twice with prewarmed PBS and lysed with 300 μl of 0.05% saponin (Sigma-Aldrich) in PBS for 5 minutes at 37 °C. Adhesion was determined at 60 minutes post infection, phagocytosis at 90 minutes post infections, and survival at 330 minutes post infection. Serial dilutions were plated on LB to quantify the number of intracellular bacteria. All experiments were carried out in duplicate on at least three independent occasions.

### Confocal microscopy

iBMDMs were seeded in 24-well plates on 13 mm glass coverslips (VWR). Infections were performed at a multiplicity of infection of 100 bacteria per iBMDM in a final volume of 500 µl complete medium. To synchronize infection, cells were centrifuged (200 x *g* for 5 min). After 60 minutes contact, cells were washed with sterile PBS and incubated in 500 µL complete medium containing gentamicin (100 µg/ml). Lysosomes were stained using cresyl violet (Ostrowski et al., 2016). 15 minutes before the end of the experiment, cresyl violet (Sigma-Aldrich) was added to the coverslips to achieve 5 µM final concentration. Coverslips were washed in PBS and fixed with 4% paraformaldehyde (PFA) (Sigma-Aldrich) for 20 min at room temperature. Coverslips were mounted with ProLong Gold antifade mountant (Invitrogen). Coverslips were visualised on a Leica SP5 Confocal microscope with the appropriate filter sets. Experiments were carried out in duplicate on three independent occasions.

### Assessment of NF-κB and Irf3 activation

To quantify the activation of the NF-κB signalling pathway, we probed Raw-Blue cells (InvivoGen) derived from Raw 264.7 macrophages containing a chromosomal integration of a secreted embryonic alkaline phosphatase (SEAP) reported inducible by NF-κB and AP-1. Cells were maintained in DMEM supplemented with 10% heat-inactivated FBS, 50 U/ml penicillin, 50 μg/ml streptomycin. 200 μg/ml Zeocin were added in alternate passages to maintain the reporter plasmid. Cells were seeded at a density of ∼50,400 cells per well in a 96-well plate with complete medium without antibiotics, incubated overnight at 37 °C in a humidified 5% CO_2_ incubator. Cells were then infected at a multiplicity of infection of 100:1 in a final volume of 190 μl antibiotic free complete medium. To synchronize infection, plates were centrifuged at 200 × *g* for 5 min. Plates were incubated at 37 °C in a humidified 5% CO_2_ atmosphere. After 1 h of contact, cells were washed twice with PBS and incubated for 5 h with 180 μl complete medium containing gentamicin (100 μg/ml) to eliminate extracellular bacteria. Supernatants were then transferred to a new 96-well plate, and QUANTI-Blue (InvivoGen) was added in a 1:9c measured at 620nm (POLARstar Omega).

To quantify the activation of Irf3 signalling pathway, Raw-Lucia ISG cells were probed. These cells are derived from Raw 264.7 macrophages after stable integration of an interferon regulatory factor (irf)-inducible Lucia luciferase reporter construct. Cells were cultured in DMEM supplemented with 10% heat-inactivated FBS, 50 U/ml penicillin, 50 μg/ml streptomycin. 200 μg/ml Zeocin were added in alternate passages to maintain the reporter plasmid. Cells were seeded and infected as described for Raw-Blue cells. At the end of the experiment, 20 µl of media were transferred to flat, white bottom luminometer plates (LUMITRAC, Greiner), 50 µl QUANTI-Luc (Invivogen) were added to the wells, and luminescence was read immediately (Promega GloMax).

### Statistics

Statistical analyses were performed with Prism (version 9.1.2) software (GraphPad Software) using one-way analysis of variance (ANOVA) with Bonferroni correction for multiple comparisons, or unpaired two-tailed Student’s t test. Error bars indicate standard errors of the means (SEM). Statistical significance is indicated in figures as follows: ns, not significant (P >0.05); *, P < 0.05; **, P < 0.01; ***, P < 0.001; and ****, P < 0.0001.

### Authors’ contribution

Conceptualization, J.A.B., A.K., and A.D.; Investigation, A.D., O.C., B.M., J. sP., R. CG., G.M., R.L., D.S., and A.K. Resources, J. sP..; Funding acquisition, J.A.B. and A.K.; Writing original draft, J.A.B., A.D., A.K. D.S.; Writing-Review and Editing A.D., D.S., J.A.B. and A.K. Supervision, J.A.B., and A.K.

## Acknowledgements

We thank the members of the J.A.B. laboratory for their thoughtful discussions and support with this project. This work was supported by Biotechnology and Biological Sciences Research Council (BBSRC, BB/P006078/1, BB/P020194/1), and Medical Research Council (MRC, MR/V032496/1) funds to J.A.B.

## Supplementary figure legends

**Figure S1. Flow cytometric analysis of myeloid subsets in lungs of *K. pneumoniae*-infected mice.**

Gating strategy utilised to identify CD11b+CD11c-SiglecF-monocytes (MN), CD11b+CD11c+SiglecF-interstitial macrophages (IMs) and CD11b-CD11c+SiglecF+ tissue resident alveolar macrophages (AMs). Kp52145 was tageed with mCherry to allow the identification of macrophages with associated bacteria and bystander cells. This gating strategy was utilised also for FACS sorting of these populations for scRNAseq.

**Figure S2. Network enrichment mapping of significantly downreulated genes of IMs.**

Analysis was performed using the g:SCS method for multiple testing correction (gProflier), the Reactome database as a data source, and the default settings for the other parameters in gProflier. Results were exported to Cytoscape and visualized using the AutoAnnotate plug.

A. Kp52145-infected IMs.

B. Bystander IMs.

**Figure S3. Network enrichment mapping of significantly upregulated genes of infected AMs.**

Analysis was performed using the g:SCS method for multiple testing correction (gProflier), the Reactome database as a data source, and the default settings for the other parameters in gProflier. Results were exported to Cytoscape and visualized using the AutoAnnotate plug.

**Figure S4. Trajectory analysis of AMs from PBS-mock infected mice, and *K. pneumoniae* infected mice.**

Monocle analysis did not reveal any clear trajectory in AMs from non-infected, or infected mice.

**Figure S5. Network enrichment mapping of significantly upregulated pathways within modules 1 and 7 of *K. pneumoniae*-infected IMs.**

Analysis was performed using the g:SCS method for multiple testing correction (gProflier), the Reactome database as a data source, and the default settings for the other parameters in gProflier. Results were exported to Cytoscape and visualized using the AutoAnnotate plug.

**Figure S6**. Expression of macrophage polarisation markers in bystander and *K. pneumonia*e-infected AMs.

Dot Plot analysis of the expression levels of genes related to M1 and M2 polarisation from the scRNAseq data set of PBS-infected AMs cells (control), and bystander and Kp52145-associated AM. Dot size reflects percentage of cells in a cluster expressing each gene; dot colour intensity reflects expression level as indicated on legend.

**Figure S7. *K. pneumoniae* induces M(Kp) polarisation in immortalized BMDMs.**

A. *arg1* mRNA levels were assessed by qPCR in wild-type iBMDMs non-infected (ni) or infected with Kp52145 for 1, 3 or 5 h. Immunoblot analysis of Arg1 and tubulin levels in lysates from non-infected (ni) and infected wild-type cells with Kp52145 for 60 or 120 min.

B. *fizz1* mRNA levels were assessed by qPCR in wild-type iBMDMs non-infected (ni) or infected with Kp52145 for 1, 3 or 5 h. Immunoblot analysis of Fizz1 and tubulin levels in lysates from non-infected (ni) and infected wild-type cells with Kp52145 for 60 or 120 min.

C. *pparg* mRNA levels were assessed by qPCR in wild-type iBMDMs non-infected (ni) or infected with Kp52145 for 1, 3 or 5 h.

D. *nos2* mRNA levels were assessed by qPCR in wild-type iBMDMs non-infected (ni) or infected with Kp52145 for 1, 3 or 5 h.

E. *il12* mRNA levels were assessed by qPCR in wild-type iBMDMs non-infected (ni) or infected with Kp52145 for 1, 3 or 5 h.

F. *il6* mRNA levels were assessed by qPCR in wild-type iBMDMs non-infected (ni) or infected with Kp52145 for 1, 3 or 5 h.

G. *tnfa* mRNA levels were assessed by qPCR in wild-type iBMDMs non-infected (ni) or infected with Kp52145 for 1, 3 or 5 h.

H. *il10* mRNA levels were assessed by qPCR in wild-type iBMDMs non-infected (ni) or infected with Kp52145 for 1, 3 or 5 h.

I. Immunoblot analysis of phospho-STAT3 (pSTAT3) and tubulin levels in lysates from non-infected (ni) and infected wild-type cells with Kp52145 for 60 or 120 min.

For all infections, after 1 h contact, medium replaced with medium containing gentamycin (100 µg/ml) to kill extracellular bacteria. Error bars are presented as the mean ± SEM of three independent experiments in duplicate. Images are representative of three independent experiments. Statistical analysis were carried out using one-way ANOVA with Bonferroni contrast for multiple comparisons test. ****P ≤ 0.0001; **P 0.01; *P ≤ 0.05; ns, P > 0.05 for the indicated comparisons.

**Figure S8. *K. pneumoniae*-induced M(Kp) polarisation is STAT6-dependent.**

A. *arg1* mRNA levels were assessed by qPCR in wild-type iBMDMs non-infected (ni) or infected with Kp52145 for 1, 3 or 5 h treated with STAT6 inhibitor AS1517499 or vehicle control.

B. *il10* mRNA levels were assessed by qPCR in wild-type iBMDMs non-infected (ni) or infected with Kp52145 for 1, 3 or 5 h treated with STAT6 inhibitor AS1517499 or vehicle control.

C. *fizz1* mRNA levels were assessed by qPCR in wild-type iBMDMs non-infected (ni) or infected with Kp52145 for 1, 3 or 5 h treated with STAT6 inhibitor AS1517499 or vehicle control.

D. *nos2* mRNA levels were assessed by qPCR in wild-type iBMDMs non-infected (ni) or infected with Kp52145 for 1, 3 or 5 h treated with STAT6 inhibitor AS1517499 or vehicle control.

E. *il12* mRNA levels were assessed by qPCR in wild-type iBMDMs non-infected (ni) or infected with Kp52145 for 1, 3 or 5 h treated with STAT6 inhibitor AS1517499 or vehicle control.

F. *isg15* mRNA levels were assessed by qPCR in wild-type iBMDMs non-infected (ni) or infected with Kp52145 for 1, 3 or 5 h treated with STAT6 inhibitor AS1517499 or vehicle control.

For all infections, after 1 h contact, medium replaced with medium containing gentamycin (100 µg/ml) to kill extracellular bacteria. Error bars are presented as the mean ± SEM of three independent experiments in duplicate. Statistical analysis were carried out using one-way ANOVA with Bonferroni contrast for multiple comparisons test. ****P ≤ 0.0001; ***P ≤ 0.001; **P≤ 0.01; *P ≤ 0.05; ns, P > 0.05 for the indicated comparisons.

**Figure S9. Neither IL10 nor type I IFN activate STAT6.**

A. Immunoblot analysis of phospho-STAT6 (pSTAT6) and tubulin levels in lysates from non-infected (ni) and cells treated with recombinant IL10 (250 μg/ml), recombinant IFNβ (1000 units/ml) or both for 3 h.

Immunoblot analysis of Arg1 and tubulin levels in lysates from non-infected (ni) and cells treated with recombinant IL10 (250 μg/ml), recombinant IFNβ (1000 units/ml) or both for 3 h.

Images are representative of three independent experiments.

**Figure S10. Knockdown efficiency of CREB in iBMDMs using siRNA.**

Efficiency of transfection of CREB siRNA (siCREB) in wild-type macrophages. mRNA levels were assessed 16 h post transfection as fold change against control non-silencing agents AllStars (siAS). Values are presented as the mean ± SEM of three independent experiments measured in duplicate. **P ≤ 0.01 unpaired t test.

**Figure S11. STAT6 governed M(Kp) metabolism.**

A. Extracellular acidification rate (ECAR, in mpH/min) of non-infected (ni) and Kp52145-infected iBMDMS (Kp52145) treated with DMSO vehicle control or the STAT6 inhibitor AS1517499 measured using Mito-stress test kit and the Seahorse XF analyser. When indicated oligomycin (2.5 μM), FCCP (2 μM), antimycin and roteanone (0.5 μM) were added to the cells.

B. Oxygen consumption rates (OCR, in pMoles/min) of non-infected (ni) and Kp52145-infected iBMDMS (Kp52145) treated with DMSO vehicle control or the STAT6 inhibitor AS1517499 measured using Mito-stress test kit and the Seahorse XF analyser. When indicated oligomycin (2.5 μM), FCCP (2 μM), antimycin and roteanone (0.5 μM) were added to the cells.

For each measurement, the standard error of the mean (SEM) of eight individual wells in three independent experiments is presented.

**Figure S12. Upregulation of genes related to glycolysis in AMs following infection.**

Dot Plot analysis of the expression levels of genes related to fatty acid oxidation (FAO) and glycolysis from the scRNAseq data set of PBS-infected AMs (control), and bystander and Kp52145-associated AMs. Dot size reflects percentage of cells in a cluster expressing each gene; dot colour intensity reflects expression level as indicated on legend.

**Figure S13. Effect of inhibition of host metabolism on *K. pneumoniae*-macrophage interface.**

A. Growth kinetics of Kp52145 cultures in LB containing the glycolysis inhibitor 2-doxyglucose (2DG, 3 μM), the FAO inhibitors oligomycin (1 μM) or etomoxir (50 μM), or DMSO vehicle control. Values are presented as the mean ± SEM of three independent experiments measured in triplicate

B. Adhesion of Kp52145 to iBMDMs treated with DMSO vehicle control or the glycolysis inhibitor 2-doxyglucose (2DG, 3 μM), the FAO inhibitors oligomycin (1 μM) or etomoxir (50 μM). Inhibitors were added 2 h before and maintained throught.

C. Phagocytosis of Kp52145 by iBMDMs treated with DMSO vehicle control or the glycolysis inhibitor 2-doxyglucose (2DG, 3 μM), the FAO inhibitors oligomycin (1 μM) or etomoxir (50 μM). Inhibitors were added 2 h before and maintained throughout.

D. Activation of NF-κB signalling measured by quantifying SEAP secreted to the supernatants of Raw-Blue cells (InvivoGen) following infection with Kp52145. Cells treated with DMSO, 2DG (3 μM), oligomycin (1 μM) or etomoxir (50 μM) 2 h before infection and maintained throughout experiment.

E. Activation of Irf3 measured by quantifying secreted Lucia luciferase to the supernatants of Raw-Lucia ISG cells (InvivoGen) following infection with Kp52145. Cells treated with DMSO, 2DG (3 μM), oligomycin (1 μM) or etomoxir (50 μM) 2 h before infection and maintained throughout experiment.

For all infections, after 1 h contact, medium replaced with medium containing gentamycin (100 µg/ml) to kill extracellular bacteria. Error bars are presented as the mean ± SEM of three independent experiments in duplicate. Statistical analysis were carried out using one-way ANOVA with Bonferroni contrast for multiple comparisons test. ****P ≤ 0.0001; ns, P > 0.05 for the indicated comparisons.

