## Supplementary Figures for "In vivo single cell transcriptomics reveals *Klebsiella pneumoniae* rewiring of lung macrophages to promote infection"

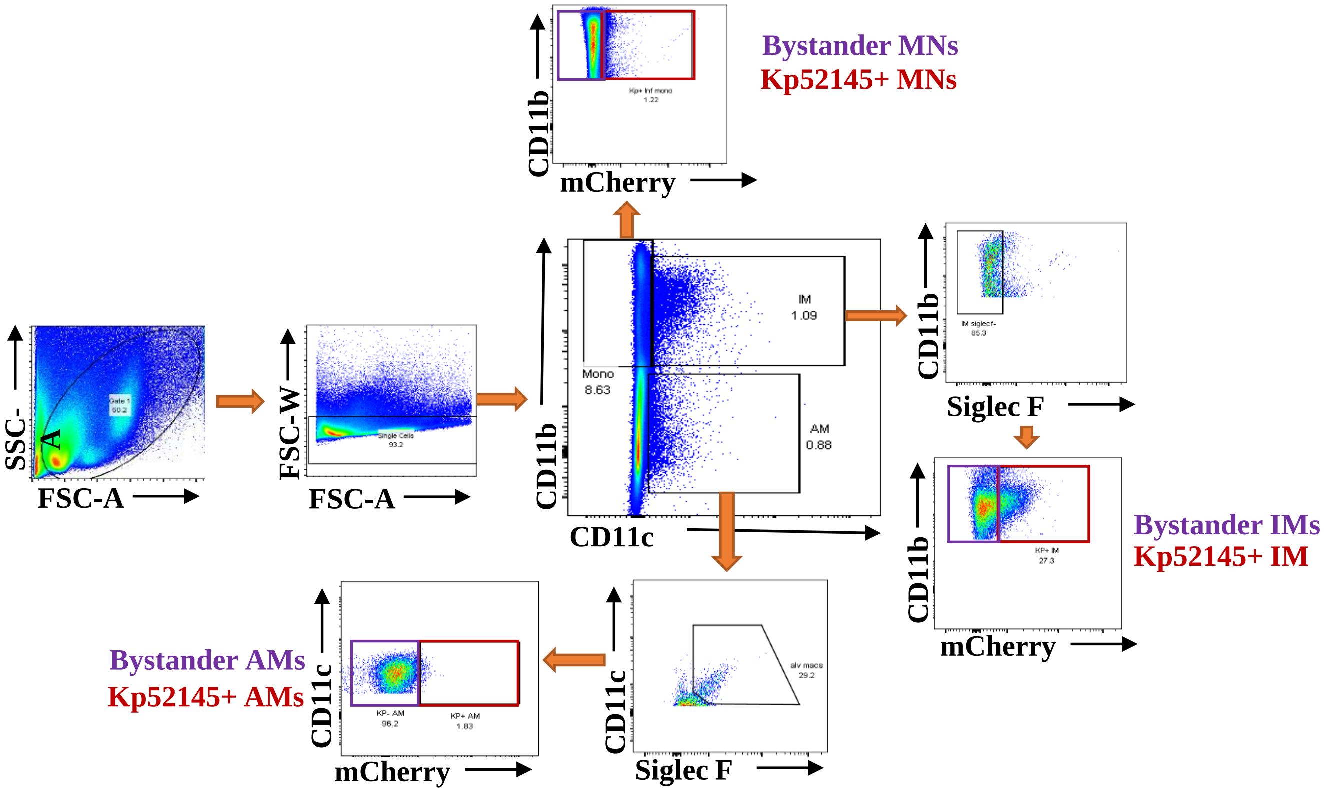
SSC-A
FSC-A
FSC-W
FSC-A
CD11b
CD11c
Mono
8.63
IM
1.09
AM
0.88
CD11b
mCherry
Bystander MNs
Kp52145+ MNs
CD11b
Siglec F
CD11b
mCherry
Bystander IMs
Kp52145+ IM
CD11c
Siglec F
CD11c
mCherry
Bystander AMs
Kp52145+ AMs

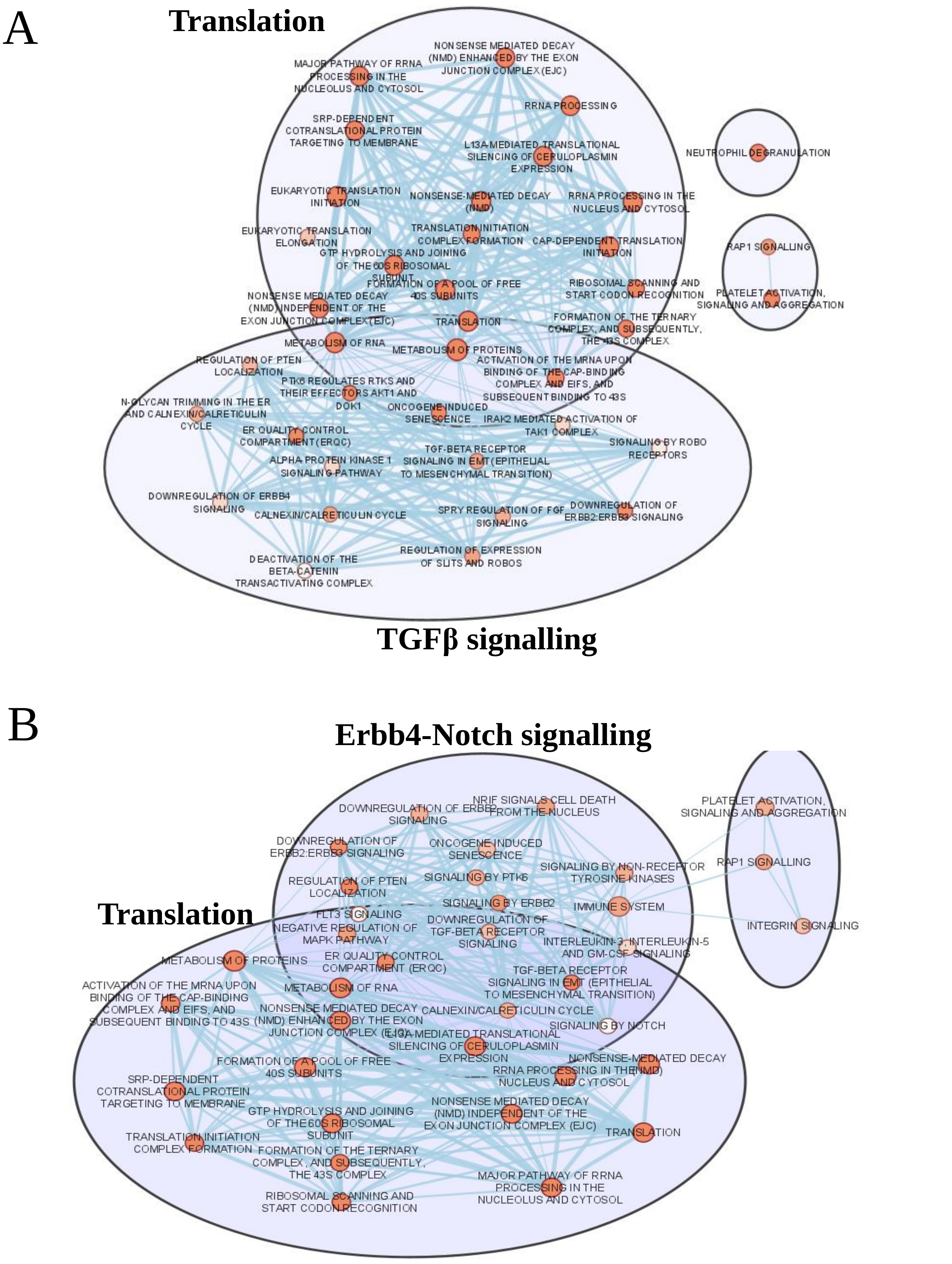
A
Translation
NONSENSE MEDIATED DECAY (NMD) ENHANCED BY THE EXON JUNCTION COMPLEX (EJC)
MAJOR PATHWAY OF RRNA PROCESSING IN THE NUCLEOLUS AND CYTOSOL
RRNA PROCESSING
SRP-DEPENDENT COTRANSLATIONAL PROTEIN TARGETING TO MEMBRANE
L13A-MEDIATED TRANSLATIONAL SILENCING OF CERULOPLASMIN EXPRESSION
NEUTROPHIL DEGRANULATION
EUKARYOTIC TRANSLATION INITIATION
NONSENSE-MEDIATED DECAY (NMD)
RRNA PROCESSING IN THE NUCLEUS AND CYTOSOL
EUKARYOTIC TRANSLATION ELONGATION
TRANSLATION INITIATION COMPLEX FORMATION
CAP-DEPENDENT TRANSLATION INITIATION
RAP1 SIGNALLING
GTP HYDROLYSIS AND JOINING OF THE 60S RIBOSOMAL SUBUNIT
FORMATION OF A POOL OF FREE 40S SUBUNITS
RIBOSOMAL SCANNING AND START CODON RECOGNITION
PLATELET ACTIVATION, SIGNALING AND AGGREGATION
NONSENSE MEDIATED DECAY (NMD) INDEPENDENT OF THE EXON JUNCTION COMPLEX (EJC)
TRANSLATION
FORMATION OF THE TERNARY COMPLEX, AND SUBSEQUENTLY, THE 43S COMPLEX
METABOLISM OF RNA
METABOLISM OF PROTEINS
REGULATION OF PTEN LOCALIZATION
ACTIVATION OF THE MRNA UPON BINDING OF THE CAP-BINDING COMPLEX AND EIFS, AND SUBSEQUENT BINDING TO 43S
PTK6 REGULATES RTKS AND THEIR EFFECTORS AKT1 AND DOK1
N-GLYCAN TRIMMING IN THE ER AND CALNEXIN/CALRETICULIN CYCLE
ONCOGENE INDUCED SENESCENCE
IRAK2 MEDIATED ACTIVATION OF TAK1 COMPLEX
ER QUALITY CONTROL COMPARTMENT (ERQC)
SIGNALING BY ROBO RECEPTORS
TGF-BETA RECEPTOR SIGNALING IN EMT (EPITHELIAL TO MESENCHYMAL TRANSITION)
ALPHA-PROTEIN KINASE 1 SIGNALING PATHWAY
DOWNREGULATION OF ERBB4 SIGNALING
CALNEXIN/CALRETICULIN CYCLE
SPRY REGULATION OF FGF SIGNALING
DOWNREGULATION OF ERBB2:ERBB3 SIGNALING
REGULATION OF EXPRESSION OF SLITS AND ROBOS
DEACTIVATION OF THE BETA-CATENIN TRANSACTIVATING COMPLEX
TGFβ signalling
B
Erbb4-Notch signalling
DOWNREGULATION OF ERBB2 SIGNALING
NRIF SIGNALS CELL DEATH FROM THE NUCLEUS
PLATELET ACTIVATION, SIGNALING AND AGGREGATION
DOWNREGULATION OF ERBB2:ERBB3 SIGNALING
ONCOGENE INDUCED SENESCENCE
SIGNALING BY NON-RECEPTOR TYROSINE KINASES
RAP1 SIGNALLING
REGULATION OF PTEN LOCALIZATION
SIGNALING BY PTK6
Translation
SIGNALING BY ERBB2
IMMUNE SYSTEM
FLT3 SIGNALING
NEGATIVE REGULATION OF MAPK PATHWAY
DOWNREGULATION OF TGF-BETA RECEPTOR SIGNALING
INTEGRIN SIGNALING
INTERLEUKIN-3, INTERLEUKIN-5 AND GM-CSF SIGNALING
METABOLISM OF PROTEINS
ER QUALITY CONTROL COMPARTMENT (ERQC)
TGF-BETA RECEPTOR SIGNALING IN EMT (EPITHELIAL TO MESENCHYMAL TRANSITION)
ACTIVATION OF THE MRNA UPON BINDING OF THE CAP-BINDING COMPLEX AND EIFS, AND SUBSEQUENT BINDING TO 43S
METABOLISM OF RNA
NONSENSE MEDIATED DECAY (NMD) ENHANCED BY THE EXON JUNCTION COMPLEX (EJC)
CALNEXIN/CALRETICULIN CYCLE
SIGNALING BY NOTCH
L13A-MEDIATED TRANSLATIONAL SILENCING OF CERULOPLASMIN EXPRESSION
FORMATION OF A POOL OF FREE 40S SUBUNITS
NONSENSE-MEDIATED DECAY (NMD)
RRNA PROCESSING IN THE NUCLEUS AND CYTOSOL
SRP-DEPENDENT COTRANSLATIONAL PROTEIN TARGETING TO MEMBRANE
GTP HYDROLYSIS AND JOINING OF THE 60S RIBOSOMAL SUBUNIT
NONSENSE MEDIATED DECAY (NMD) INDEPENDENT OF THE EXON JUNCTION COMPLEX (EJC)
TRANSLATION
TRANSLATION INITIATION COMPLEX FORMATION
FORMATION OF THE TERNARY COMPLEX, AND SUBSEQUENTLY, THE 43S COMPLEX
MAJOR PATHWAY OF RRNA PROCESSING IN THE NUCLEOLUS AND CYTOSOL
RIBOSOMAL SCANNING AND START CODON RECOGNITION

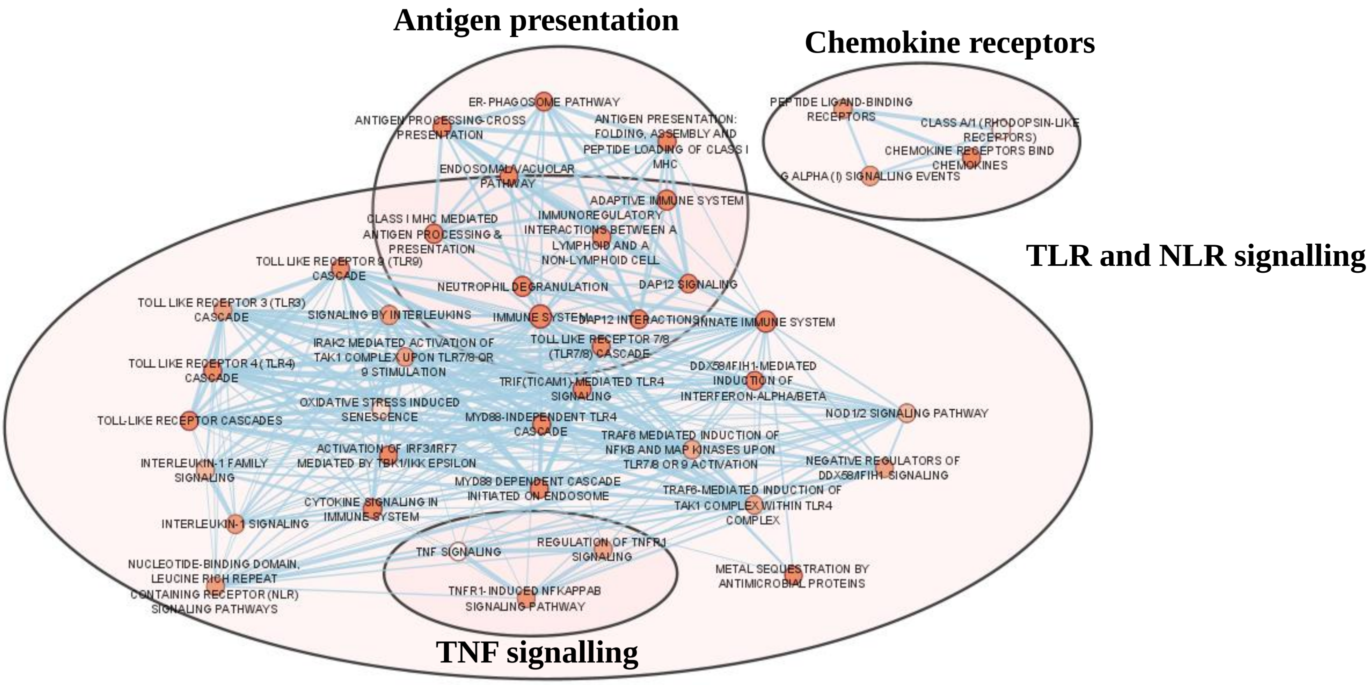

Antigen presentation
Chemokine receptors
TLR and NLR signalling
TNF signalling
ER-PHAGOSOME PATHWAY
ANTIGEN PROCESSING-CROSS PRESENTATION
ANTIGEN PRESENTATION: FOLDING, ASSEMBLY AND PEPTIDE LOADING OF CLASS I MHC
ENDOSOMAL/VACUOLAR PATHWAY
ADAPTIVE IMMUNE SYSTEM
CLASS I MHC MEDIATED ANTIGEN PROCESSING & PRESENTATION
IMMUNOREGULATORY INTERACTIONS BETWEEN A LYMPHOID AND A NON-LYMPHOID CELL
PEPTIDE LIGAND-BINDING RECEPTORS
CLASS A/1 (RHODOPSIN-LIKE RECEPTORS)
CHEMOKINE RECEPTORS BIND CHEMOKINES
G ALPHA (I) SIGNALLING EVENTS
TOLL LIKE RECEPTOR 9 (TLR9) CASCADE
NEUTROPHIL DEGRANULATION
DAP12 SIGNALING
TOLL LIKE RECEPTOR 3 (TLR3) CASCADE
SIGNALING BY INTERLEUKINS
IMMUNE SYSTEM
DAP12 INTERACTIONS
INNATE IMMUNE SYSTEM
IRAK2 MEDIATED ACTIVATION OF TAK1 COMPLEX UPON TLR7/8 OR 9 STIMULATION
TOLL LIKE RECEPTOR 7/8 (TLR7/8) CASCADE
TOLL LIKE RECEPTOR 4 (TLR4) CASCADE
DDX58/IFIH1-MEDIATED INDUCTION OF INTERFERON-ALPHA/BETA
TRIF(TICAM1)-MEDIATED TLR4 SIGNALING
OXIDATIVE STRESS INDUCED SENESCENCE
NOD1/2 SIGNALING PATHWAY
TOLL-LIKE RECEPTOR CASCADES
MYD88-INDEPENDENT TLR4 CASCADE
TRAF6 MEDIATED INDUCTION OF NFKB AND MAP KINASES UPON TLR7/8 OR 9 ACTIVATION
INTERLEUKIN-1 FAMILY SIGNALING
ACTIVATION OF IRF3/IRF7 MEDIATED BY TBK1/IKK EPSILON
NEGATIVE REGULATORS OF DDX58/IFIH1 SIGNALING
MYD88 DEPENDENT CASCADE INITIATED ON ENDOSOME
TRAF6-MEDIATED INDUCTION OF TAK1 COMPLEX WITHIN TLR4 COMPLEX
CYTOKINE SIGNALING IN IMMUNE SYSTEM
INTERLEUKIN-1 SIGNALING
REGULATION OF TNFR1 SIGNALING
TNF SIGNALING
NUCLEOTIDE-BINDING DOMAIN, LEUCINE RICH REPEAT CONTAINING RECEPTOR (NLR) SIGNALING PATHWAYS
METAL SEQUESTRATION BY ANTIMICROBIAL PROTEINS
TNFR1-INDUCED NFKAPPAB SIGNALING PATHWAY

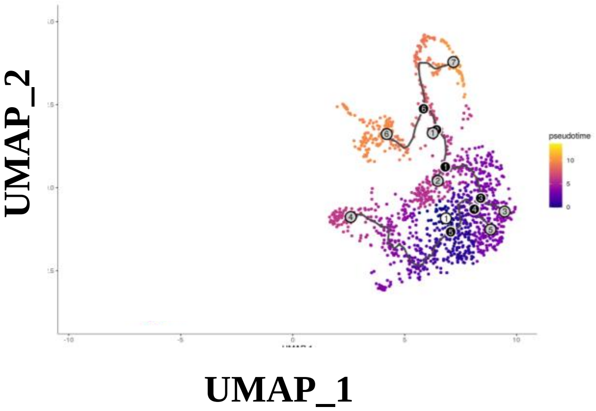
UMAP_2
pseudotime
UMAP_1

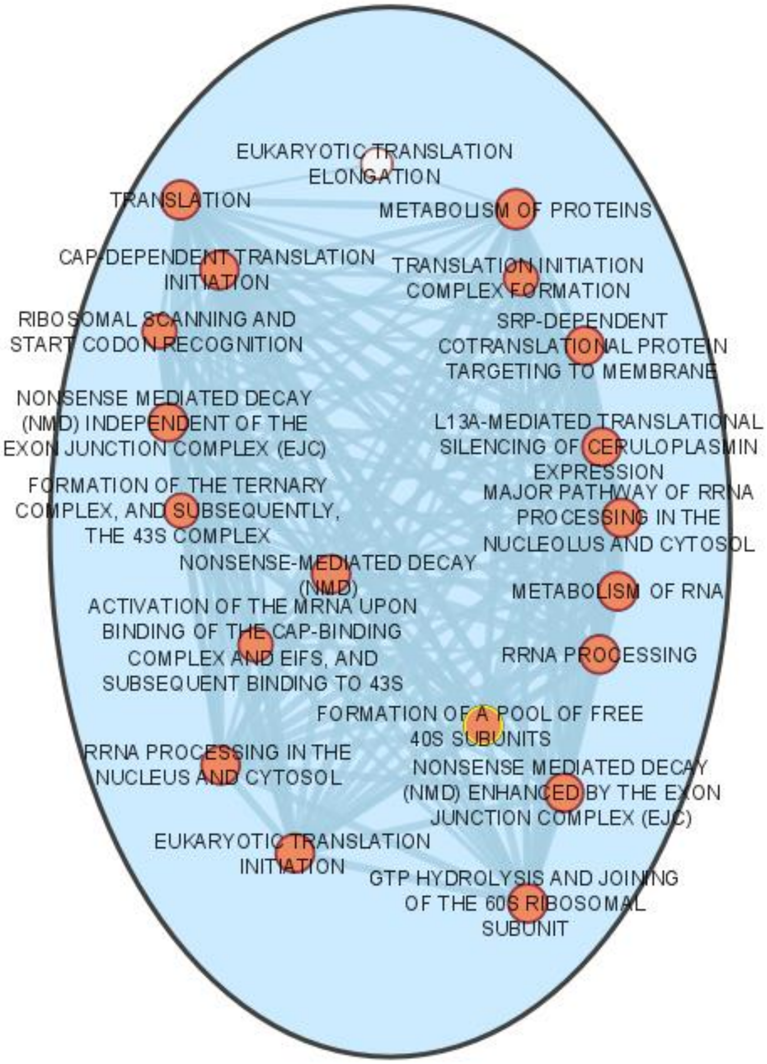
EUKARYOTIC TRANSLATION ELONGATION
TRANSLATION
METABOLISM OF PROTEINS
CAP-DEPENDENT TRANSLATION INITIATION
TRANSLATION INITIATION COMPLEX FORMATION
RIBOSOMAL SCANNING AND START CODON RECOGNITION
SRP-DEPENDENT COTRANSLATIONAL PROTEIN TARGETING TO MEMBRANE
NONSENSE MEDIATED DECAY (NMD) INDEPENDENT OF THE EXON JUNCTION COMPLEX (EJC)
L13A-MEDIATED TRANSLATIONAL SILENCING OF CERULOPLASMIN EXPRESSION
FORMATION OF THE TERNARY COMPLEX, AND SUBSEQUENTLY, THE 43S COMPLEX
MAJOR PATHWAY OF RRNA PROCESSING IN THE NUCLEOLUS AND CYTOSOL
NONSENSE-MEDIATED DECAY (NMD)
METABOLISM OF RNA
ACTIVATION OF THE MRNA UPON BINDING OF THE CAP-BINDING COMPLEX AND EIFS, AND SUBSEQUENT BINDING TO 43S
RRNA PROCESSING
FORMATION OF A POOL OF FREE 40S SUBUNITS
RRNA PROCESSING IN THE NUCLEUS AND CYTOSOL
NONSENSE MEDIATED DECAY (NMD) ENHANCED BY THE EXON JUNCTION COMPLEX (EJC)
EUKARYOTIC TRANSLATION INITIATION
GTP HYDROLYSIS AND JOINING OF THE 60S RIBOSOMAL SUBUNIT

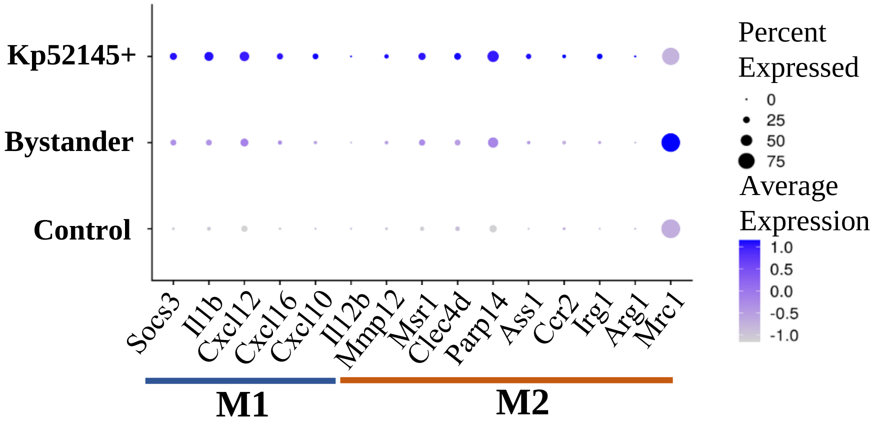
Kp52145+
Bystander
Control
Socs3
Il1b
Cxcl12
Cxcl16
Cxcl10
Il12b
Mmp12
Msr1
Clec4d
Parp14
Ass1
Ccr2
Irg1
Arg1
Mrc1
M1
M2
Percent Expressed
0
25
50
75
Average Expression
1.0
0.5
0.0
-0.5
-1.0

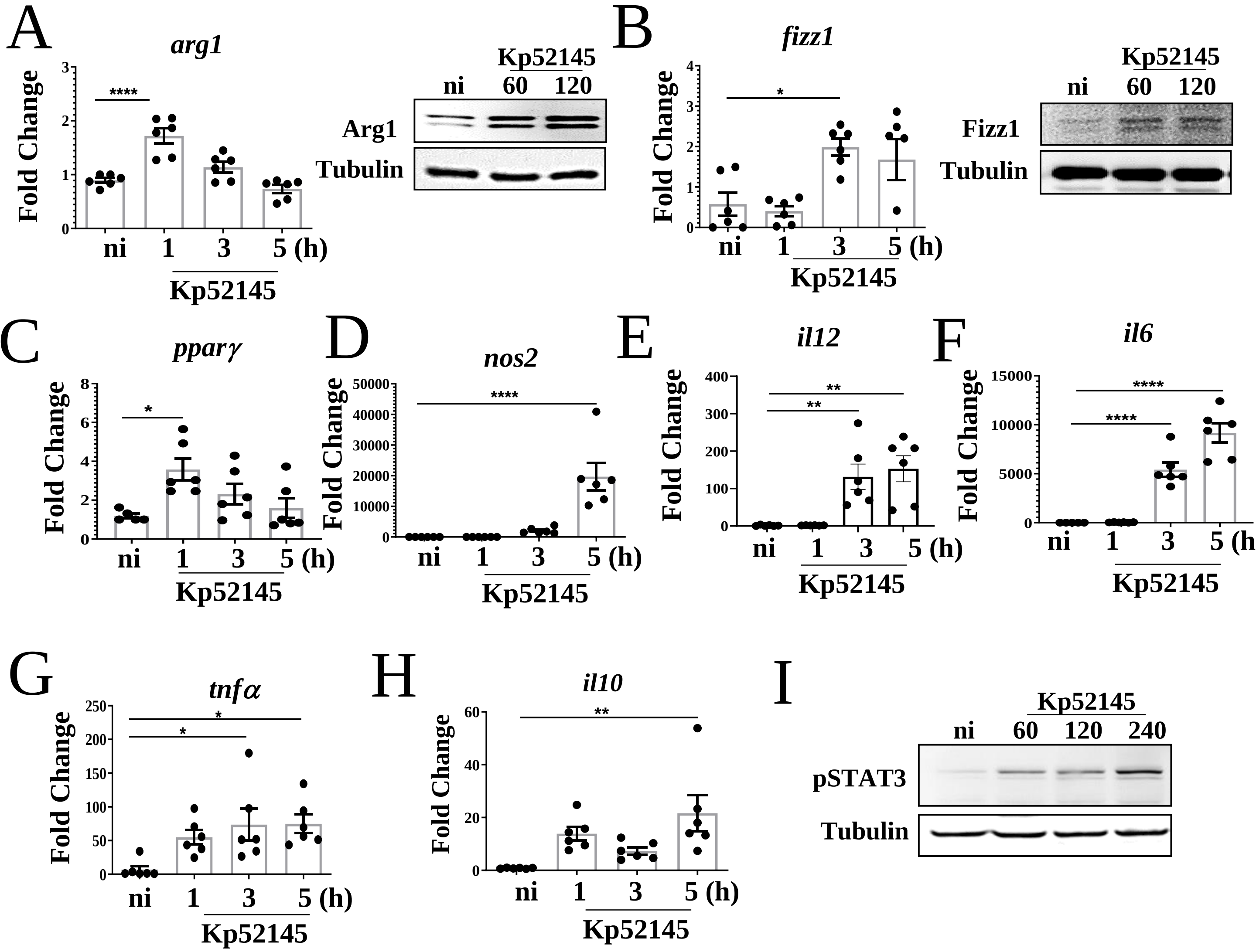
A
arg1
Fold Change
****
ni 1 3 5 (h)
Kp52145
Kp52145
ni 60 120
Arg1
Tubulin
B
fizz1
Fold Change
*
ni 1 3 5 (h)
Kp52145
Kp52145
ni 60 120
Fizz1
Tubulin
C
pparγ
Fold Change
*
ni 1 3 5 (h)
Kp52145
D
nos2
Fold Change
****
ni 1 3 5 (h)
Kp52145
E
il12
Fold Change
**
**
ni 1 3 5 (h)
Kp52145
F
il6
Fold Change
****
****
ni 1 3 5 (h
Kp52145
G
tnfα
Fold Change
*
*
ni 1 3 5 (h)
Kp52145
H
il10
Fold Change
**
ni 1 3 5 (h)
Kp52145
I
Kp52145
ni 60 120 240
pSTAT3
Tubulin

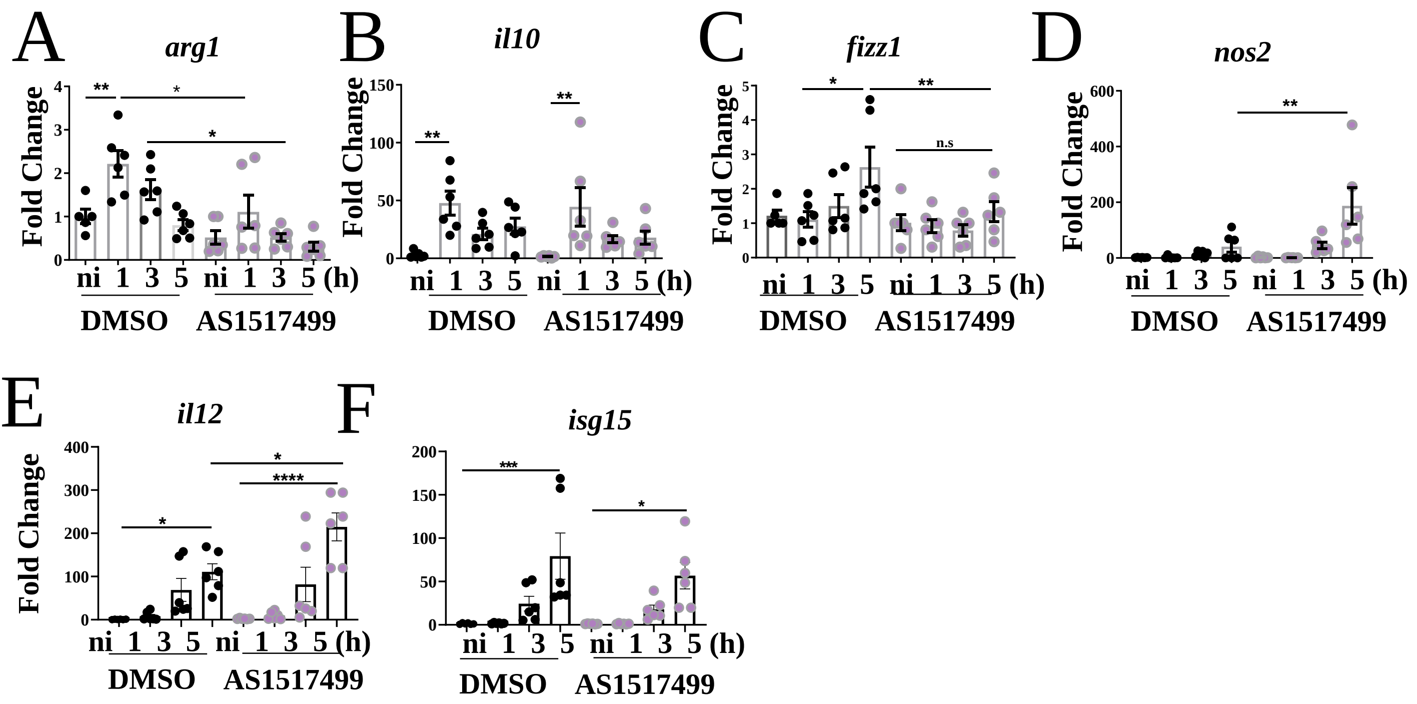
A
arg1
B
il10
C
fizz1
D
nos2
E
il12
F
isg15
Fold Change
ni 1 3 5 ni 1 3 5 (h)
DMSO
AS1517499
n.s

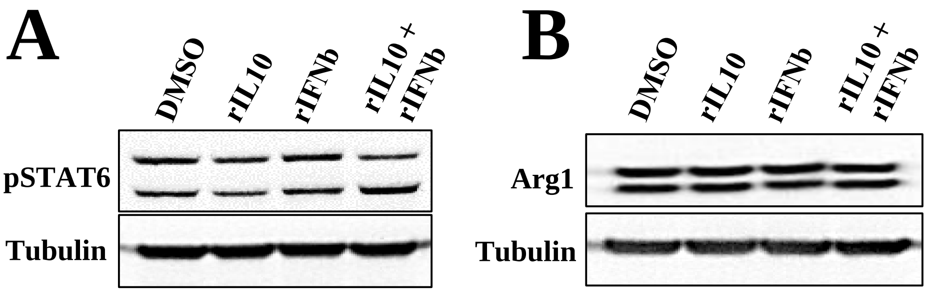
A
DMSO
rIL10
rIFNb
rIL10 + rIFNb
pSTAT6
Tubulin
B
DMSO
rIL10
rIFNb
rIL10 + rIFNb
Arg1
Tubulin

A

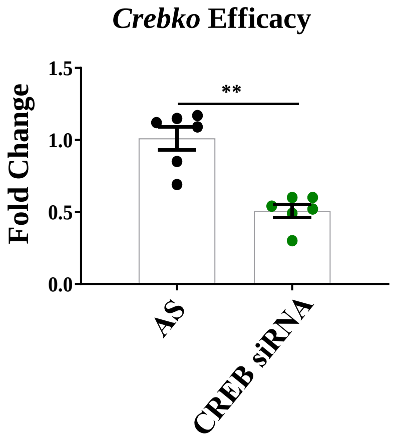

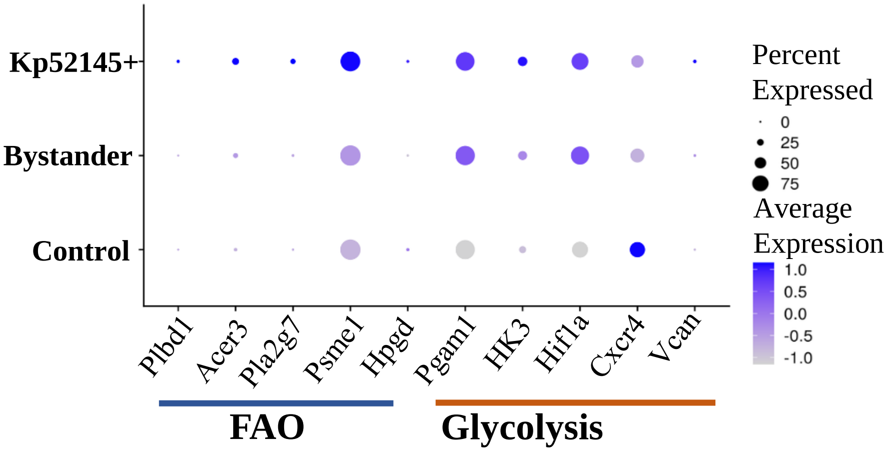

Kp52145+
Bystander
Control
Plbd1
Acer3
Pla2g7
Psme1
Hpgd
Pgam1
HK3
Hif1a
Cxcr4
Vcan
FAO
Glycolysis
Percent Expressed
0
25
50
75
Average Expression
1.0
0.5
0.0
-0.5
-1.0

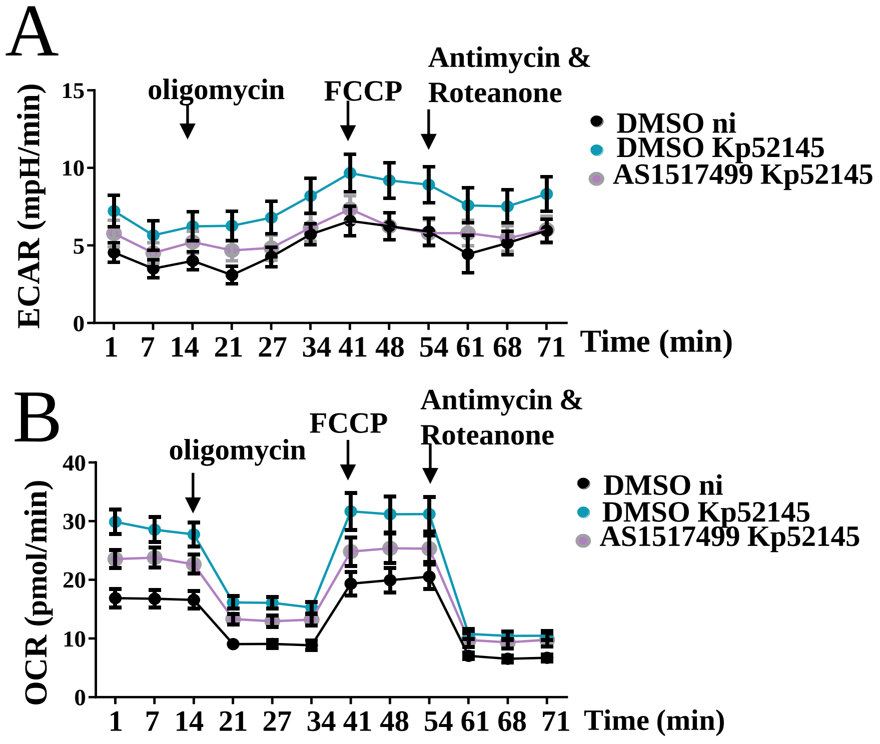
A
oligomycin
FCCP
Antimycin &
Roteanone
DMSO ni
DMSO Kp52145
AS1517499 Kp52145
ECAR (mpH/min)
15
10
5
0
1 7 14 21 27 34 41 48 54 61 68 71
Time (min)
B
oligomycin
FCCP
Antimycin &
Roteanone
DMSO ni
DMSO Kp52145
AS1517499 Kp52145
OCR (pmol/min)
40
30
20
10
0
1 7 14 21 27 34 41 48 54 61 68 71
Time (min)

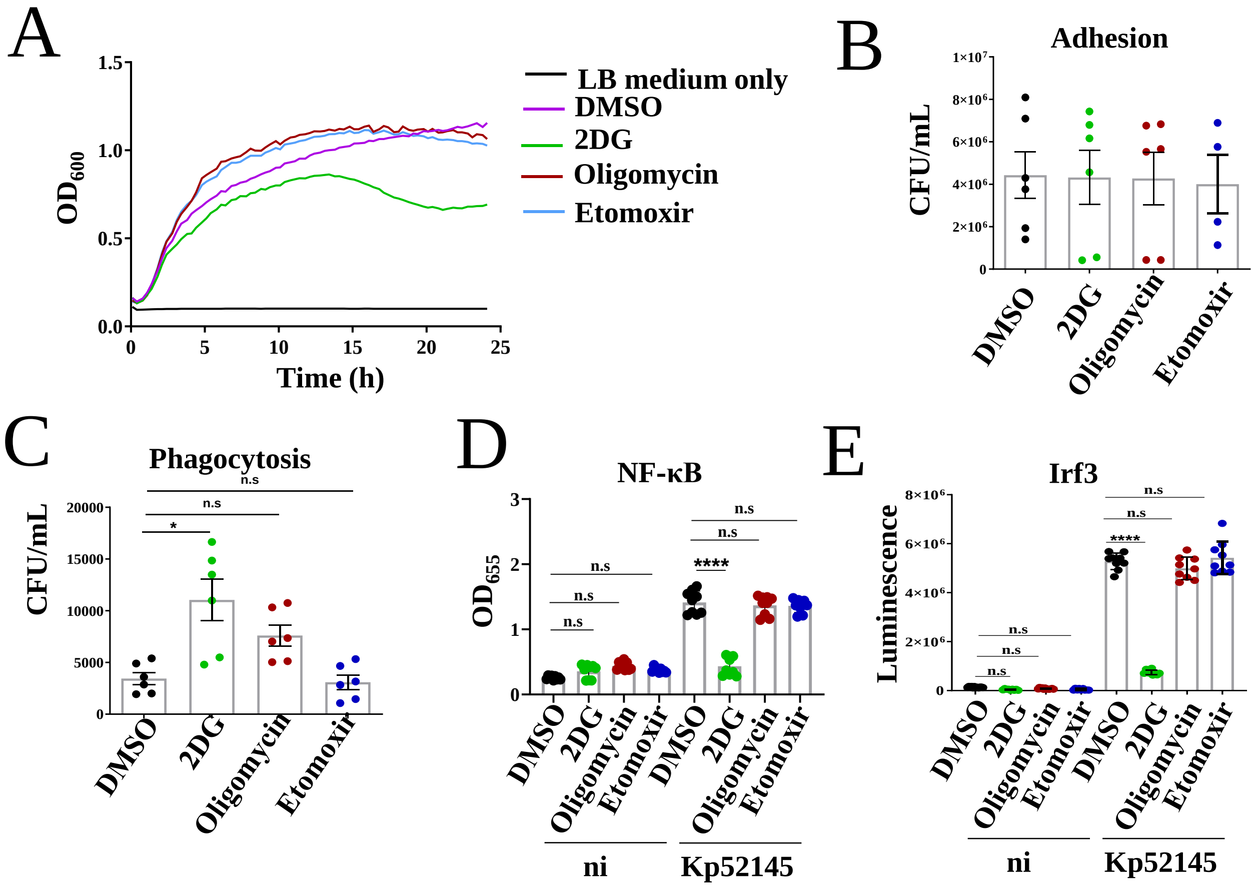
A
B
C
D
E
OD600
Time (h)
LB medium only
DMSO
2DG
Oligomycin
Etomoxir
Adhesion
CFU/mL
Phagocytosis
NF-κB
OD655
Irf3
Luminescence
n.s
*
****
ni
Kp52145
