## Supplementary table 3 for "In vivo single cell transcriptomics reveals *Klebsiella pneumoniae* rewiring of lung macrophages to promote infection"

**Supplementary Table 3**: **List of primers used for RT-qPCR in this study.**

| **Gene** | **Primer sequence** |
| --- | --- |
| **Mouse** |  |
| *arg1* | (Forward) 5 “CAG AAG AAT GGA AGA GTC AG 3”  (Reverse) 5” CAG ATA TGC AGG GAG TCA CC 3” |
| *il10* | (Forward) 5’ GGA CTT TAA GGG TTA CTT GGG TTG CC 3’  (Reverse) 5’ CAT GTA TGC TTC TAT GCA GTT GAT GA3’ |
| *nos2* | (Forward) 5’ TGG CTC GCT TTG CCA CGG ACG AGA CGG 3’  (Reverse) 5’ GGA GCT GCG ACA GCA GGA AGG CAG CGG G 3’ |
| *tnfα* | (Forward) 5’ TTC TGT CTA CTG AAC TTC GGG GTG ATC GGT CC 3’  (Reverse) 5’ GTA TGA GAT AGC AAA TCG GCT GAC GGT GTG GG 3’ |
| *isg15* | (Forward) 5’ GGG GCC ACA GCA ACA TCT AT 3’  (Reverse) 5’ CGC TGG GAC ACC TTC TTC TT 3’ |
| *mx-1* | (Forward) 5’ GAC TAC CAC TGA GAT GAC CCA GC 3’  (Reverse) 5’ ATT TCC TCC CCA AAT GTT TTC A 3’ |
| *pparg* | (Forward) 5’ TGT GGG GAT AAA GCA TCA GGC 3’  (Reverse) 5’ CCG GCA GTT AAG ATC ACA CCT AT 3’ |
| *il6* | (Forward) 5’ ATG GAT GCT ACC AAA CTG GAT 3’  (Reverse) 5’ TGA AGG ACT CTG GCT TTG TCT 3’ |
| *il12b* | (Forward) 5’ ACA GAG GAG GGG TGT AAC CA 3’  (Reverse) 5’ TAG CGA TCC TGA GCT TGC AC 3’ |
| *fizz1* | (Forward) 5’ GGT CCC AGT GCA TAT GGA TGA GAC CAT AGA 3’  (Reverse) 5’ CAC CTC TTC ACT CGA GGG ACA GTT GGC AGC 3’ |
| *klf4* | (Forward) 5” TGC CAG ACC AGA TGC AGT CAC 3”  (Reverse) 5” GTA GTG CCT GGT CAG TTC ATC 3” |
| *hprt* | (Forward) 5’ GAT CAG TCA ACG GGG GAC AT 3’  (Reverse) 5’ GGT CCT TTT CAC CAG CAA GC 3’ |
| **Human** |  |
| *arg1* | (Forward) 5“ TGG ACA GAC TAG GAA TTG GCA3”  (Reverse) 5” CCA GTC CGT CAA CAT CAA AAC T 3” |
| *il10* | (Forward) 5’ TCA CCT TCC AGT GTC TCG GA 3’  (Reverse) 5’ TAG CTG GGA TTA CAG GTG CG 3’ |
| *nos2* | (Forward) 5’ CAG CGG GAT GAC TTT CCA A 3’  (Reverse) 5’ AGG CAA GAT TTG GAC CTG CA 3’ |
| *tnfα* | (Forward) 5’ CTT TGG AGT GAT CGG CCC C 3’  (Reverse) 5’ GTT ATC TCT CAG CTC CAC GCC 3’ |
| *chi3l1* | (Forward) 5’ GAT AGC CTC CAA CAC CCA GA 3’  (Reverse) 5’ AAT TCG GCC TTC ATT TCC TT 3’ |
| *ido* | (Forward) 5” GCG CTG TTG GAA ATA GCT TC 3”  (Reverse) 5” CAG GAC GTC AAA GCA CTG AA 3” |
| *mrc1/cd206* | (Forward) 5’ GGC GGT GAC CTC ACA AGT AT 3’  (Reverse) 5’ ACG AAG CCA TTT GGT AAA CG 3’ |
| *il1rn* | (Forward) 5’ GGA ATC CAT GGA GGG AAG AT 3’  (Reverse) 5’ TCT CGC TCA GGT CAG TGA TG 3’ |
| *isg56* | (Forward) 5’ CAC CAT TGG CTG CTG TTT AGC TCC 3’  (Reverse) 5’ GGC AGC CGT TCT GCA GGG TTT T 3’ |
| *hprt* | (Forward) 5’ TTG CTT TCC TTG GTC AGG CA 3’  (Reverse) 5’ ATC CAA CAC TTC GTG GGG TC 3’ |
